# A Single-Cell Atlas of DNA Methylation in Autism Spectrum Disorder Reveals Distinct Regulatory and Aging Signatures

**DOI:** 10.1101/2025.06.17.660162

**Authors:** Katherine W. Eyring, Cuining Liu, Nasser Elhajjaoui, Kevin D. Abuhanna, Yi Zhang, Zachary von Behren, Eleazar Eskin, Daniel H. Geschwind, Chongyuan Luo

## Abstract

Autism spectrum disorder (ASD) is a common, genetically and clinically heterogeneous neurodevelopmental condition. Despite this diversity, studies of postmortem brain tissue have revealed convergent molecular changes across the cortex, including reduced synaptic function in subsets of excitatory and inhibitory neurons and increased glial reactivity. Whether these features are reflected in cell type–specific epigenetic signatures remains unknown. Here, we present the first single-cell analysis of DNA methylation (DNAm) coupled with transcriptomics in ASD. Using snmCT-seq, we profiled DNAm and gene expression from over 60,000 nuclei across 49 donors. We identified thousands of differentially methylated regions (DMRs) in ASD, enriched in promoters and regulatory elements active during both prenatal development and in adult cortex. ASD-related methylation changes were spatially localized but uncorrelated with gene expression, and were small in magnitude compared to robust age-associated effects. Age-DMRs were concentrated in excitatory neurons, enriched in known cognitive aging pathways, and revealed distinct roles for CG and non-CG methylation in the aging brain. Finally, we explored age-by-diagnosis interactions, identifying a reduction in inhibitory neuron abundance with age in ASD relative to controls, highlighting this area as a promising direction for future research.

**Highlights:**

- We generate a single cell multi-omic dataset, jointly profiling DNA methylation and gene expression in autistic and neurotypical donors
- We identify thousands of cell type informed differentially methylated regions (DMRs) in ASD, particularly in excitatory neurons from superficial cortical lamina and microglia
- ASD-DMRs are enriched in promoters and known regulatory regions, but not strongly tied to gene expression
- Age effects on DNA methylation are profound, cell type specific, and concentrated in excitatory neurons

## Introduction

DNA methylation (DNAm), or the addition of a methyl group to the 5th carbon of cytosine (mC), is a ubiquitous epigenetic mark known to vary across tissues [1–3], age [4–7] and cell-types [8–10], also reviewed in [11]. DNAm plays a critical role in cellular differentiation and the regulation of gene expression (reviewed in [12]). Epigenetic regulation generally [13–17], and DNAm specifically, have also been implicated in the etiology of neurodevelopmental disorders (NDDs) including autism spectrum disorder (ASD). Genetic variation in “writers” and “readers” of DNAm causes profound neurological syndromes. Variants in de novo methyltransferases (DNMT), which establish and maintain DNAm [18–21], cause autosomal dominant cerebellar ataxia, deafness and narcolepsy (*DNMT1,* [22]) and Tatton-Brown-Rahman syndrome (TBRS, *DNMT3A*, [23, 24]), while Rett Syndrome is caused by variation in methyl cytosine binding protein 2 (*MECP2*, [25]). Autistic features are often observed in Rett syndrome and TBRS, and *DNMT3A* itself is a high-confidence ASD risk gene [17, 26–29]. Loss of *DNMT3A* or the introduction of patient mutations into mice impairs neuronal maturation and elicits behavioral phenotypes in mice [30, 31].

Genomic context informs the function of mC. Most CpG (cytosine followed by guanine) sites in the adult human genome are methylated (>70%), while regulatory regions, like promoters and enhancers, are distinctly hypomethylated in a cell-type-specific fashion [8, 10, 32]. In addition, methylation outside of CpG sites (hereafter referred to as mCH, where H = A, T, C) accumulates early in life, specifically in neurons [1, 6, 33], implicating the developmental stage and cell types most relevant to NDDs.

A convergent transcriptomic [34–39] and epigenomic profile [40–43] is emerging from the study of postmortem cortical tissue from autistic donors. Generally, these studies of bulk tissue describe down-regulation of synapse-related transcripts, up-regulation of transcripts related to immune signaling and loss of regional patterning. These observations have been refined by recent single-cell studies [39, 44] that identified compositional and cell-type specific changes in gene expression that, together, contribute to this signature. ASD molecular pathology has also been observed in regulatory miRNA and long-non-coding RNA [37, 43], genomic enhancers (implicated by H3K27Ac binding, [41]) and CpG methylation detected at CpG sites covered by the Infinium Human Methylation 450K BeadChip [42, 45, 46] and in bulk whole-genome bisulfite sequencing [47]. DNAm can be incredibly cell-type specific [9, 10]. However, it is unknown whether previously described changes in bulk tissue reflect compositional changes [48], bona fide changes in DNA methylation in specific cell types, or a combination of both. In this study, we expand upon existing knowledge by profiling DNA methylation patterns in ASD with cell-type resolution. We interpret our results by integrating methylation and gene expression data jointly generated from individual cells using the recently developed snmCTseq method [49] and in the context of published data on molecular phenotypes in autism. In addition, we leverage this unique dataset to define age- and cell type-specific methylomic variation. To our knowledge, this data represents the first population scale single-cell study to investigate ASD-associated changes in the DNA methylome.

## Results

### Single-cell multi-omic profiling of ASD and neurotypical frontal cortex

To investigate cell-type specific methylation patterns in autism, we first isolated nuclei from the prefrontal cortex (Brodmann area 9) of 26 idiopathic autistic donors and 23 non-autistic controls (**Fig. 1a**). Donors ranged in age, from 2 to 60 years, and age did not differ significantly between groups (**Fig. 1b,** p = 0.65, *Wilcoxon rank sum test*). Due to the strong male bias in ASD prevalence, we only profiled male donors. Isolated nuclei were sorted, 1 per well, into 384-well plates based on the presence or absence of the neuronal marker NeuN (*RBFOX3*), maintaining an approximate 2:1 neuron to non-neuron ratio per donor. We then jointly assayed gene expression and DNA methylation from individual cells using snmCT-seq [49] for 236 plates.

**Figure 1.**
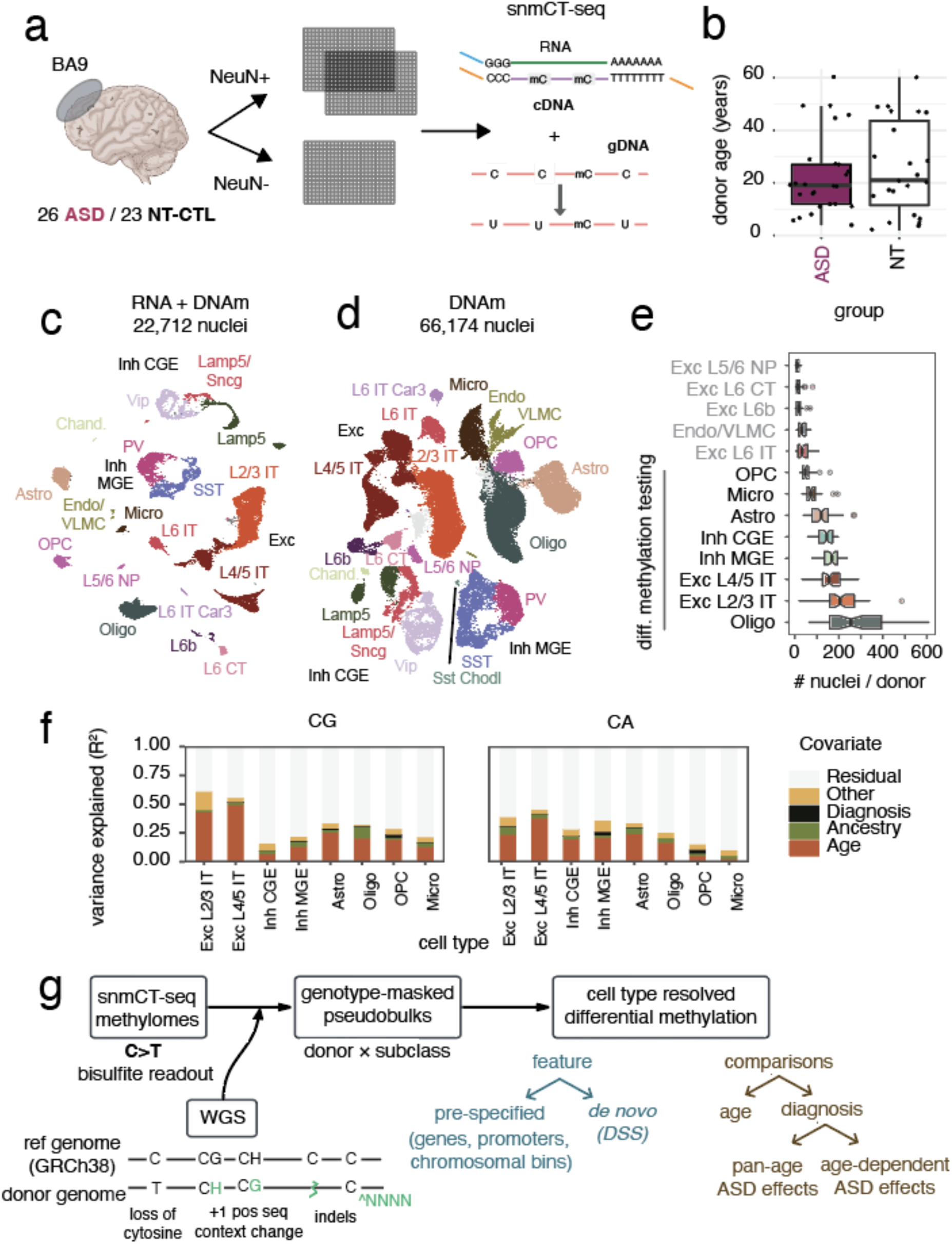
*Dataset overview*. (a) schematic of data processing in this study. (b) age distribution of donors in this dataset, separated and colored by diagnosis. UMAP representations of the dataset integrating RNA and methylome data (c) or using genome- wide DNA methylation alone (d). Cell type labeling and coloring are consistent in **c** - **e**. (e) distributions of nuclei per donor per cell type. (f) variance explained by biological variables across cell types and genomic contexts. (g) graphical summary of main analyses in paper.

Biochemical processing (reverse transcription, cDNA amplification and bisulfite conversion) was performed in each well before libraries were tagged with well- and plate-specific indices and pooled for sequencing. A detailed protocol describing the experimental steps in snmCT-seq is provided through protocol.io (https://www.protocols.io/view/snmcat-v2-x54v9jby1g3e/v2; the *in vitro* GpC methyltransferase treatment step in the snmCAT-seq protocol omitted in this study).

We distinguished between reads derived from genomic DNA and RNA based on their methylation levels in the non-CG (CH) context as previously described ([49], RNA reads were engineered to have high methylation levels, *Methods*) and pre-processed data from each modality separately before integration. Following methylome quality control checks (**Fig. S1a-e**), we recovered data from 66,174 nuclei (median: 1,347 per donor). Nuclei were sequenced to an average depth of 1.5 million reads, 83% of which derived from genomic DNA. We observed anticipated global methylation levels ≥0.75 in the CpG context, ≤0.12 in the CpH-context, and ≤0.03 in the CpC (**Fig. S1c**-**e**). We recovered high-quality RNA libraries from 27,515 nuclei following QC filtering (**Fig. S1f-h**, *Methods*). The yield of nuclei per donor did not significantly differ between groups (**Fig. S1i**, p = 0.82, *Welch’s t-test*). We leveraged information from both modalities to define and annotate the clusters referenced in the remainder of this study.

Unsupervised clusters were defined after integrating genome-wide 100kb bin methylation (in CG and non-CG contexts) with gene expression (**Fig. 1c, d**; *Methods*). Donors were well-represented in every cluster (**Fig. S2a**) and clusters were readily annotated on the basis of canonical marker gene expression or gene body hypomethylation (**Fig. S3**) or projection into reference datasets (**Fig. S2b**). Annotations were generally concordant across RNA and methylation modalities (**Fig. S2c, d**), aligned with our NeuN staining and the known accumulation of mCH in neurons (**Fig. S2e, f**), and reflected anticipated cortical cell types.

We identified 28 clusters in our unsupervised integration of RNA and DNA methylation data that were annotated and aggregated into subclasses (hereafter referred to as cell types). Five clusters correspond to non-neuronal cell types: astrocytes (Astro), oligodendrocytes (Oligo), oligodendrocyte precursor cells (OPC), microglia (Micro) and 1 small cluster containing both endothelial (Endo) and vascular associated cells / pericytes (or vascular lepotomeningeal cells, VLMCs). We observed ten inhibitory neuron clusters, which could be grouped into those originating from the medial and caudal ganglionic eminences in development (MGE- and CGE-, respectively) and included MGE-derived parvalbumin (*PV*) and somatostatin (*SST*) expressing cells and CGE-derived *VIP* (vasoactive intestinal peptide), *SNCG* and *LAMP5* subtypes. Twelve clusters corresponded to different subtypes of excitatory neurons. Two clusters were annotated as superficial layer excitatory intra-telencephalic neurons (Exc L2/3 IT) and constituted the largest neuronal population (**Fig. 1e**). 4 clusters were aggregated into the L4/5 IT subclass, reflecting a gradient of laminar specific expression patterns (**Fig. S3**). A number of smaller, yet distinct, clusters were annotated as different deep-layer excitatory subtypes: L6 IT, L6 corticothalamic projecting (CT), L6b, L5/6 near-projecting (NP). To analyze methylomes under this unified annotation schema, including cells for which high-quality gene expression was not available, we trained a weighted nearest neighbor classifier to assign cell type annotations based on the methylome alone (*Methods;* hold-out accuracy ≥97.4%).

### Transcriptomic signatures of ASD

We first assessed transcriptomic differences between our control and autistic donors. Between groups, donors did not differ in sample quality metrics (**Fig. S4**) or cell-type abundance (**Fig. S5**). We assessed cell-type abundance using two different approaches: one, based on our subclass and cluster labels and the other using a label-free nearest neighbors-based method [50]. We did not observe any significant changes to subclass or cluster abundance after FDR correction and no consistent changes across methods.

We next assessed differential gene expression (DGE) between NT-CTL and ASD samples in pseudobulk expression (generated by summing gene counts from the same cell type and donor) using DESeq2 (**Fig. S6**). As has been reported previously [39, 44], we detected the largest number of differentially expressed genes (DEGs) in excitatory L2/3 IT neurons (608 DEGs, padj < 0.05). Of note, two clusters (**Fig. S6a**) comprised the L2/3 IT subclass and differed in their expression of genes known to distinguish L2 and L3 neurons, including *PDGFD* and *LINC01500* (L2) and *COL5A2* (L3; **Fig. S6b,** [51]). When we performed DGE analysis at the cluster level, DEGs were concentrated specifically in the putative L2 cluster (**Fig. S6c, d**). The increased number of DEGs in putative L2 neurons compared to L3 neurons was robust to down-sampling analysis (**Fig. S6e**, *Methods*). We observe a significant correlation between ASD effects in our data and those published in a previous single cell study of gene expression in ASD (Pearson’s correlation coefficient = 0.35, p < 2.2E-16, of diagnosis effect size within Exc L2/3 neurons; **Fig. S6f,** [44]), supporting the robustness of our results. Differentially expressed genes were enriched in 129 functional pathways (gene set enrichment analysis, padj < 0.1), 81% of which were observed in Exc L2/3 neurons. Many of the top pathways suggested upregulated transcripts converged on the cellular response to stress, including the unfolded protein response pathway (**Fig. S6g**), as has been observed previously [39]. Many of the top DEGs, driving gene set enrichment, are known to be expressed in the brain and have been implicated in neurodevelopmental syndromes previously (for example, *YWHAG* [52]; **Fig. S6h**).

### Strategies to identify ASD-associated DNAm signatures

Genetic variants such as single nucleotide variants (SNVs) in the genome can have direct effects on the estimation of DNA methylation in whole-genome bisulfite data when they overlap with a cytosine (position 0) or the base downstream of a cytosine (position 1). For example, a C>T single-nucleotide variant may be interpreted as a complete loss of cytosine methylation. In addition, methylation strongly varies by +1 sequence context (**SFig. 1c-e)**. To control for high impact donor genetic variation on the methylome profiling, we performed whole-genome sequencing (40.9X ± 10.3X; mean ± sd coverage) and genotyping to identify and mask high impact variants in our methylome data (“genotype masking”, *Methods*). In this “genotype masked” data, we then performed differential methylation analysis in pseudobulk data (aggregating by donor and cell type).

Human brain development is associated with DNA methylation dynamics at different scales, ranging from individual differentially methylated cytosines (base-resolution) to mega-base scale methylation valleys [6, 7, 53]. We investigated cell-type specific differential methylation in ASD using two complementary methods, each tuned to detect different genomic features. First, we called *de novo* differentially methylated regions (DMRs) using the DSS [54–56], a sensitive two-group DMR caller that exploits correlation between adjacent CG-context cytosines. In the second approach, we analyzed pre-defined features, including genes, promoters and 500bp bins tiling the genome. While the former provides single cytosine resolution, the latter offers higher coverage due to the region binning (**SFig. 7a, b**), more flexible covariate adjustment structures, and testing of non-CG methylation levels. Henceforth, *de novo* regions identified by DSS will be referred to as DMRs (differentially methylated regions), whereas pre-defined regions will be referred to as differentially methylated bins, promoters or genes (DMBs, DMPs, DMGs, respectively). We focused on the eight most abundant subclasses in our methylation analyses (**Fig. 1e**), further aggregating some subclasses (for example, combining different MGE derived interneurons) to increase our coverage (achieving up to 10X pseudobulk coverage in our most abundant cell types).

We began by characterizing the largest sources of overall methylome variance in our dataset. As expected, cell type explained 80-99% of the global variance across combined pseudobulk methylomes (**Fig S7c, d**). Within each cell type, age explained more variance than any other known donor features (**Fig 1f, S7e**). We thus examined differential methylation by diagnosis and age, in addition to an exploratory analysis of age-dependent ASD-effects, as schematized in **Fig 1g**.

### ASD-associated DMRs overlap with cell-type-specific regulatory elements

CG methylation at promoters and enhancers is a reliable indicator of regulatory activity; the depletion and enrichment of CG methylation is correlated with active and inactive states, respectively [32]. Consistent with past reports [42, 47], we did not find significant differences in global methylation levels between diagnostic groups (p > 0.15, beta regression, Wald test). We began by investigating pre-defined features (promoters, genes) whose methylation levels varied by diagnosis, irrespective of donor age (pan-age), but identified few (244 genes or promoters total, 1-27 per cell type; *Methods*). Testing in pre-defined regions may be unable to detect differential methylation that, for example, overlaps only one portion of a gene or a short region.

We thus also identified pan-age *de novo* differentially (CG) methylated regions (DMRs) associated with ASD in a genome-wide and cell-type informed fashion, using a rigorous computational strategy including permutation of the diagnosis label (see *Methods;* **Fig 2a**). Across 8 cell types tested, we identified 31,165 DMRs, which were more often hypo-methylated in ASD than hyper-methylated **(Fig. 2b**). DMRs ranged in size (median ± standard deviation = 132 ± 185 bp, examples in **SFig. 8, 9**) and were very cell-type specific (<2% of DMRs overlap other DMRs across cell types). The number of DMRs detected per cell type was not a function of the number of cells per cluster, which ranged from (2,548 to 13,887, **Fig. 2b**). In a hierarchical clustering of ASD-DMRs, most DMRs cluster by cell type and then by age (in excitatory neurons), consistent with methylation patterns associated with cell type and age effects being much larger than those that vary with diagnosis (**Fig. 1f**, **S9a**).

**Figure 2.**
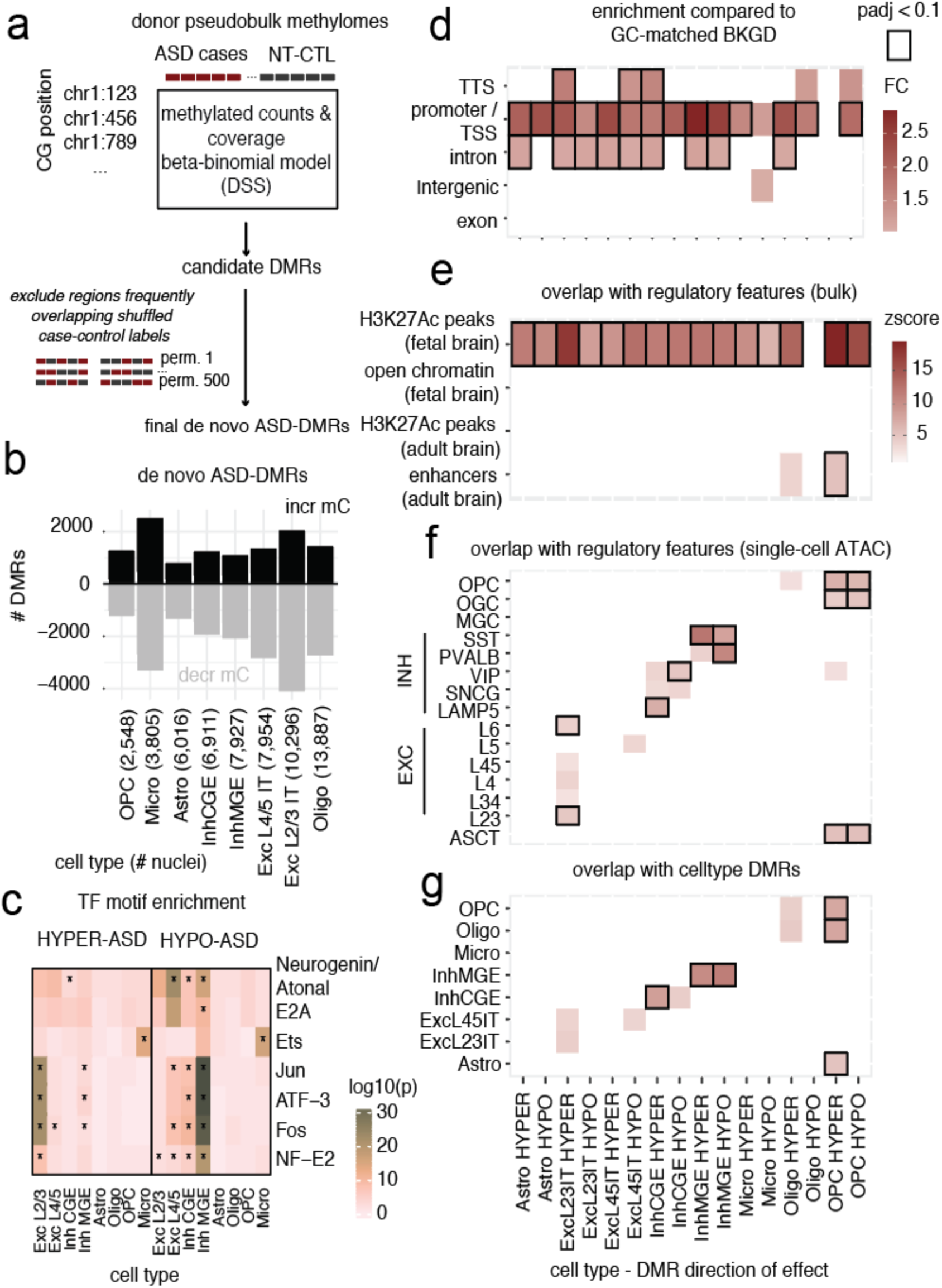
*ASD DMRs.* (a) Schematic outlining our DMR calling approach. (b) Number of ASD-associated DMRs per cell type. Direction of effect indicated by color and direction of the bar. (c) Transcription factor (TF) motif enrichment in ASD-DMRs. (d) Enrichment of ASD-DMRs with genomic features; fold change depicted. (e) Enrichment of ASD-DMRs with regulatory regions identified in studies of bulk prenatal and adult brain tissue (e), cell type specific regulatory elements in adult cortex (f), and cell type defining DMRs (g). In d-g, only significant enrichment (p<0.01) is shown, with emboldened tiles if padj < 0.1. In *c*, motifs with -log10(pval) > 15 are shown.

The potential value of using a single-cell approach to identify DMRs associated with ASD is underscored by the cell type specificity of DNA methylation (**Fig. S7c**; [9, 10]) and the increased number of DMRs identified here compared to previous efforts using analysis of bulk DNA, which identified hundreds of DMRs in the prefrontal cortex of autistic donors [42, 47].

We first assessed whether specific transcription factor (TF) binding motifs were overrepresented in our ASD-DMRs using *HOMER* [57]. TGA*TCA motifs, characteristic of Fos/AP1-family TFs (which include *Fos*, *ATF3* and *Jun*), were notably enriched in two ASD-DMR sets–those hypomethylated in MGE-derived interneurons and regions hypermethylated in Exc L2/3 IT neurons (**Fig. 2c**). This enrichment is particularly interesting given that Fos-family TF binding is known to be sensitive to motif methylation [58] and is altered in ASD based on TF footprinting analysis from bulk [59] and single-cell data [39].

Consistent with the known distribution of genomic features, most DMRs fall into intergenic and intronic regions; though, ASD-DMRs overlap promoters twice as often as expected by chance (5.3% promoters in ASD-DMR set compared to 2.2% promoters in the genomic background; median log2FC across cell types and directions = 1.15, significant in all but two DMR sets (OPC_HYPER, MICRO_HYPO), Bonferroni-corrected p < 0.004). This promoter enrichment was evident relative to a background of all tested regions (“candidate DMR regions” in *Methods*) and in a smaller background set matching the GC-content of our ASD-DMRs (**Fig. 2d**, *Methods*) and is supported by chromHMM genomic annotation analysis (**Fig. S10a**, [60, 61]). To test whether the observed promoter enrichment of ASD-DMRs might reflect a bias inherent to *DSS’* use of the correlation between nearby CG sites, we contrasted the genomic feature enrichment of ASD- DMRs to alternative sets of DMRs called between cell types or age groups using the same dataset and *de novo* DMR calling methodology. Promoter enrichment was broadly observed in all de novo DMR sets, but it was most pronounced in the ASD-control comparison set (**Fig. S10b**). Our cell type DMRs significantly overlapped those previously identified in [62] (**Fig. S10d**) using different donors and methods, reinforcing the validity of our data and approach.

We next asked whether our ASD-DMRs overlapped known regulatory regions in the brain by assessing enrichment in four sets of previously published regulatory regions : H3K27Ac peaks called in prenatal brain [63], accessible chromatin regions in prenatal brain [64], H3K27Ac peaks and high-confidence enhancers in adult cortex [63] (**Fig. 2e**). We observed significant enrichment of ASD-DMRs with H3K27Ac-bound regions in the prenatal brain (Bonferroni-corrected p < 0.02, excluding OLIGO_HYPO; **Fig. 2e**). This enrichment was not specific to ASD- DMRs, however. DMRs that distinguish cell types were also strongly enriched in prenatal enhancers and open chromatin (**Fig. S10c**), consistent with methylation patterns generally reflecting activity in the developing brain. Hyper-methylated ASD-DMRs in OPCs and Oligos also overlapped adult brain enhancers (**Fig. 2e**).

We then extended our analysis to cell-type informed, candidate cis-regulatory elements (cCREs) identified in an independent single cell ATAC-seq analysis of the adult brain [65] (**Fig. 2f**). We found that ASD-DMRs significantly overlapped cCREs in their corresponding cell types (36% of ASD-DMRs (11,271 / 31,165) were in cCREs). Hyper-methylated ASD-DMRs in Exc L2/3 IT neurons significantly overlapped cCREs observed in Exc IT neurons, while DMRs identified in MGE- and CGE- derived interneurons and OPCs also significantly overlapped cell type aligned cCREs (**Fig. 2f**). The cCREs overlapping ASD-DMRs were relatively cell type specific (median # of brain cell types in which a given cCRE observed = 9, range = 1 – 102) and 3,847 have been previously annotated to genes [65]. ASD-DMRs also significantly overlapped regions that are uniquely hypomethylated in their corresponding cell types [62], as may be expected from cell type-specific regulatory regions (**Fig. 2g**).

To develop our understanding of the functional implications of differential methylation in ASD, we next sought to connect our DMRs to genes. We annotated DMRs based on their overlap with genes and promoters, as well as using cCRE-based annotations ([65], *Methods*). The overlap of ASD-DMRs with prenatal brain enhancers (**Fig. 2e**), prompted us to first consider whether they may reflect altered epigenetic regulation at an early stage of development and implicate prenatally expressed genes. We did not, however, observe higher expression of genes implicated by ASD-DMRs in prenatal samples compared to postnatal samples in the BrainSpan database ([66], **Fig. S11a**).

Next, we explored the relationships between our ASD-DMRs and extensive published knowledge around the molecular pathology of ASD. First we assessed whether ASD-DMRs overlap putative enhancers that are dysregulated in ASD [41] and observed strong overlap between DMRs hypermethylated in microglia, which were associated with genes involved in morphological and immune signaling (**Fig. S11b**), and overlapped down-regulated regions identified in H3K27Ac profiling of ASD cortex (**Fig. S11c**). Next, we assessed whether our cell-type resolved ASD-DMRs overlapped differentially methylated sites and regions identified in previous (bulk) analysis from ASD cortical tissue [42, 47]. We found limited overlap between different bulk ASD methylome analyses (**Fig. S11d**). While the absolute overlap between published ASD-DMRs and those identified in this study was small (<3% of previously reported ASD-DMRs were detected in our dataset), it was more than expected by chance (**Fig. S11e**). Published ASD-DMRs also overlapped cell type DMRs (**Fig. S11f**).

We then considered how the ASD methylome signature that we’ve identified relates to known transcriptomic changes in ASD. Genes implicated by DMRs did not significantly overlap those differentially expressed, in the corresponding cell-types, in data from this study or a larger dataset [39]. In a complementary analysis, we returned to effect sizes determined in our pre-specified analysis of genes and promoters, but found no correlation in diagnosis effect sizes between methylation and gene expression (all p > 0.05, Pearson’s correlation). In sum, cell-type specific ASD-DMRs overlap likely regulatory regions marked by accessible chromatin but are not associated with diagnosis-associated transcriptomic changes.

### Age-associated remodeling of DNA methylome – de novo DMR discovery

A particular strength of this dataset is the broad distribution of ages represented by our donors (2 - 60 years old; **Fig 1b**), which allowed assessment of cell type-resolved methylation signatures of aging across the lifespan. Age explained the largest proportion of methylome variation within almost all cell types, in genic- and intergenic- features, and in CA- and CG-cytosine contexts (**Fig. 1f; Fig. S7e**). Global methylation and its accumulation over age varied by cell type; mCG/CG increased with age in excitatory neurons (beta regression Wald P < 5x10-4 in L2/3 IT and L4/5 IT, P = 0.05 - 0.60 in other cell types, **Fig. 3a**). Cell type composition also varied with age (**Fig. S12**). The abundance of OPCs decreased, while the relative abundance of oligodendrocytes increased with age, as has been observed in mice [67] and human [68]. The abundance of inhibitory neurons decreased with age, as has been reported previously [69], with a pronounced decrease in the abundance of CGE-derived neurons specifically in autistic donors relative to controls (**Fig. S12,** nominal p (age*diagnosis interaction term) = 0.02).

**Figure 3.**
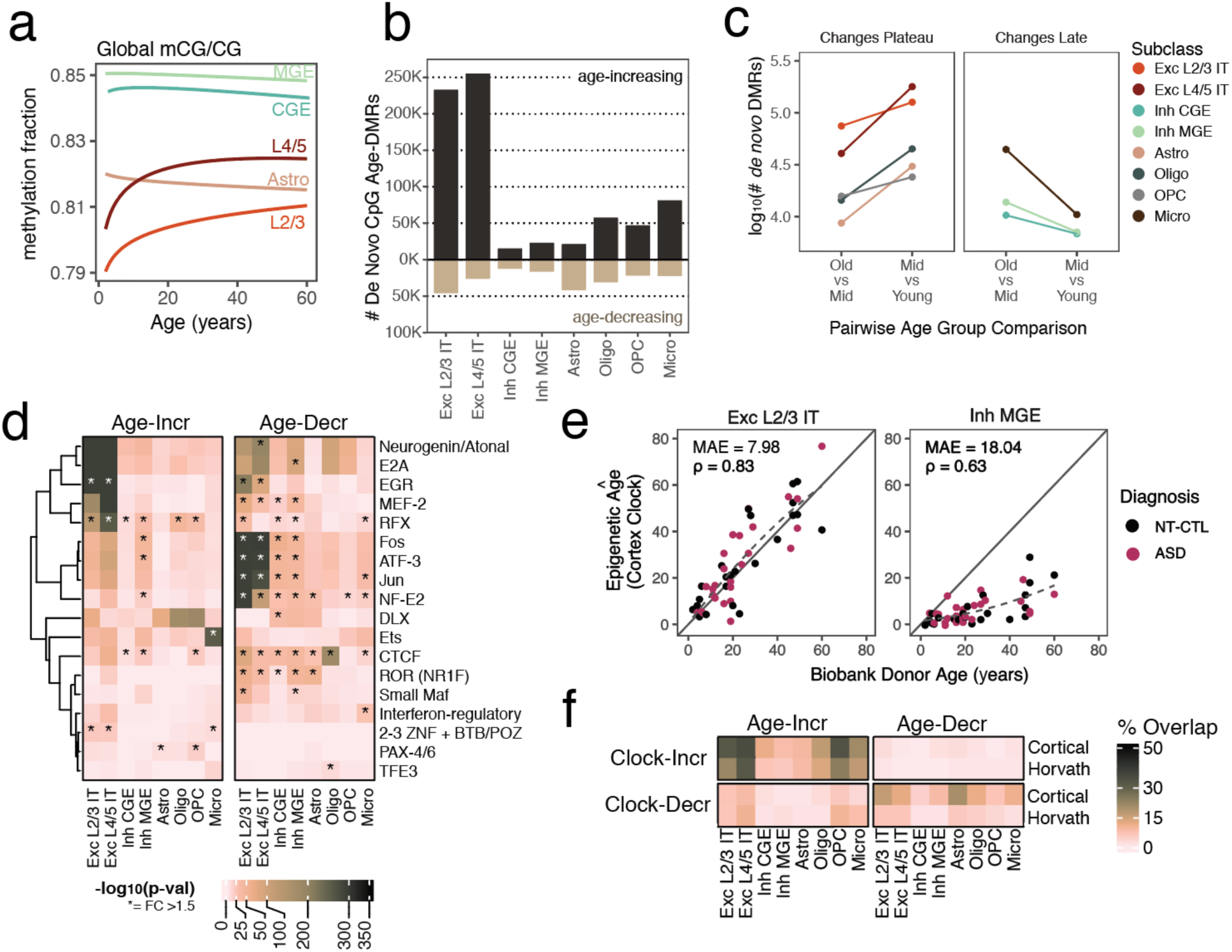
De novo age-DMRs. (a) Global CG-methylation fraction across five subclasses. (b) Number and directionality of unique *de novo* age-DMRs detected from pairwise comparisons of three age strata. (c) Total number of age-DMRs detected, by pairwise comparison and subclass. (d) Subset of significant ranscription factor motif enrichments in age-DMRs. Each motif included has at least one contrast with FDR < 0.05, fold-change ≥ 1.5, and >100 sequence motifs overlapped in the set. (e) Example application of the cortex age clock to pseudobulk data, with mean absolute error (MAE) and Spearman correlation ρ used to measure the similarity of the predicted epigenetic age to the biobank donor record. (f) Fractional overlap between array-based epigenetic clock probes and age-DMRs, split by predicted direction of methylation change with aging.

We went on to identify just over 1.4 million *de novo* CpG DMRs associated with age (age-DMRs, *Methods*) in a series of pairwise comparisons between young donors (2 - 16 years), young adults (17 – 30 years) and adult donors (40 - 60 years). Age-DMRs generally accumulated methylation with age and spanned 625,306 non-overlapping regions (10.3% of genome; **Fig. 3b**). Age effects were most pronounced (in variance explained (**Fig 1f, S7e**) and sheer number of DMRs (**Fig. 3b**)) in excitatory neurons, followed by non-neuronal cells and then inhibitory neurons (**Fig. 3b**). The number of age-associated features detected per cell type was uncorrelated to cluster size (Spearman *ρ*≤0.18), mirrored in number of age-DEGs in our paired transcriptome data and were consistent with previous reports [69] (**Fig. S13a-c**). In most cell types, an order of magnitude more age-DMRs were detected when comparing young to young adult donors than when comparing young adult to adult donors, suggesting significant remodeling before adulthood; in contrast, inhibitory neurons and microglia had more mCpG changes later in life (**Fig. 3c**).

Transcription factor motif enrichment was abundant in age-DMRs (**Fig. 3d**), and was often shared among multiple subclasses. We noted a striking loss of methylation with age in AP-1 transcription factor complex (*FOS*, *JUN*) motifs and an accumulation of methylation in motifs associated with TFs involved in neurodevelopment, including Neurogenin/Atonal TFs (e.g., *NEUROD1*, *OLIG1*, *ATOH1*, *BHLHE22*) and the *RFX* TF family [70]. Almost all cell types exhibited significant hypomethylation at CCCTC-binding factor (*CTCF*) binding motifs with advancing age, which complements mounting evidence of changes to 3D chromatin conformation during neuronal maturation, aging and Alzheimer’s disease [62, 68, 71, 72]. We additionally reproduced cell type specific aging signatures previously reported, including age-accumulation of methylation over EGR/EGF motifs in excitatory neurons [69] and accumulation in Ets-family motifs (including PU.1 and ELK) in microglia [68].

Lastly, we assessed the overlap between our age-DMRs and the well-established “epigenetic clocks”, which use array-based measurements of cytosine methylation to predict chronological age. Although the clocks have shown high accuracy across multiple bulk tissues and cohorts, there has been limited assessment on brain cell types from single-nucleus data (one other dataset [68]). We considered the canonical “Horvath” clock [5], trained on pan-tissue data, and a “cortical clock” trained specifically on human prefrontal cortex [73]. Clock performance (as quantified by mean absolute error and Spearman correlation between predicted and biological age) was highly cell-type dependent, with high performance of Exc L2/3 in both clocks and poor performance of MGE-derived inhibitory neurons and microglia in both clocks (**Fig. 3e**, **S14**). This differential performance by cell type, appeared more attributable to true biological differences in epigenetic aging (age-DMRs overlapped 0% to 35% of probes by cell type; **Fig. 3f**) as opposed to technical factors like sequencing depth. For example, Inh MGE remained one of the least age-predictive cell types even after subsampling all pseudobulk data to the same depth, whereas higher-performance cell types, such as Exc L2/3, approached the performance of a high coverage “pseudo-tissue” constructed by summing a donor’s counts across all cell types (**Fig. S15**). The cell type-specific performance was also not clearly attributable to differences in cell type abundance: MGE and Micro were outperformed by CGE and OPC, cell types with similar or lower cell abundances in our dataset (**Fig 1e**). Lastly, we found no evidence of differences in predicted epigenetic age between ASD cases and neurotypical controls (**SFig. 14b, d**), consistent with a previous array study [42].

### Age-associated remodeling of DNA methylome – pre-defined features

We extended our analysis of age-related changes to the DNA methylome by quantifying changes in pre-defined features (genes, promoters and chromosomal bins; *Methods*), which allowed us to model age numerically and study cytosines at non-CpG dinucleotides. Non-CG methylation is most pronounced in neurons and human brain development is marked by rapid accumulation of mCH in the first few years of life, followed by more gradual accumulation of mCH into early adulthood in bulk analysis of the cortex and sorted neurons [6, 7, 33]. We focused on analysis of CpA- sequence contexts due to its relatively high prevalence and correlation with other non-CH motifs (**SFig. 16**). We found that the trajectory of global mCA accumulation varied significantly by cell type; mCA increased with age in excitatory neurons, astrocytes and MGE-derived interneurons (**Fig. 4a**). In excitatory neurons, specifically L4/5 neurons, mCA accumulation plateaued by 25 years of age (**Fig. 4a, S16a**). This pattern was supported by analysis of age-associated differential methylated 500bp bins (DMBs; **Fig 4b**). We examined common

**Figure 4.**
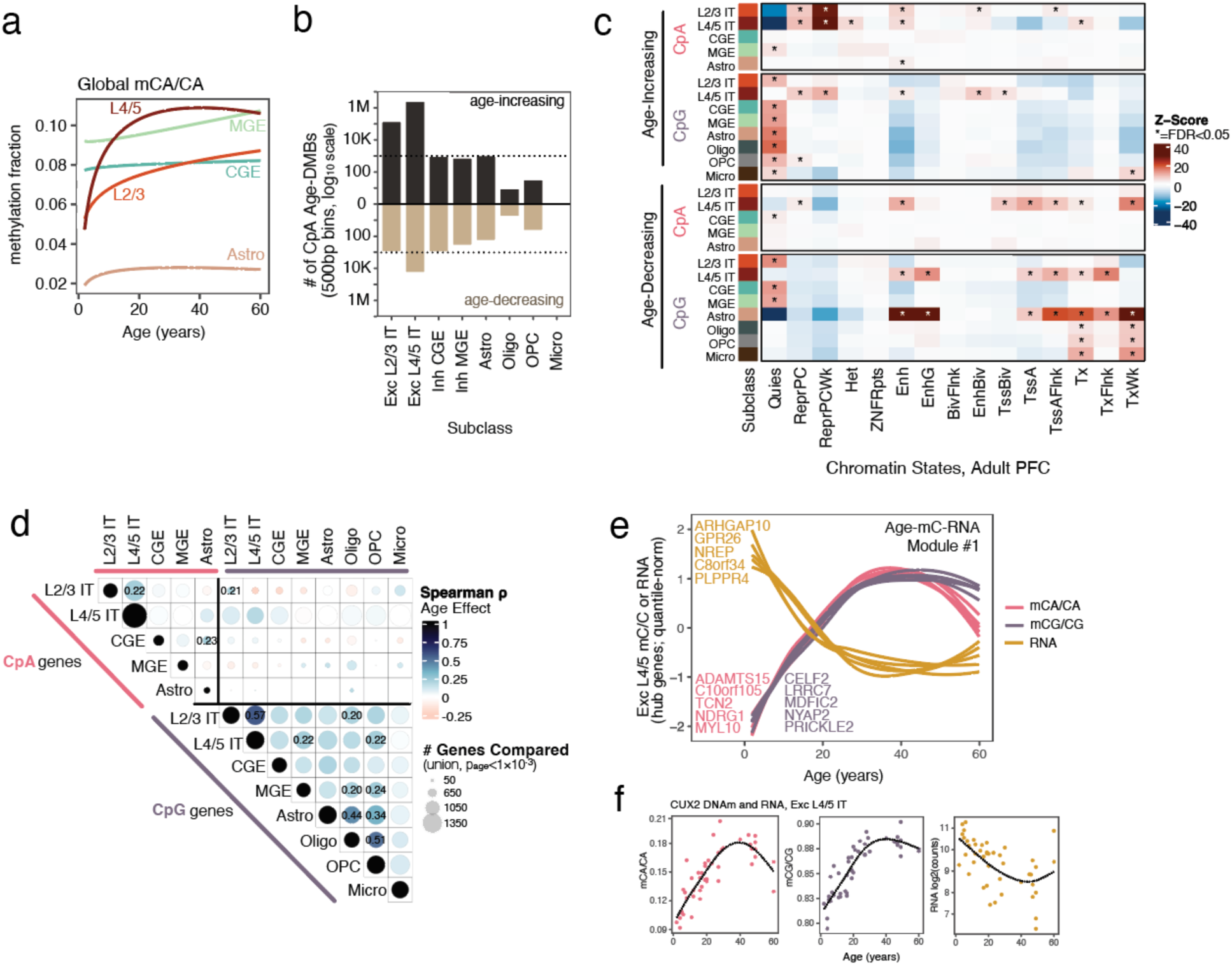
**Age associated CA-methylation**. (a) Global CG-methylation patterns with age in five subclasses. b) The log10-number of age-varying 500bp bins (age-DMBs) detected per subclass and direction. (c) Enrichments of age-DMBs in chromatin state annotations (ChromHMM) derived from adult pre-frontal cortex. d) Correlation in the genewise age effect for pairs of subclass and cytosine sequence context combinations. The value of the Spearman correlation is printed for ρ ≥ 0.20 and indicated by the color of the circle, whereas the size of the circle indicates the number of nominally age-associated bins (union of P<1x10-3 bins in either set) used for comparisons to contextualize the smaller number of age-DMBs in some subclasses and low overlap of CA- and CG- varying bins. (e) Example of multi-‘omic network module generated in Exc L4/5 IT from the paired methylome and RNA data. Each network module contains genes maximally correlated both with age and etween-‘omes to prioritize age-associated features. The genes plotted are among the top features by connectivity. (f) Example of developmental gene CUX2 showing anti-correlation between methylation (mCA/CA and mCG/CG) and gene expression levels (log2-normalized counts, adjusted for library size) in donor seudobulks. methylation patterns over age in bins (“trajectory clustering”, *Methods*) and found ≥75% of CA- DMBs in Exc L2/3 neurons plateaued to mCA/CA > 0.60 by age 25; as opposed to a minority of bins monotonically increasing with age or the 22% of plateauing bins in Inh MGE (**Fig S16c**). By contrast, inhibitory neurons derived from MGE or CGE only modestly gained mCA in the age range sampled in this study, although rapid gain of mCA has been reported earlier in development [62].

The global accumulation of mCA with age was particularly dramatic in Exc L4/5 IT neurons, wherein 89% of bins tested (2,522,864 / 2,253,833 bins in this cell type) significantly gained methylation with age. In a complementary analysis, we included each pseudobulk’s global methylation fraction as a covariate in our regression models and re-identified chromosomal bins associated with age. We found that the majority of DMBs varied in tandem with the global mC level. In excitatory neurons only 5.5% (L2/3) and 0.4% of (L4/5) of DMBs remained significantly associated with age after adjusting for global methylation. That is, most age-varying CA-effects do not deviate from this genome-wide plateau pattern.

We then examined enrichments between genomic annotations and age-DMBs, first assessing transposable elements, which are both theorized to be a destabilizing element in aging [74] and to be regulated by CH-methylation [75]. In excitatory neurons, mCA accumulated with age in transposable elements including Long Terminal Repeats (LTR) and Short Interspersed Nuclear Elements (SINE/Alu), while mCG accumulated in LINE elements with age (**Fig S16d**). We then examined overlap of age-DMBs with chromatin states. In excitatory neurons, regions gaining mCA (and mCG) with age significantly overlapped regions of inactive chromatin, marked by polycomb repression and heterochromatin; whereas regions losing mCG with age were more likely to overlap active regions such as transcriptional start sites (**Fig 4c**). However, this finding should be contextualized by the relative paucity of mCA-loss over age versus gain (**Fig 4a,b**).

We then examined genes and promoters whose methylation varied with age (DMGs and DMPs, respectively). CA- and CG- methylation was more correlated within genes and promoters (median Spearman ρ = 0.17, up to max of 0.57 between excitatory neurons; **Fig 4d**), than in chromosomal bins (ρ = 0.07, max 0.48). Age-associated changes in gene body mCG were often shared across cell types (**Fig. 4d**). Genes losing mCG and/or mCA with age were enriched for pathways (GSEA, FDR < 0.05) related to the cellular response to stress (e.g. DNA damage response, cellular response to stress), oxidative phosphorylation, chromatin remodeling and synapse organization. Similarly, across multiple cell types, genes gaining mC with age implicated G-protein signaling and the adaptive immune response.

We used a semi-supervised (i.e., donor age-aware) sparse canonical correlation network approach [76] to identify small networks whose features were correlated with age and across modalities (age-mC-RNA networks, *Methods*). In the lead module in Exc L4/5 neurons (**Fig 4e**), we find significant enrichment in genes related to glutamatergic synapses, dendritic spine development, and axonogenesis that decrease in expression with age while accumulating methylation within the gene body (e.g., *GRM3, MCTP1, RELN, GRM7, HOMER1, PLPPR4, CASK, ROBO2, RELN*). This module also includes TFs active in development (ex. *MEF2C*, *CELF2*, and *CUX2)* that exhibit high inverse correlations between methylation and gene expression over age (**Fig 4f**).

### Exploration of age-dependent ASD effects on the methylome

We next leveraged the broad age range of donors in this dataset to perform exploratory analyses on diagnosis effects that vary with age, in contrast to pan-age ASD effects (examples in **Fig. 5a**, **Fig S16a-b**). We again turned to modeling pre-defined features (promoters, genes, chrom500 bins) to be able to robustly model age and age-diagnosis interaction effects (*Methods*). As an example, the promoter of *STXBP1,* a gene that has been confidently linked to ASD and Ohtahara

**Fig 5.**
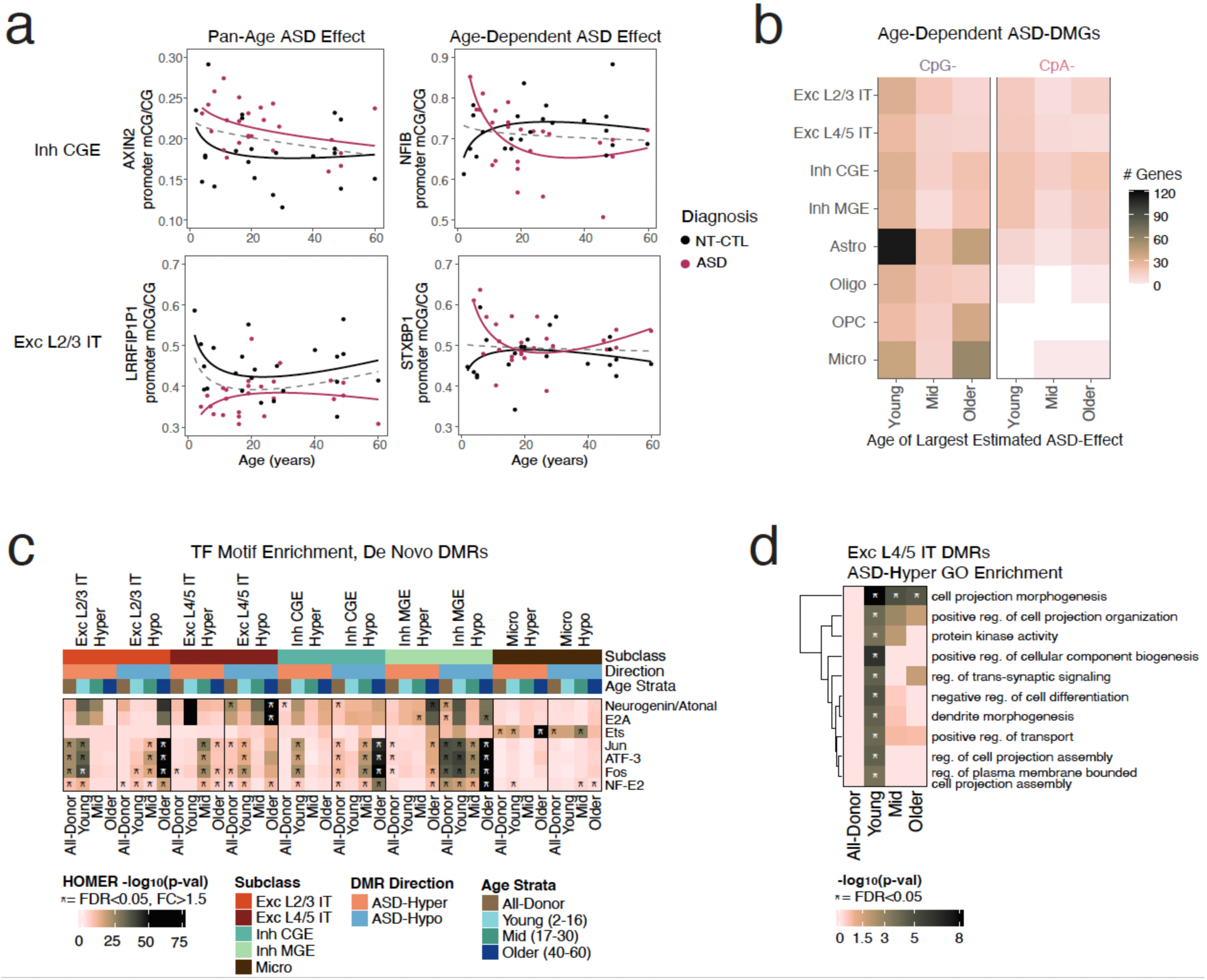
Assessing age-varying methylome associations with ASD. (a) Examples of pan-age and age-dependent diagnosis effects in promoters (ASD-DMPs). (b) For each age-dependent gene (ASD-DMG), the estimated difference between ASD cases and controls was estimated based on the beta regression model of methylation levels, then assigned to the strata for which the bin has highest absolute effect size. (c) TF motif enrichments reported for pan-age *de novo* ASD-DMRs in Fig. 2C are checked for enrichment in age-stratified ASD-DMRs. Asterisk indicates significance and high effect size (FDR < 0.05, FC > 1.5) (d) Example GO enrichments by age strata for Exc L4/5 hypermethylated DMRs. Asterisk indicates statistical significance (q-value < 0.05).

Syndrome [77, 78], is hyper-methylated specifically in Exc L2/3 neurons in young autistic donors (**Fig. 5a**). We considered whether age-dependent diagnosis effects were reflected at the transcriptional level, but did not detect any significant age-dependent diagnosis effects in this dataset or in gene expression data from a larger scRNAseq study ([39], **SFig. 17**).

We detected similar numbers of age-dependent ASD effects in each cell type (**Fig. S18a**), but the timing of effect varied between cell types: the largest mean differences observed in neuronal cell types and astrocytes were concentrated in young donors (**Fig. 5b, S18d**). In contrast to the promoter-enriched pan-age diagnosis effects detected both in de novo ASD-DMRs (**Fig. 2d**) and DMBs (**Fig. S16e**), age-dependent ASD-DMBs were enriched in intergenic regions and quiescent or chromatin-repressed regions (**Fig S18e**).

Given the extensive effects of age on the DNA methylome, we also repeated the *de novo* ASD- DMR discovery procedure (including rigorous permutation of case-control labels, **Fig. 2a**) within three age-stratified subsamples of the cohort to further control for age and potentially detect age-dependent patterns (**Fig. S19a-b**). Age-stratified diagnosis DMRs show a similar distance to the closest sequence variants relative to all-donor DMRs, suggesting that age-stratified diagnosis DMRs are not more likely to be caused by genetic variation (**Fig. S19c**). In each age strata, we again observe enrichment in promoters (**Fig. S10b**) and enrichment of the AP-1/Fos TF family motifs in hypomethylated DMRs in MGE-derived interneurons (**Fig 5c, S19d**), highlighting the robustness of these effects. However, we also detect additional strata-specific motifs: for example, ETS-like TF motifs that were specifically enriched in hypermethylated DMRs in microglia of adult donors (**Fig. 5c**). We performed gene ontology (GO) enrichment analysis on promoters implicated by age-stratified diagnosis DMRs and observed stronger GO enrichment than when called in all-donor DMRs. As an example, DMRs that were hypermethylated in ASD in Exc L4/5 IT neurons across all donors showed no significant GO enrichment, while those identified in young donors were enriched in terms related to neuron projection development, dendrite morphogenesis, and cell differentiation (**Fig. 5d**).

## Discussion

Identifying molecular phenotypes associated with ASD is an essential step towards understanding the underlying biology and developing novel therapeutics. In addition, by implicating specific cell types in pathophysiology, we move closer to linking molecular phenotypes to circuit function and behavior. In this work, we perform the first single-cell analysis of DNA methylation in ASD, profiling DNA methylation and gene expression in the same individual cortical cells derived from 49 donors across the lifespan. We report moderate differences associated with diagnosis that do not explain differential gene expression, but do implicate altered methylation of gene regulatory sites, including fetal brain enhancers and cell type specific candidate cis regulatory elements in the adult brain. We describe profound changes to the methylome associated with age and highlight age-dependent analysis of epigenetic changes as a promising avenue for future investigation in ASD research. The concordance between age-associated changes we report and those previously published supports the validity of our analytic framework.

In this work, we present comprehensive analysis of cell type resolved DNA methylation patterns in idiopathic ASD. ASD-DMRs were generally hypomethylated, which is consistent with loss of function mutations that have been observed in patient mutations in *DNMT3A* [30]. We observed significant overlaps between ASD-DMRs and gene regulatory regions identified in a series of independent studies [62, 63, 65]. ASD-DMRs, across cell types, overlapped promoters more than expected by chance, even after accounting for potential technical bias from CG-content.

This observation is also consistent with promoter enrichment observed in differentially methylated regions identified in a bulk, whole-genome bisulfite analysis of ASD cortical tissue [47]. ASD-DMRs and all alternative (age- and cell type related) DMR sets called in this study significantly overlapped fetal brain enhancers [63] reinforcing the idea that much of the DNA methylation landscape is established before 2 years of age [6].

Both mCG and mCH accumulation can inhibit gene expression [33, 79]; though, the correlation between DNA methylation levels regulatory elements and gene expression varies significantly across cell types [6]. Gene and promoter methylation levels were not correlated with differential gene expression in ASD and, in aging, most genes accumulated methylation with age, independent of expression levels. We considered whether gene methylation levels might reflect historical gene expression by assessing expression of genes implicated by *de novo* DMRs in prenatal and postnatal samples. We did not, however, observe elevated expression levels in prenatal samples. *De novo* ASD-DMRs overlapped candidate cis-regulatory regions, defined by accessible chromatin, in their corresponding adult cell types [65], as well as fetal brain enhancers. Taken together, our results suggest that differential DNA methylation impacts regulatory sites that are active in both the fetal and adult brain, and that ASD may impact cell-type specific gene regulatory programs.

Activity dependent immediate early genes, like *Fos*, have been implicated in ASD previously [39, 59] and their dysregulation is consistent with a transcriptional signature of decreased neuronal activity. This is further supported by differential methylation of Fos family motifs observed in ASD in this dataset. If the methylation of Fos/AP-1 motifs reflects activity, our data suggests increased activity of MGE-derived inhibitory neurons and decreased activity of upper-layer excitatory neurons (particularly in young donors) which is supported by single-cell gene expression analysis [39]. While the binding of transcription factors to TGA*TCA motifs is sensitive to methylation [58] and neuronal activity can influence methylation [80, 81], direct assessment of the relationship between Fos/AP-1 family motif enrichment and activity are needed. The overlap between regions and genes implicated by differential methylation in ASD and those implicated by other modalities (gene expression, H3K27Ac binding) was limited with the exception of hyper-methylated DMRs in microglia. These DMRs functionally implicated pathways related to morphology and immune signaling and overlapped down-regulated H3K27Ac sites in bulk ASD cortex [41]. Differential methylation in microglia in ASD has been suggested previously [40, 47] and supports microglia being one of the most impacted cell types in ASD [39]. In our exploratory analysis of age-dependent diagnosis effects, age-dependent diagnosis effects were observed in microglia, which could be consistent with an age-dependent microglial ASD signature [37] that is compensatory, which has been suggested previously [38, 40].

In addition, we leveraged multimodal single cell data to perform differential abundance and transcriptional analysis in well-defined clusters. Previous work has reported an increased abundance of gray-matter astrocytes and reactive microglia in ASD samples [39, 44]. We hypothesize that we were unable to detect these changes due to the limited number of our non-neuronal cells in this dataset, within which we could not discern different subtypes (ex. gray-matter vs. white-matter resident astrocytes or a subset of reactive microglia). While multiple studies have implicated superficial layer excitatory neurons as a nexus of ASD transcriptional dysregulation [14, 39, 44], we extend this work by our data showing that L2 neurons specifically may exhibit the greatest transcriptional changes.

We investigated age-associated DNA methylation changes from donors aged 2 to 60 at cell-type resolution. Prior studies have investigated changes throughout adulthood, and we expand upon their findings by profiling donors under the age of 20 [68, 69] and considering non-CpG methylation. We report that Exc L2/3 and L4/5 neurons exhibit a rapid increase then plateau before age 25 in methylation levels within both CpG- and CpA- chromosomal bins. This observation reinforces previous reports of the accumulation of mCH, concurrent with synaptic maturation, in bulk cortical tissue [6] and shows the phenomenon is driven by DNA methylation specifically in excitatory neurons.

Consistent with past reports [69], we find that excitatory neurons uniquely accumulate DNA methylation in repressed genomic regions, perhaps as a compensatory response to the loss of heterochromatin with age. Consistent with this, in mouse models of aging, heterochromatin is specifically lost in excitatory neurons, but not glial or inhibitory neurons, resulting in increasing chromatin accessibility and gene expression (including of transposable elements) from said regions [74]. Age-associated increases in methylation may generally compensate for the loss of other repressive mechanisms over age, thus helping protect against age-associated “epigenetic erosion” [82, 83]. Almost all cell types exhibited significant hypomethylation at CCCTC-binding factor (CTCF) binding motifs, which complements mounting evidence of changes to 3D chromatin conformation during neuronal maturation, aging and Alzheimer’s disease [62, 68, 71, 72]. Methylation levels in the inhibitory subclasses profiled here plateau before the ages represented in this dataset and we identified few differentially methylated or expressed features in inhibitory neurons over age. While it is difficult to distinguish between aging process associated with typical maturation and more adverse aging processes in our age-DMRs, we speculate that concomitant loss of methylation at AP-1 family motifs and accumulation of methylation at Neurogenin/Atonal motifs may reflect competition between synaptic activity and neuronal differentiation, as has reported in ATACseq data [84]. Lastly, we assessed the performance of epigenetic age clocks in across different cell types. We found that clock performance varied widely by cell type, as reported by others [68], and suggest that there are opportunities to develop cell-type specific age clocks using single-nucleus data. Further, variable clock performance supports the highly cell-type specific nature of methylome-age associations.

Our study has some important limitations. First, a substantial portion of nuclei did not pass transcriptomic QC. This dropout was particularly pronounced in glial cells, which have lower RNA content than neurons [44]. While we are confident in the quality of our RNA data, which reflects expected cell type, age and diagnosis expression patterns, our dataset is limited in size, broadly limiting our power to detected changes in DNA methylation and specifically limiting our ability to profile non-neuronal cell types. Second, our stringent filtering criteria for selecting pre- defined features for differential methylation testing excluded many expressed genes from analysis. The focus on variable regions may bias our pre-defined region analysis towards regions of the methylome under less regulatory constraint (e.g., regions overlapping non-expressed genes; genome quiescent regions) and result in underestimates of methylome-transcriptome pairing, although *de novo* DMR discovery was not limited by these inclusion criteria. Our ASD-DMRs did not cover enough of the genome to be able to robustly partition heritability, leaving open the question of whether or not regions that impart genetic risk for ASD are differentially methylated. Lastly, we have studied DNA methylation using bisulfite conversion, which is unable to resolve differences between methyl-cytosine and hydroxy- methylation cytosine. The latter is known to be highly dynamic with age [85, 86] and can positively correlate with gene expression, complicating the functional interpretations of our DMRs.

Our data indicate that changes to the DNA methylome, within and beyond CG sites, are moderate in idiopathic ASD and independent of gene expression changes, consistent with previous work in bulk tissue suggesting DNA methylation profiles in ASD were distinct from those identified in other modalities [40]. A more dramatic methylomic signature may be detectable in individuals with syndromic forms of ASD (ex. Rett Syndrome) or rare variation in *DNMT3A*, as has been suggested in bulk analysis [42, 47]. Our results also provide insight into cell type specific changes in brain that accompany aging as well as potential interactions between aging and diagnosis, both of which are exciting areas of future investigation.

## Acknowledgements

Human tissue was received from the NIH NeuroBioBank at the University of Maryland Brain and Tissue Bank and from Autism BrainNet, a resource of the Simons Foundation Autism Research Initiative (SFARI). Autism BrainNet also manages the Autism Tissue Program (ATP) collection, previously funded by Autism Speaks. We are grateful and indebted to the families who donated tissue for research purposes to Autism BrainNet, the ATP and UM-BTB.

This work used computational and storage services associated with the Hoffman2 Shared Cluster provided by UCLA Office of Advanced Research Computing’s Research Technology Group. Flow cytometry experiments were performed in the UCLA Broad Stem Cell Research Center Flow Cytometry Core. Whole-genome sequencing library preparation was performed by Briana Hernandez and the UCLA Technology Center for Genomics and Bioinformatics.

Thanks to Drs. Jason Ernst, Hao Feng, Eran Mukamel for methodological discussion, Dr. Brie Wamsley for discussion and tissue coordination and Thordar Han for preliminary analysis of external ASD datasets.

This work was supported by NIMH grants R01MH125252 and U01MH130995 to C.L.

## Methods

### Tissue collection and donor characteristics

Human tissue was received from the NIH NeuroBioBank at the University of Maryland Brain and Tissue Bank and from Autism BrainNet, a resource of the Simons Foundation Autism Research Initiative (SFARI). Autism BrainNet also manages the Autism Tissue Program (ATP) collection, previously funded by Autism Speaks. We are grateful and indebted to the families who donated tissue for research purposes to Autism BrainNet, the ATP and UM-BTB. We received tissue isolated from the prefrontal cortex (PFC; Brodmann area 9, BA9) of 52 donors. 3 female donors were excluded from further analysis. 1 low RIN donor was excluded from RNA analysis.

Additional information on donor phenotyping (including ADI-R scores) is in the supplemental methods.

### Nuclei isolation from human brain tissue

Small pieces (<50 mg) of tissue were ground in liquid nitrogen with pre-chilled mortar and pestle to generate a homogenous sample and stored at -80°C until use. On the first day of processing, ground tissue was resuspended in nuclei isolation buffer and dounced to break up tissue. Nuclei were then isolated using an iodixanol gradient, counted and stained (using a fluorescent NeuN antibody) prior to being sorted into individual wells of a 384-well plate for processing. Full protocol details are in the supplemental methods.

### mCT processing and library preparation

All solutions were prepared in a laminar flow cabinet, and solution distribution and mixing were performed using Beckman Biomek i7 liquid handler.

sn-mCT-seq processing and library preparation were performed as previously described [49]. A detailed bench protocol is shared through protocol.io (https://www.protocols.io/view/snmcat-v2-x54v9jby1g3e/v2). The following text refers to specific sections in the snmCAT protocol.

Nuclei were kept on ice until sorting (*Step 10*) into pre-cooled 384-well PCR plates (ThermoFisher 4483285) containing reverse transcription reaction reagents (1uL per well, prepared in *Step 9*). We prepared cDNA amplification mix as in *Step 12,* and ran 16 amplification cycles for non-neuronal nuclei and 14 cycles for neuronal nuclei.

We performed bisulfite conversion (*Step 14*) as follows: to each bottle of CT conversion reagent (Zymo D5003-1), we added 7.9 mL of M-solubilization buffer (Zymo D5021-7) and 3 mL of M-dilution buffer (Zymo D5002-2). This mixture was vigorously mixed for at least 15 minutes before the addition of 1.6 mL M-reaction buffer (Zymo D5021-8). 25 µL of this reaction mixture were added to each well of the 384-well plate. Plate were then vortexed, sealed and incubated for 98 °C - 8 min, 64 °C - 3.5 h and then held at 4 °C.

Reaction mixtures (30 µl) were transferred to filter plates (Agilent 201025-100) and mixed with 90 µL of M-Binding Buffer (Zymo D5006-3) before incubating for 5 minutes at room temperature. Plates were then sealed and centrifuged for 5 minutes. Flow-through was discarded, then samples were washed with 30 uL per well of M-Wash Buffer (Zymo D5040-4) and centrifuged again. Flow-through was again discarded. 30 uL of M-Desulphonation Buffer (Zymo D5040-5) was added to each well and the plates were incubated at room temperature for 15 min, then centrifuged. The plates were washed twice with 30 µL / well of M-Wash Buffer, then eluted into a clean 384-well plate with 7 µL EB Buffer (Qiagen 19086) containing 500 nM well-indexed random primers.

A detailed bench protocol for single-cell methylome library preparation is shared through protocol.io (https://www.protocols.io/view/snm3c-seq3-kqdg3x6ezg25/v1). We performed *steps 13 - 22* of the snm3c protocol.

Libraries were sequenced using Illumina Novaseq 6000 instrument on S4 and XP flowcells and 100 bp paired-end configuration with a 2-5% PhiX spike-in.

### Bioinformatic alignment and feature quantification.

The workflow for going from raw fastq files to modality-specific, aligned reads is comprehensively outlined in the following repository: https://github.com/chooliu/snmCTseq_Pipeline. Deviations from analysis performed in the flagship paper [49] include the implementation of paired-end alignment and quantification methods, which increased mapping rate and coverage, while ensuring that read 1 and read 2 features were not quantified twice when insert size is low. Additionally, we deduplicated RNA reads.

Generally,

● Samples submitted for sequencing contain pooled libraries from across 384-well plates. Plate identity is indicated by i5/i7 indices and plate-level demultiplexing is performed prior to the snmCTseq pipeline.
● Plate-level fastqs are demultiplexed by well-specific barcodes prior to trimming of technical sequences (barcodes/adaptors) and low-quality bases (<Q20).
● Reads were aligned to the GENCODE v40 reference genome (GRCh38.p13 genome and genic annotation files)
● Methylome reads were mapped using Bismark [87] v0.24.0 in a two-stage process in order to increase mapping efficiency for PBAT-based reactions and other single-cell methylome preparations [88]: (i) first, reads are mapped in paired-end mode, then (ii) reads unable to map in stage i and singleton reads are mapped in single-end mode. PCR and sequencer optical duplicates using Picard MarkDuplicates [89]. Reads with mapping score ≤ 10 were excluded (samtools view -h -q 10; samtools v1.16.1 [90]). We classified Bismark alignments containing at least three CpH-context cytosines and mCH/CH ≤ 0.50 as DNA reads using the “XM:Z” alignment flag. We then inferred cytosine-level methylation using allcools [91].
● In parallel, RNA-originating reads were mapped with STAR v2.7.10a [92], also using the two-stage (paired-end, followed by single-end) mapping procedure for RNA quantification. Alignments that were duplicates identified by Picard or had mapping score

≤ 10 were excluded. To retain RNA reads, each alignment with at least three CpH- context cytosines and mCH/CH ≥0.90 based on the “MD:Z” flag were retained. RNA counts of each gene were then quantified using featureCounts [93].

### Nuclei-level quality control

We performed nuclei-level quality control checks for RNA and DNA methylation data separately before data integration.

#### Methylome

We used methylome quality control metrics previously established for brain data [49, 91], requiring:

● Bismark mapping rate > 50%,
● >1x10^5^ final aligned reads,
● global methylation fractions of mCG/CG > 0.50, mCH/CH < 0.20, and mCCC/CCC < 0.03, where the last measure is a proxy for bisulfite conversion rate. Global methylation fraction is defined as *f* := ∑c mc/(mc+uc) for all cytosines *c* from the given cytosine sequence context.

#### Transcriptome

Feature counts obtained for a given cell (across single- and paired-end alignments) were aggregated. All downstream steps were performed in Seurat [94]. Nuclei were included in transcriptomic analysis if they met the following criteria:

● 500 < nGenes / cell < 8,000
● % mitochondrial transcripts < 10
● nCount > 10,000
● % intergenic bases < 10 (output from Picard’s RnaSeqMetrics)

These criteria removed almost all nuclei from 1 donor, with low RIN, who was subsequently excluded from RNA analysis.

Genes were included in downstream analysis if they were expressed (count > 0) in at least 10 nuclei.

### Data Clustering, Integration and Annotation

#### Methylome

We used the allcools pipeline (v1.0.8) [91] to perform variable feature selection, dimension reduction and clustering based on CpG- and the CpH- methylation fractions in non-overlapping 100kb genomic bins (on autosomal chromosomes) that had less than 20% base overlap with the ENCODE unified exclusion list (encodeproject.org/files/ENCFF356LFX). We also excluded bins that were outliers in terms of coverage (<200, ≥2000). We then derived principal components (PCs) from variable bins (the top 25^th^ percentile), which were used in UMAP embedding. Clusters were determined using an iterative approach in which leiden clustering was repeated 1,000 times with random seeds and an 80% subsample of the nuclei (ConsensusClustering; *k* neighbors = 20, resolution = 1.5), resulting in a tally of 62 leiden clusters across samples. Clusters were annotated on the basis of gene body (CpG and CpH) hypomethylation in canonical marker genes [10, 95].

#### Transcriptome

Variable feature selection, dimension reduction and clustering were performed using the default Seurat pipeline [94]. Counts were scaled and normalized using the *LogNormalize* method. We derived PCs from 3,000 variable genes and used 20 PCs to find neighbors and in UMAP embedding (leiden resolution = 0.8). We identified 17 clusters, using only gene expression data, and annotated them on the basis of canonical marker gene expression and predicted subclass labels based on data from [95] embedded in Azimuth [96].

### Modality integration and label transfer

We integrated transcriptome and methylome data from individual nuclei using a weighted nearest-neighbor approach implemented using methylome and RNA PCs and default parameters in the *FindMultiModalNeighbors* function in Seurat [96]. Dual modality clusters were annotated as described above.

To annotate nuclei that passed only methylome QC using cell type labels derived from both data types, we transferred cluster labels constructed from the joint RNA-methylome WNN embeddings to the methylome-only nuclei. We predicted cluster labels (model outcome) from the 100kb-mCH and 100kb-mCG principal components (model predictors) using the weighted k-nearest neighbors algorithm [97]. In the final hold-out test set, accuracy was 98.7% at the subclass annotation level. After applying the final model to methylome-only nuclei, we excluded nuclei that either had “Unannotated” labels or were members of methylation-only based leiden clusters with majority “Unannotated” labels, resulting in the final nuclei count of 65,023 single-nucleus methylomes. Full details in supplemental methods.

### Differential gene expression analysis

In our primary analysis, data were aggregated to pseudobulk and differential gene expression analysis was performed using DESeq2 [98]; both steps were implemented using muscat [99].

Clusters, donors and features were filtered using the following criteria, before pseudobulk aggregation:

● Tested clusters with greater than 500 total nuclei
● Generated pseudobulk count matrix from donors with at least 10 nuclei in a given cell type
● Tested genes expressed (count > 0) in 20% of nuclei within a cluster

We estimated and controlled for variables independent of the diagnosis effect using RUVseq [100] (using the residual-based method, RUVr; k = 5).

To perform the downsampling analysis in *SFig. 6e*, we randomly sampled 20 cells per donor per cell type across 500 permutations, repeated the procedure described above and reported the number of DEGs (padj < 0.1).

To assess whether we could recapitulate previously published results, we implemented the same type of differential expression models as the reference paper, as detailed in supplemental methods.

### Whole Genome Sequencing, genotyping and ancestry inference

Genomic DNA was extracted using Zymo’s Quick-DNA mini-prep plus kit (#4068) and libraries were prepared using Roche’s KAPA Hyper kit (#07962363001) and sequenced on a Novaseq6000 (in 2x100bp and 2x150bp configurations). Reads were processed using the GATK4 best practices workflow [101] for short variant germline discovery (SNPs and indels; accessed in January 2023). The reads were aligned using bwa-mem v123. The final mean autosomal alignment depths were 40.9X ± 10.3 (across-sample mean and sd), with no significant mean difference in depth or other quality metrics by diagnosis or age in years (t-test p-value > 0.59).

WGS genotypes were joined to the 1000 Genomes reference panel (Byrska-Bishop [102] variant- filtered and phased release; internationalgenome.org/data-portal/data-collection/30x-grch38), we removed variants that were lower frequency (MAF < 0.05) or overlapped previously reported regions of long-range LD (e.g., HLA region; regions extracted from [103]. We ran principal component analysis on the resulting joined genotypes (flashPCA v2.0 [104]) and retained the first two PCs for modeling. Continental ancestry was assigned to each donor using the weighted k-nearest neighbor algorithm, and, in parallel, using ADMIXTURE. Both PCA and ADMIXTURE continental ancestry predictions were consistent with biobank records of donor Self-Identified Race/Ethnicity, where available.

### Genotype Masking of the Methylome

Donor genetic deviation from the reference genome introduces noise into methylation quantification in bisulfite sequencing. To minimize this effect, we set the methylation counts and coverage of a given donor’s cytosines to zero if the cytosine overlapped a (1) loss-of-cytosine, a (2) change in the cytosine’s +1 context from CG>CH or CH>CG, or (3) an indel. These “genotype- masked” calls were subsequently used prior to any differential methylation testing by diagnosis, age, or cell-type.

### Differential Methylation Identification Strategies.

After removing cytosines overlapping the ENCODE Unified Exclusion list or with zero coverage in most samples (zero in ≥75% of samples), we ran DSS with the default prior and a 500bp smoothing window to call *de novo* DMRs. We aggregated the single cytosine results into larger candidate regions by aggregating sites within ±250bp, then retaining regions with at ≥2 multiple testing adjusted significant cytosines (Storey q-value < 0.05) and at least one significant cytosine with methylation fraction (*f)* estimated effect size |Δ| > 0.10.

For the ASD-DMRs, we excluded regions that were regularly detected in 500 label-shuffling permutations (more than 20% of the time). We repeated this process for all-donor ASD-DMRs, as well as age-stratified subsets of the data: “young” (2-16), “mid” (17-30), and “older” (40-60) samples within a given subclass as a method to control for age effects. The threshold delineating young and mid for age-stratified ASD-DMRs were selected to approximately maintain the same sample sizes in each group (n=19, n=20, n=13).

Using this DSS to DMR procedure, we additionally called age-DMRs within each cell type by pairwise testing between the young, mid, and older age strata. Unless otherwise noted, we primarily report and analyze the union of the three pairwise tests within each subclass. To call cell-type DMRs, we repeated the DMR calling procedure between each pairwise comparison of the eight cell types (e.g., Astro versus Exc L2/3, Astro versus Inh MGE, etc.). A cell-type DMR for subclass *s* is identified as hypo-methylated in cell-type *s* compared to ≥5 other subclasses.

### DMR Calling with Pre-Defined Features

We considered pre-defined features of *<u>genes</u>* (GENCODE v40 protein coding and non-coding), *<u>promoters</u>* (+2kb, -1kb from first exon TSS), and full-genome non-overlapping *<u>genomic bins</u>* tiling the genome (500bp). For each region *r* within a given pseudobulk (subclass *s and* donor *i*) we summed the values of methylated counts (*m*c) and unmethylated counts (*u*c) for each cytosine *c* within the region: i.e., m*r = ∑cmc* and u*r = ∑cuc*. The *mr* and *ur* values were calculated separately for cytosines *c* within two dinucleotide sequence contexts (CpA- and CpG- cytosines), with the focus on CpA-methylation as it is the most prevalent form of non-CpG methylation and is correlated with other forms of CpH-methylation. We did not consider strandedness in any analysis and counts for CG- sites were merged to the “+” strand during pseudobulking.

Regions exhibiting high between-donor variability (IQR > 0.05 across donors in methylation fraction *fr*, where *fr* := m*r*/(*mr* + *ur*)) and sufficient coverage (≥5 counts in at least 75% of donors for genes and promoters; ≥2 counts in 75% of donors for 500bp bins) were modeled as the outcome of a beta-binomial model (*aod* R package implementation, logit-link) with diagnosis (indicator variable for case), age (coded as linear and log(age) terms, then scaled and centered), and an diagnosis-by-age interaction (ASD-indicator, times linear- and log-age), and population stratification (first two genotype PCs) as model covariates. Features that failed to converge after 25,000 iterations were excluded; these were frequently bins at the edge of the parameter space and often exhibited quasi-complete separation (i.e., strong age or diagnosis effects, but computationally unstable due to low between-donor variance from mean *f* close to 0 or 1).

We tested the significance of each regression coefficient with a Wald test, followed by Storey q-value < 0.05 multiple testing correction to define statistical significance. We defined two forms of diagnosis effect. First, <u>pan-age ASD effects</u> are defined by significant diagnosis coefficient, but no significant ASD-by-age interaction effects. In contrast, <u>age-dependent ASD effects</u> are defined by a significant interaction term (q-value < 0.05). Due to the correlative nature of these analyses, the significant interactions can also be conceptualized as a different pattern of aging observed between ASD cases and controls. We also defined <u>age-effects</u> as significant linear- or log-age effects, focusing on interpretation of age-bins without significant interaction terms.

### Annotating DMRs to genes and motif enrichment analysis

All differentially methylated regions (DMRs) were annotated to genes using the *annotatePeaks* function in the HOMER suite [57], using the same genome annotation file as during alignment. In addition, DMRs that overlapped candidate cis regulatory elements identified in [65] were annotated with gene assignments made in [65]. If a DMR overlapped multiple cCREs, we prioritize annotations as in *annotatePeaks*. Both sets of annotations are provided in Supplemental File 5.

DMR motif enrichment analysis was performed using the *findMotifsGenome* function in HOMER 1. [57] and the fasta file corresponding to the same genome used for alignment.

### Gene ontology and gene set enrichment analysis

#### Transcriptome

We performed gene set enrichment analysis of differentially expressed genes using the *fgseaMultilevel* function of fgsea [105] and the biological process (BP) and molecular function (MF) subsets of the C5 ontology gene sets available here: https://www.gsea-msigdb.org/gsea/msigdb/collections.jsp (download July 2024). Input lists were ordered by effect size (log FC) * -log10(pvalue) and we considered gene sets between 25 and 500 genes in size.

#### Methylome

We assessed gene ontology (GO) enrichment analysis on DMRs annotated to transcriptional start sites using gprofiler2 [106]. We observed an association between genic DMRs (in transcriptional start and termination sites, exons and introns) and gene length. To adjust for this, we performed GO analysis of genic DMRs using goseq [107].

### Enrichment analyses

Enrichment analyses were performed using regioneR / regioneReloaded [108, 109].

To assess overlap between DMRs and genomic features, we compared the number of observed overlaps to a distribution of values (# overlaps) across 1E6 permutations, sampling from a GC-content and coverage-matched background.

To assess overlap between DMRs and regions from published datasets, we quantified the number of overlaps using the *permTest* function in regioneR across 500 - 1E5 permutations resampling regions (*resampleRegions*) from a universe of regions with coverage in our methylation analysis (a subset of background sets were also length matched to the target set).

### Cross-Modality Correlation Analyses.

The criteria outlined in *DMR Calling with Pre-Defined Features* precluded analysis of gene body mCA in many protein-coding genes. To be able to correlate gene body mC and gene expression (*SFig. 13f*), we also assessed promoter and gene body cytosine methylation following the steps outlined in Section 4.8 of the edgeR user guide : https://www.bioconductor.org/packages/devel/bioc/vignettes/edgeR/inst/doc/edgeRUsersGuide. pdf.

Due to limited overall pairwise correlations between methylation and RNA features, we additionally attempted to prioritize genic features of interest by generating age-mC-RNA networks with SmCCNet [76], a semi-supervised (donor age-aware) network approach that maximizes across-modality canonical correlations with a L1-penalty on the gene weights to induce sparsity in the networks. The network adjacency matrix is constructed using the canonical weights; that is, each network module contains features correlated with age and/or across ‘omes. Note that genes selected do not necessarily have high *rg*. Each methylome gene fraction (CpA- and CpG-gene fractions; concatenated across sequence context and restricted to genes tested i.e., IQR *fr* > 0.05) and RNA (log2-normalized counts) was quantile-normalized across donors and residualized for diagnosis effect (mean ASD effect regressed out) prior to network construction. Only protein-coding genes were included. Enrichr [110] was used to perform over-representation analysis on Gene Ontology [111] Biological Process terms on genes present in each module (GO_Biological_Process_2025; Benjamini-Hochberg FDR < 0.10).

### Age Trajectory Clustering.

We selected the top 2,500 age-DMBs of each subclass by likelihood ratio (full beta-binomial model compared to reduced model without age or interaction terms) to identify sets of bins exhibiting similar patterns with age. To focus on The model coefficients were used to predict methylation values (assuming NT-CTL samples, age 2 to 60) was performed using dtwclust v1.23- 1 [112] with the dtw_basic distance and the Partitioning Around Medoids clustering algorithm with *k* = 10, a larger *k* to err on identifying more unique patterns.

### Epigenetic Age Clocks.

The probe lists were extracted from the 353-probe and 111-probe Horvath age clocks [5] and the cortex age clock [73], then converted to their corresponding GRCh38 single-cytosine coordinates using sesameData v1.20.0 [113]. We extracted a ±1bp region in each of the donor-by-subclass methylome pseudobulk to calculate the corresponding methylation fractions as a proxy for *fp* for probe *p* (often “β” in array literature) at each probe, with the ±1bp expansion due to the high cross-strand symmetry and correlation of CpG-methylation. The epigenetic age estimate was then calculated for each of the pseudobulks, per usual clock calculations ^Age = g(*∑pwpfp)* where *wp* are the probe weights (LASSO coefficients) and where *g(x)* is the calibration function shared by both clocks to account for the faster epigenetic changes reported before adult maturation [5, 73]: g(x) = (1+ τ)exp(x)-1 for x < 0 and (1 + τ)*x + τ for x ≥ 0 for τ = 20.

We evaluated evidence of cell-type specific epigenetic aging differences between ASD and CTL by evaluating whether the residual epigenetic age (^Age - ^Ageloess) differed by group with a t-test, where the loess mean trend (^Ageloess, span=1) between the true age and epigenetic age was used as a “calibration curve” correcting for prediction errors between the clock age and chronological age specific to our dataset. We also evaluate the performance of each clock in predicting the biobank ground truth donor age by tabulating the Mean Absolute Error (MAE) and Spearman correlation between the predicted and ground truth ages.

We performed three additional analyses to assess the robustness of our findings to coverage. First, we constructed a high coverage “pseudo-tissue” by calculating a given donor’s methylation fraction from all eight major subclasses together; as each clock is validated in bulk data, this composite thus represents a performance bound due to technical effects of transferring the array-based technology to our sequencing-based measurements. Second, we hypothesized that given the spatial correlation of CG-methylation—particularly with the high proportion of clock probes overlapping either a promoter or CpG-island (55% in cortex clock, 81% in Horvath-353)—that padding fractional methylation calculations to include sites within a ±*w* base window of each probe may reduce noise. We thus also evaluated the impact of padding the *f* calculation with all cytosines in a ±25bp, ±50bp, or ±100bp range. Third, we calculated the mean across-pseudobulk coverage at each probe site and repeated all epigenetic age prediction when each pseudobulk and pseudo-tissue were sampled with replacement to this mean coverage level. This coverage resampling was performed 100 times.

## References

1. Hon, G.C., et al., Epigenetic memory at embryonic enhancers identified in DNA methylation maps from adult mouse tissues. Nat Genet, 2013. 45(10): p. 1198–206.

2. Schultz, M.D., et al., Human body epigenome maps reveal noncanonical DNA methylation variation. Nature, 2015. 523(7559): p. 212-6.

3. Ziller, M.J., et al., Charting a dynamic DNA methylation landscape of the human genome. Nature, 2013. 500(7463): p. 477-81.

4. He, Y., et al., Spatiotemporal DNA methylome dynamics of the developing mouse fetus. Nature, 2020. 583(7818): p. 752-759.

5. Horvath, S., DNA methylation age of human tissues and cell types. Genome Biol, 2013. 14(10): p. R115.

6. Lister, R., et al., Global epigenomic reconfiguration during mammalian brain development. Science, 2013. 341(6146): p. 1237905.

7. Price, A.J., et al., Divergent neuronal DNA methylation patterns across human cortical development reveal critical periods and a unique role of CpH methylation. Genome Biol, 2019. 20(1): p. 196.

8. Lister, R., et al., Human DNA methylomes at base resolution show widespread epigenomic differences. Nature, 2009. 462(7271): p. 315-22.

9. Loyfer, N., et al., A DNA methylation atlas of normal human cell types. Nature, 2023. 613(7943): p. 355-364.

10. Luo, C., et al., Single-cell methylomes identify neuronal subtypes and regulatory elements in mammalian cortex. Science, 2017. 357(6351): p. 600-604.

11. Luo, C., P. Hajkova, and J.R. Ecker, Dynamic DNA methylation: In the right place at the right time. Science, 2018. 361(6409): p. 1336-1340.

12. Smith, Z.D., S. Hetzel, and A. Meissner, DNA methylation in mammalian development and disease. Nat Rev Genet, 2025. 26(1): p. 7–30.

13. De Rubeis, S., et al., Synaptic, transcriptional and chromatin genes disrupted in autism. Nature, 2014. 515(7526): p. 209-15.

14. Parikshak, N.N., et al., Integrative functional genomic analyses implicate specific molecular pathways and circuits in autism. Cell, 2013. 155(5): p. 1008–21.

15. Pinto, D., et al., Convergence of genes and cellular pathways dysregulated in autism spectrum disorders. Am J Hum Genet, 2014. 94(5): p. 677–94.

16. de la Torre-Ubieta, L., et al., The Dynamic Landscape of Open Chromatin during Human Cortical Neurogenesis. Cell, 2018. 172(1-2): p. 289–304 e18.

17. Satterstrom, F.K., et al., Large-Scale Exome Sequencing Study Implicates Both Developmental and Functional Changes in the Neurobiology of Autism. Cell, 2020. 180(3): p. 568–584 e23.

18. Lei, H., et al., De novo DNA cytosine methyltransferase activities in mouse embryonic stem cells. Development, 1996. 122(10): p. 3195–205.

19. Li, E., T.H. Bestor, and R. Jaenisch, Targeted mutation of the DNA methyltransferase gene results in embryonic lethality. Cell, 1992. 69(6): p. 915–26.

20. Okano, M., et al., DNA methyltransferases Dnmt3a and Dnmt3b are essential for de novo methylation and mammalian development. Cell, 1999. 99(3): p. 247–57.

21. Robert, M.F., et al., DNMT1 is required to maintain CpG methylation and aberrant gene silencing in human cancer cells. Nat Genet, 2003. 33(1): p. 61–5.

22. Winkelmann, J., et al., Mutations in DNMT1 cause autosomal dominant cerebellar ataxia, deafness and narcolepsy. Hum Mol Genet, 2012. 21(10): p. 2205–10.

23. Tatton-Brown, K., et al., Mutations in the DNA methyltransferase gene DNMT3A cause an overgrowth syndrome with intellectual disability. Nat Genet, 2014. 46(4): p. 385–8.

24. Weinberg, D.N., et al., The histone mark H3K36me2 recruits DNMT3A and shapes the intergenic DNA methylation landscape. Nature, 2019. 573(7773): p. 281-286.

25. Amir, R.E., et al., Rett syndrome is caused by mutations in X-linked MECP2, encoding methyl-CpG-binding protein 2. Nat Genet, 1999. 23(2): p. 185–8.

26. Feliciano, P., et al., Exome sequencing of 457 autism families recruited online provides evidence for autism risk genes. NPJ Genom Med, 2019. 4: p. 19.

27. Ruzzo, E.K., et al., Inherited and De Novo Genetic Risk for Autism Impacts Shared Networks. Cell, 2019. 178(4): p. 850–866 e26.

28. Sanders, S.J., et al., Insights into Autism Spectrum Disorder Genomic Architecture and Biology from 71 Risk Loci. Neuron, 2015. 87(6): p. 1215–1233.

29. Zhou, X., et al., Integrating de novo and inherited variants in 42,607 autism cases identifies mutations in new moderate-risk genes. Nat Genet, 2022. 54(9): p. 1305–1319.

30. Christian, D.L., et al., DNMT3A Haploinsufficiency Results in Behavioral Deficits and Global Epigenomic Dysregulation Shared across Neurodevelopmental Disorders. Cell Rep, 2020. 33(8): p. 108416.

31. Li, J., et al., Dnmt3a knockout in excitatory neurons impairs postnatal synapse maturation and increases the repressive histone modification H3K27me3. Elife, 2022. 11.

32. Stadler, M.B., et al., DNA-binding factors shape the mouse methylome at distal regulatory regions. Nature, 2011. 480(7378): p. 490-5.

33. Guo, J.U., et al., Distribution, recognition and regulation of non-CpG methylation in the adult mammalian brain. Nat Neurosci, 2014. 17(2): p. 215–22.

34. Gandal, M.J., et al., Broad transcriptomic dysregulation occurs across the cerebral cortex in ASD. Nature, 2022. 611(7936): p. 532-539.

35. Gandal, M.J., et al., Transcriptome-wide isoform-level dysregulation in ASD, schizophrenia, and bipolar disorder. Science, 2018. 362(6420).

36. Gupta, S., et al., Transcriptome analysis reveals dysregulation of innate immune response genes and neuronal activity-dependent genes in autism. Nat Commun, 2014. 5: p. 5748.

37. Parikshak, N.N., et al., Genome-wide changes in lncRNA, splicing, and regional gene expression patterns in autism. Nature, 2016. 540(7633): p. 423-427.

38. Voineagu, I., et al., Transcriptomic analysis of autistic brain reveals convergent molecular pathology. Nature, 2011. 474(7351): p. 380-4.

39. Wamsley, B., et al., Molecular cascades and cell type-specific signatures in ASD revealed by single-cell genomics. Science, 2024. 384(6698): p. eadh2602.

40. Ramaswami, G., et al., Integrative genomics identifies a convergent molecular subtype that links epigenomic with transcriptomic differences in autism. Nat Commun, 2020. 11(1): p. 4873.

41. Sun, W., et al., Histone Acetylome-wide Association Study of Autism Spectrum Disorder. Cell, 2016. 167(5): p. 1385–1397 e11.

42. Wong, C.C.Y., et al., Genome-wide DNA methylation profiling identifies convergent molecular signatures associated with idiopathic and syndromic autism in post-mortem human brain tissue. Hum Mol Genet, 2019. 28(13): p. 2201–2211.

43. Wu, Y.E., et al., Genome-wide, integrative analysis implicates microRNA dysregulation in autism spectrum disorder. Nat Neurosci, 2016. 19(11): p. 1463–1476.

44. Velmeshev, D., et al., Single-cell genomics identifies cell type-specific molecular changes in autism. Science, 2019. 364(6441): p. 685-689.

45. Nardone, S., et al., DNA methylation analysis of the autistic brain reveals multiple dysregulated biological pathways. Transl Psychiatry, 2014. 4(9): p. e433.

46. Nardone, S., et al., Dysregulation of Cortical Neuron DNA Methylation Profile in Autism Spectrum Disorder. Cereb Cortex, 2017. 27(12): p. 5739–5754.

47. Vogel Ciernia, A., et al., Epigenomic Convergence of Neural-Immune Risk Factors in Neurodevelopmental Disorder Cortex. Cereb Cortex, 2020. 30(2): p. 640–655.

48. Yap, C.X., et al., Brain cell-type shifts in Alzheimer’s disease, autism, and schizophrenia interrogated using methylomics and genetics. Sci Adv, 2024. 10(21): p. eadn7655.

49. Luo, C., et al., Single nucleus multi-omics identifies human cortical cell regulatory genome diversity. Cell Genom, 2022. 2(3).

50. Dann, E., et al., Differential abundance testing on single-cell data using k-nearest neighbor graphs. Nat Biotechnol, 2022. 40(2): p. 245–253.

51. Hodge, R.D., et al., Conserved cell types with divergent features in human versus mouse cortex. Nature, 2019. 573(7772): p. 61-68.

52. Staff, N., *List of genes having strong statistical support for association to mental illnesses*. 2023, National Institute of Mental Health National Institutes of Health.

53. Clemens, A.W., et al., MeCP2 Represses Enhancers through Chromosome Topology-Associated DNA Methylation. Mol Cell, 2020. 77(2): p. 279–293 e8.

54. Feng, H., K.N. Conneely, and H. Wu, A Bayesian hierarchical model to detect differentially methylated loci from single nucleotide resolution sequencing data. Nucleic Acids Res, 2014. 42(8): p. e69.

55. Park, Y. and H. Wu, Differential methylation analysis for BS-seq data under general experimental design. Bioinformatics, 2016. 32(10): p. 1446–53.

56. Wu, H., et al., Detection of differentially methylated regions from whole-genome bisulfite sequencing data without replicates. Nucleic Acids Res, 2015. 43(21): p. e141.

57. Heinz, S., et al., Simple combinations of lineage-determining transcription factors prime cis-regulatory elements required for macrophage and B cell identities. Mol Cell, 2010. 38(4): p. 576–89.

58. Yin, Y., et al., Impact of cytosine methylation on DNA binding specificities of human transcription factors. Science, 2017. 356(6337).

59. Yin, J., et al., Alterations in chromatin accessibility and conformation elucidate genetic mechanisms in ASD. medRxiv, 2025.

60. Roadmap Epigenomics, C., et al., Integrative analysis of 111 reference human epigenomes. Nature, 2015. 518(7539): p. 317-30.

61. Ernst, J. and M. Kellis, ChromHMM: automating chromatin-state discovery and characterization. Nat Methods, 2012. 9(3): p. 215–6.

62. Heffel, M.G., et al., Temporally distinct 3D multi-omic dynamics in the developing human brain. Nature, 2024. 635(8038): p. 481-489.

63. Li, M., et al., Integrative functional genomic analysis of human brain development and neuropsychiatric risks. Science, 2018. 362(6420).

64. Trevino, A.E., et al., Chromatin and gene-regulatory dynamics of the developing human cerebral cortex at single-cell resolution. Cell, 2021. 184(19): p. 5053–5069 e23.

65. Li, Y.E., et al., A comparative atlas of single-cell chromatin accessibility in the human brain. Science, 2023. 382(6667): p. eadf7044.

66. Allen Human Brain Atlas: BrainSpan.

67. Allen, W.E., et al., Molecular and spatial signatures of mouse brain aging at single-cell resolution. Cell, 2023. 186(1): p. 194–208 e18.

68. Zemke, N.R., et al., Epigenetic and 3D genome reprogramming during the aging of human hippocampus. bioRxiv, 2024.

69. Chien, J.F., et al., Cell-type-specific effects of age and sex on human cortical neurons. Neuron, 2024. 112(15): p. 2524–2539 e5.

70. Harris, H.K., et al., Disruption of RFX family transcription factors causes autism, attention-deficit/hyperactivity disorder, intellectual disability, and dysregulated behavior. Genet Med, 2021. 23(6): p. 1028–1040.

71. Nativio, R., et al., The chromatin conformation landscape of Alzheimer’s disease. bioRxiv, 2024.

72. Liu, Z., et al., Deciphering aging at three-dimensional genomic resolution. Cell Insight, 2022. 1(3): p. 100034.

73. Shireby, G.L., et al., Recalibrating the epigenetic clock: implications for assessing biological age in the human cortex. Brain, 2020. 143(12): p. 3763–3775.

74. Zhang, Y., et al., Single-cell epigenome analysis reveals age-associated decay of heterochromatin domains in excitatory neurons in the mouse brain. Cell Res, 2022. 32(11): p. 1008–1021.

75. Guo, W., M.Q. Zhang, and H. Wu, Mammalian non-CG methylations are conserved and cell-type specific and may have been involved in the evolution of transposon elements. Sci Rep, 2016. 6: p. 32207.

76. Liu, W., et al., Smccnet 2.0: a comprehensive tool for multi-omics network inference with shiny visualization. BMC Bioinformatics, 2024. 25(1): p. 276.

77. Banerjee-Basu, S. and A. Packer, SFARI Gene: an evolving database for the autism research community. Dis Model Mech, 2010. 3(3-4): p. 133–5.

78. Saitsu, H., et al., De novo mutations in the gene encoding STXBP1 (MUNC18-1) cause early infantile epileptic encephalopathy. Nat Genet, 2008. 40(6): p. 782–8.

79. Barres, R., et al., Non-CpG methylation of the PGC-1alpha promoter through DNMT3B controls mitochondrial density. Cell Metab, 2009. 10(3): p. 189–98.

80. Guo, J.U., et al., Neuronal activity modifies the DNA methylation landscape in the adult brain. Nat Neurosci, 2011. 14(10): p. 1345–51.

81. Stroud, H., et al., Early-Life Gene Expression in Neurons Modulates Lasting Epigenetic States. Cell, 2017. 171(5): p. 1151–1164 e16.

82. Xiong, X., et al., Epigenomic dissection of Alzheimer’s disease pinpoints causal variants and reveals epigenome erosion. Cell, 2023. 186(20): p. 4422–4437 e21.

83. Yang, J.H., et al., Loss of epigenetic information as a cause of mammalian aging. Cell, 2023. 186(2): p. 305–326 e27.

84. Patrick, R., et al., The activity of early-life gene regulatory elements is hijacked in aging through pervasive AP-1-linked chromatin opening. Cell Metab, 2024. 36(8): p. 1858–1881 e23.

85. Zhao, J., et al., A genome-wide profiling of brain DNA hydroxymethylation in Alzheimer’s disease. Alzheimers Dement, 2017. 13(6): p. 674–688.

86. Occean, J.R., et al., Gene body DNA hydroxymethylation restricts the magnitude of transcriptional changes during aging. Nat Commun, 2024. 15(1): p. 6357.

87. Krueger, F. and S.R. Andrews, Bismark: a flexible aligner and methylation caller for Bisulfite-Seq applications. Bioinformatics, 2011. 27(11): p. 1571–2.

88. Krueger, F. Single cell PBAT. 2019; Available from: https://felixkrueger.github.io/Bismark/faq/single_cell_pbat/.

89. Institute, B. *Picard Tools*. Available from: http://broadinstitute.github.io/picard/

90. Danecek, P., et al., Twelve years of SAMtools and BCFtools. Gigascience, 2021. 10(2).

91. Liu, H., et al., DNA methylation atlas of the mouse brain at single-cell resolution. Nature, 2021. 598(7879): p. 120-128.

92. Dobin, A., et al., STAR: ultrafast universal RNA-seq aligner. Bioinformatics, 2013. 29(1): p. 15–21.

93. Liao, Y., G.K. Smyth, and W. Shi, featureCounts: an efficient general purpose program for assigning sequence reads to genomic features. Bioinformatics, 2014. 30(7): p. 923–30.

94. Butler, A., et al., Integrating single-cell transcriptomic data across different conditions, technologies, and species. Nat Biotechnol, 2018. 36(5): p. 411–420.

95. Bakken, T.E., et al., Comparative cellular analysis of motor cortex in human, marmoset and mouse. Nature, 2021. 598(7879): p. 111-119.

96. Hao, Y., et al., Integrated analysis of multimodal single-cell data. Cell, 2021. 184(13): p. 3573–3587 e29.

97. Schliep, K. and K. Hechenbichler Weighted k-Nearest-Neighbor Techniques and Ordinal Classification. 2004. 399, DOI: 10.5282/ubm/epub.1769.

98. Love, M.I., W. Huber, and S. Anders, Moderated estimation of fold change and dispersion for RNA-seq data with DESeq2. Genome Biol, 2014. 15(12): p. 550.

99. Crowell, H.L., et al., *muscat detects subpopulation-specific state transitions from multi- sample multi-condition single-cell transcriptomics data*. Nat Commun, 2020. 11(1): p. 6077.

100. Risso, D., et al., Normalization of RNA-seq data using factor analysis of control genes or samples. Nat Biotechnol, 2014. 32(9): p. 896–902.

101. Van der Auwera, G.A., et al., From FastQ data to high confidence variant calls: the Genome Analysis Toolkit best practices pipeline. Curr Protoc Bioinformatics, 2013. 43(1110): p. 11 10 1-11 10 33.

102. Byrska-Bishop, M., et al., High-coverage whole-genome sequencing of the expanded 1000 Genomes Project cohort including 602 trios. Cell, 2022. 185(18): p. 3426–3440 e19.

103. Price, A.L., et al., Long-range LD can confound genome scans in admixed populations. Am J Hum Genet, 2008. 83(1): p. 132–5; author reply 135-9.

104. Abraham, G., Y. Qiu, and M. Inouye, FlashPCA2: principal component analysis of Biobank-scale genotype datasets. Bioinformatics, 2017. 33(17): p. 2776–2778.

105. Korotkevich, G., et al., Fast gene set enrichment analysis. bioRxiv, 2021.

106. Kolberg, L., et al., gprofiler2 -- an R package for gene list functional enrichment analysis and namespace conversion toolset g:Profiler. F1000Res, 2020. **9**.

107. Young, M.D., et al., Gene ontology analysis for RNA-seq: accounting for selection bias. Genome Biol, 2010. 11(2): p. R14.

108. Gel, B., et al., regioneR: an R/Bioconductor package for the association analysis of genomic regions based on permutation tests. Bioinformatics, 2016. 32(2): p. 289–91.

109. Malinverni, R., et al., regioneReloaded: evaluating the association of multiple genomic region sets. Bioinformatics, 2023. 39(11).

110. Kuleshov, M.V., et al., Enrichr: a comprehensive gene set enrichment analysis web server 2016 update. Nucleic Acids Res, 2016. 44(W1): p. W90–7.

111. Gene Ontology, C., et al., The Gene Ontology knowledgebase in 2023. Genetics, 2023. 224(1).

112. Sarda-Espinosa, A. dtwclust: Time Series Clustering Along with Optimizations for the Dynamic Time Warping Distance. 2024; Available from: https://cran.r-project.org/package=dtwclust.

113. Zhou, W., et al., SeSAMe: reducing artifactual detection of DNA methylation by Infinium BeadChips in genomic deletions. Nucleic Acids Res, 2018. 46(20): p. e123.

