## Supplemental Figures for "A Single-Cell Atlas of DNA Methylation in Autism Spectrum Disorder Reveals Distinct Regulatory and Aging Signatures"

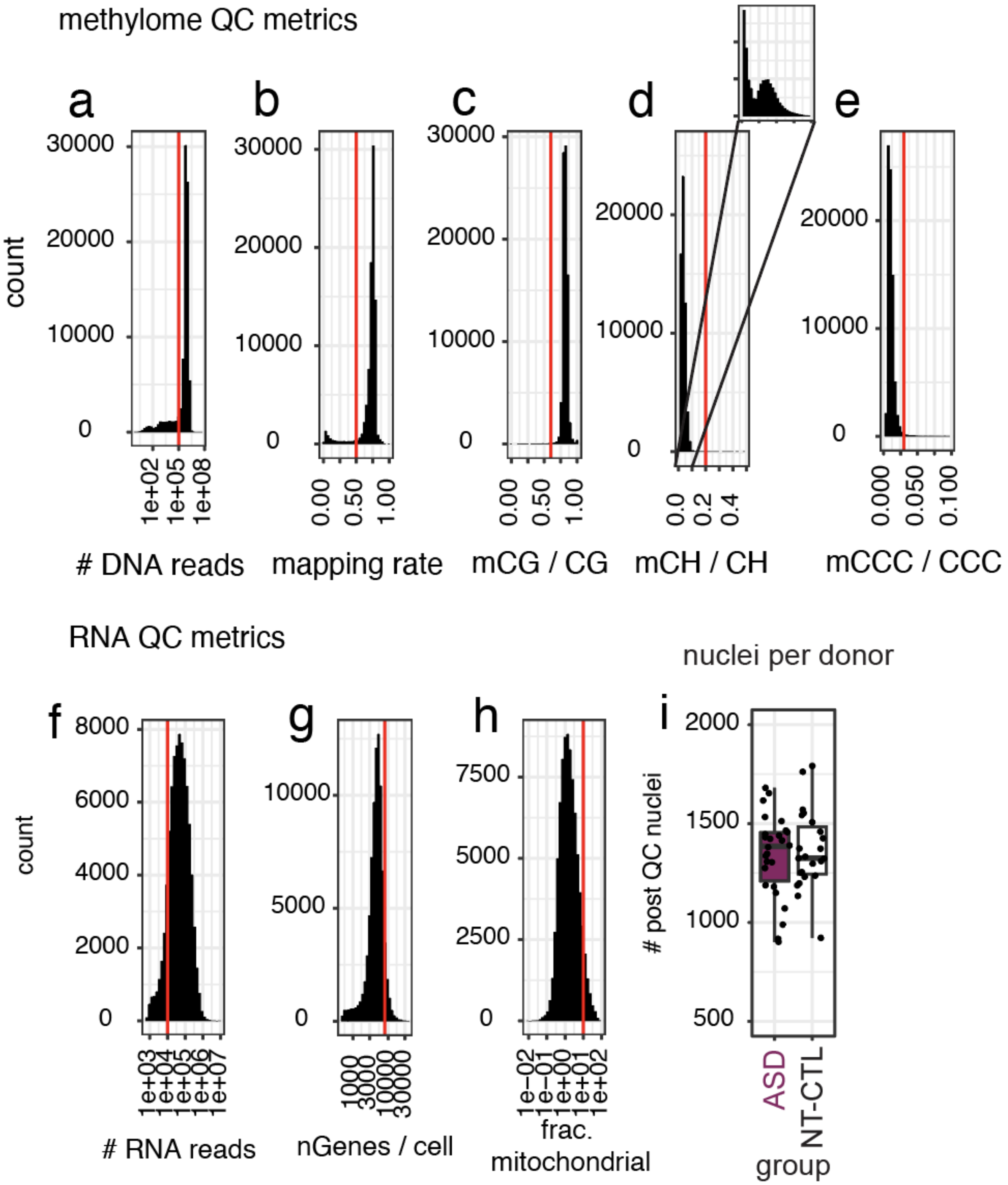


**Figure S1. Nuclei-level quality control metrics, by modality.** Methylome reads per cell, after mCT filtering (a), mapping rate (b) and mCG (c), mCH (d) and mCCC (e) rates. RNA reads per cell (f), genes per cell (g) and mitochrondrial fraction of transcripts (h). In a – h, filtering thresholds are indicated on each plot. (i) number of nuclei recovered per donor, grouped by diagnosis, after methylome QC filtering.

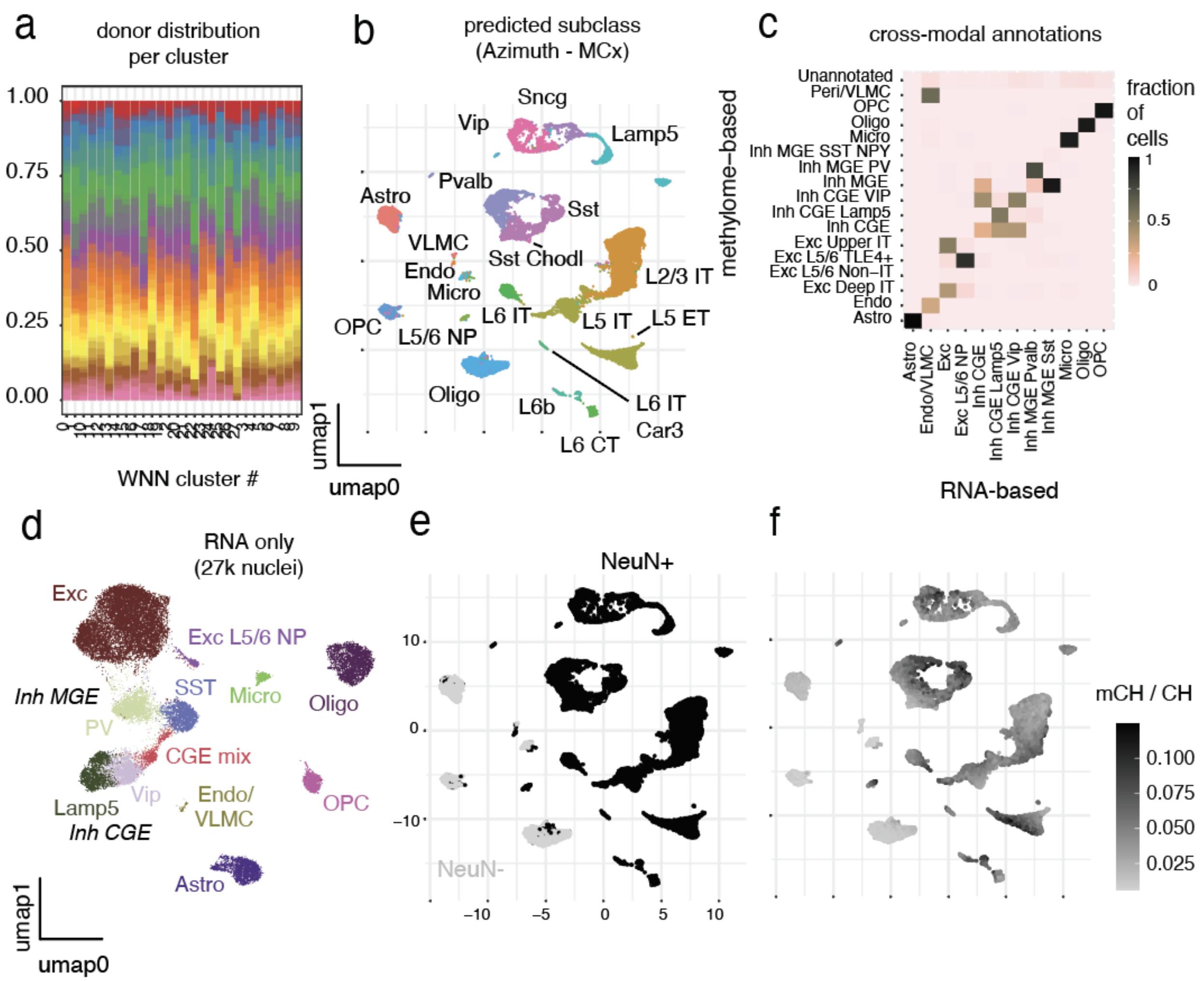


**Figure S2. Cell type annotation.** Donor contribution to each WNN cluster (a). Predicted cell type labels, based on motor cortex reference data from (Bakken, Jorstad et al. 2022) (b). Concordance of annotation across modalities (c). UMAP representation data based only on gene expression (d). UMAP representation of WNN data, colored by NeuN staining (e) and absolute mCH levels (f).

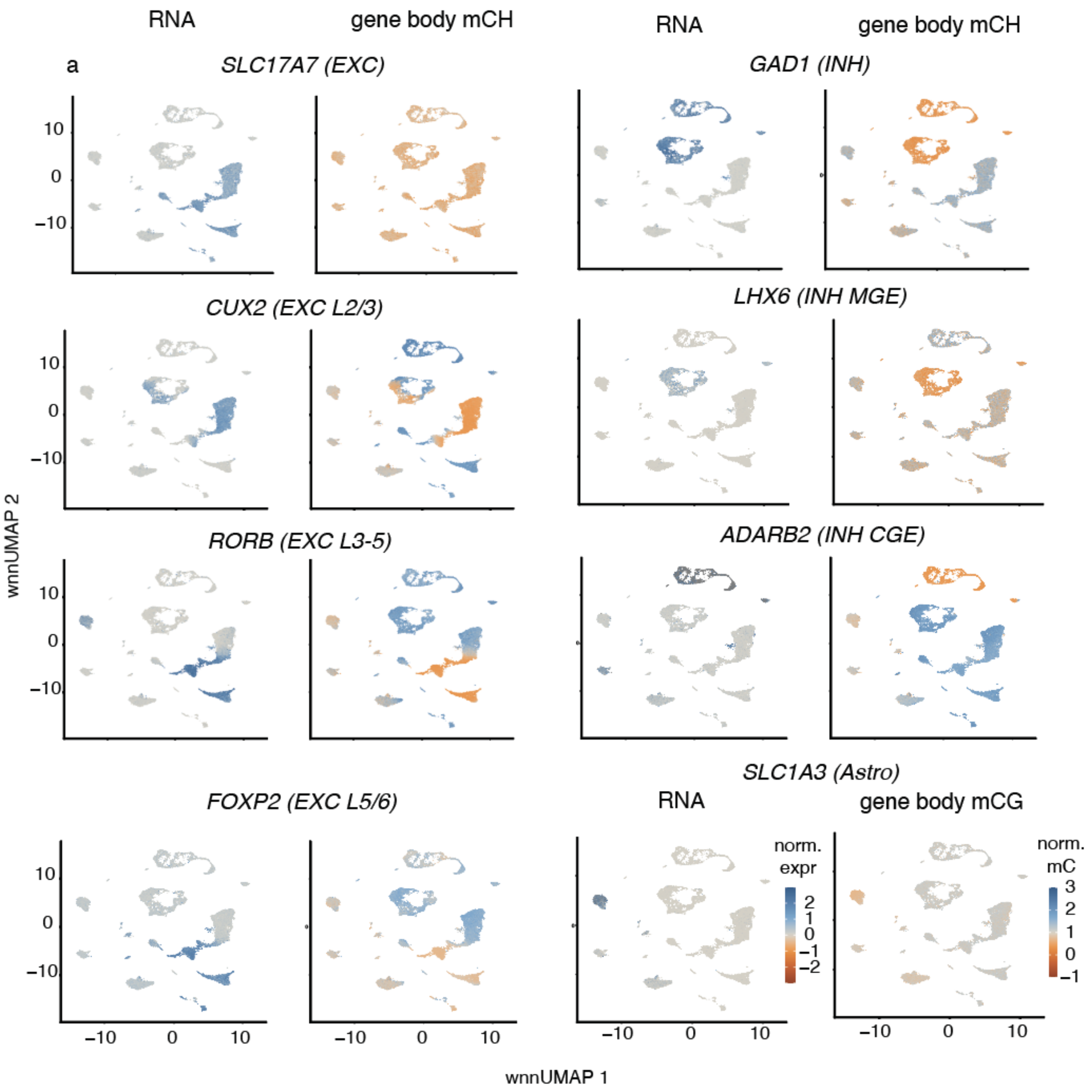


**Figure S3. Example marker gene expression and (normalized) gene body mCH and mCG methylation.**

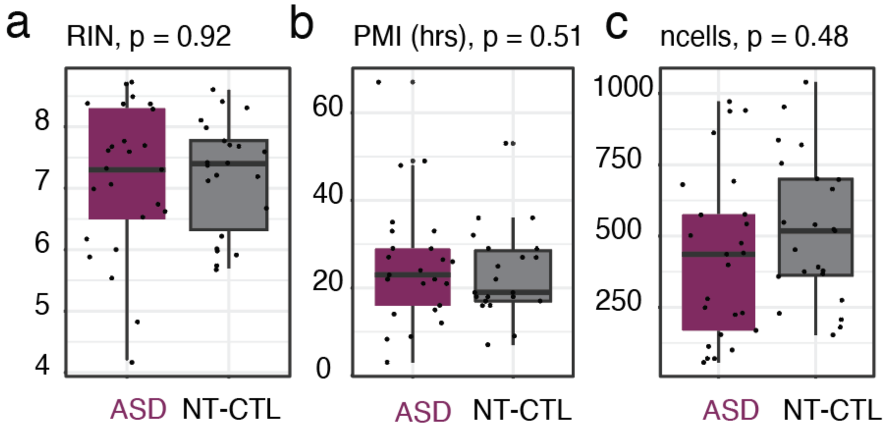


**Figure S4. Donor-level RNA QC**. Donor RNA integrity number (RIN, a), post-mortem interval (PMI, b) and number of post-QC transcriptome cells (c).

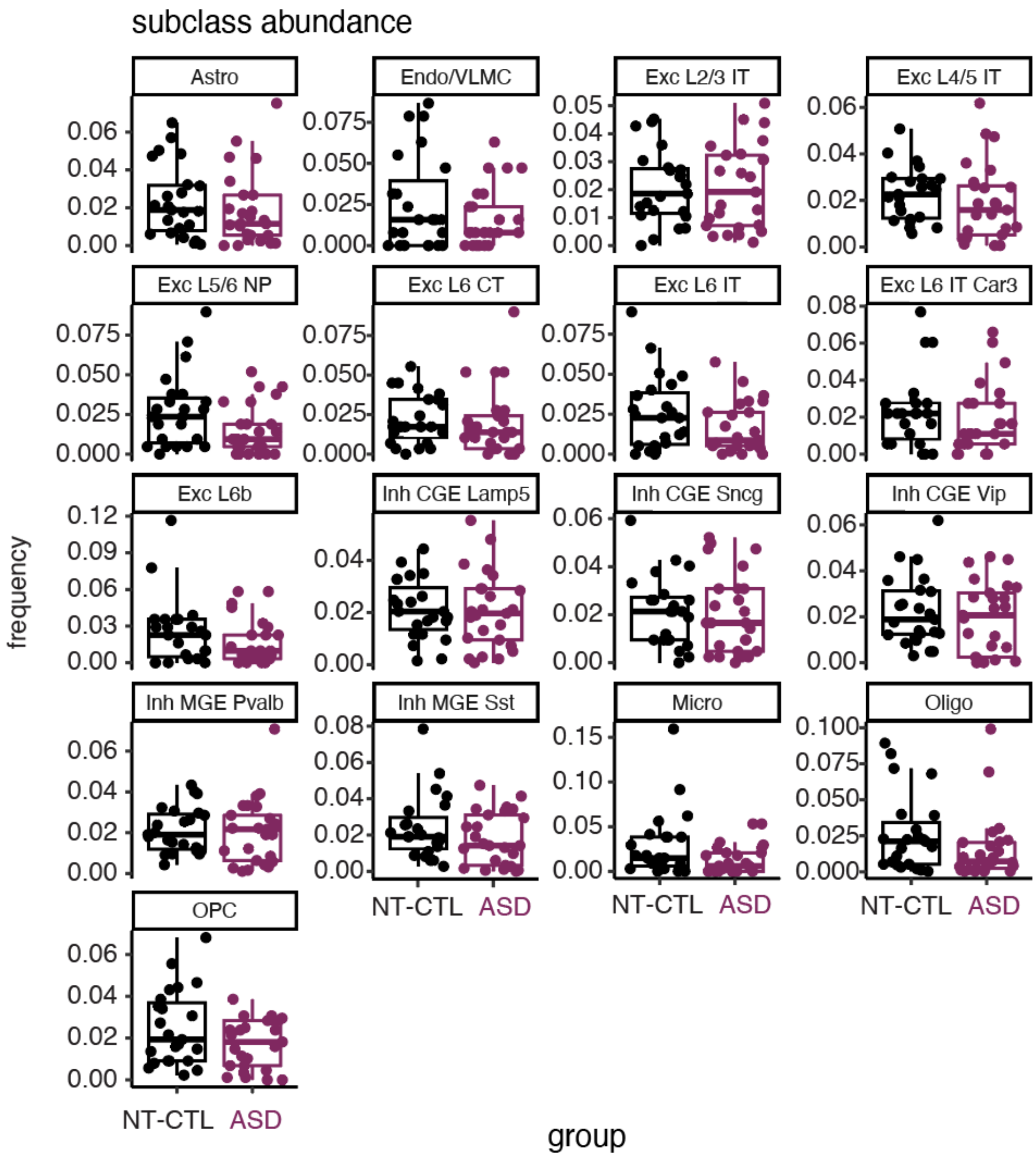


**Figure S5. Cell type composition analysis in ASD**. Donor contribution to each subclass, split by diagnosis. No differences significant (padj < 0.1) after FDR correction.

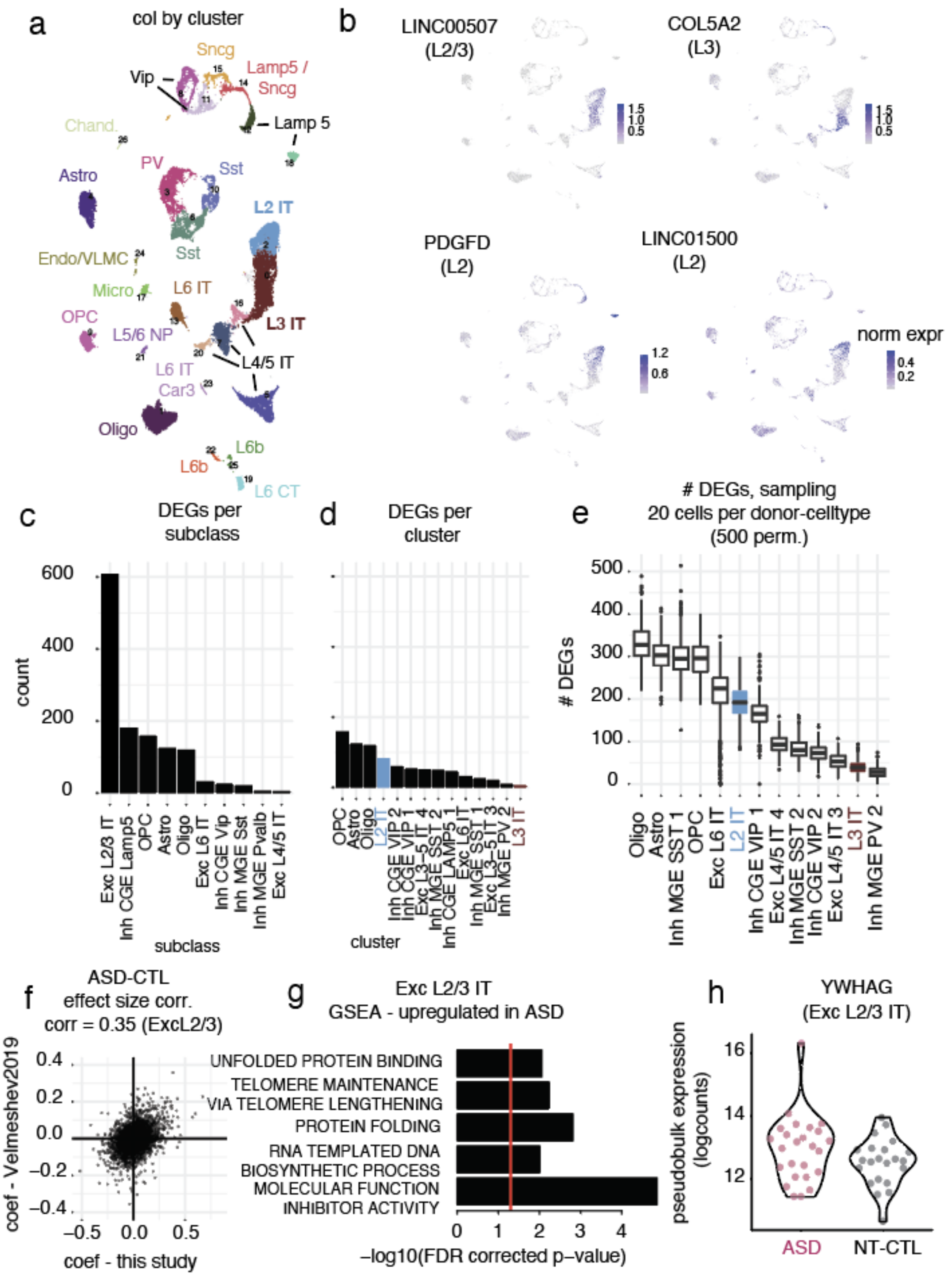


**Figure S6. Differential gene expression analysis**. (a) WNN umap, colored by cluster. (b) Expression of layer II and layer III cortical markers identified in (Hodge et al., 2019). (c) Number of differentially expressed genes (DEGs, padj < 0.05) identified at two levels of annotation: subclass (c) and cluster (d). (e) Number of DEGs identified when randomly sampling 20 cells per donor-celltype across 500 permutations. (f) ASD-CTL effect size correlation in Exc L2/3 IT neurons in this dataset and in (Velmeshev et al., 2019). *In this analysis, effect sizes from the data in this study are reported after processing in the same framework used in (Velmeshev et al., 2019) (see Methods). (g) Gene set enrichment analysis (GSEA) of ASD-upregulated transcripts in Exc L2/3 IT neurons. (h) pseudobulk expression of an example DEG.

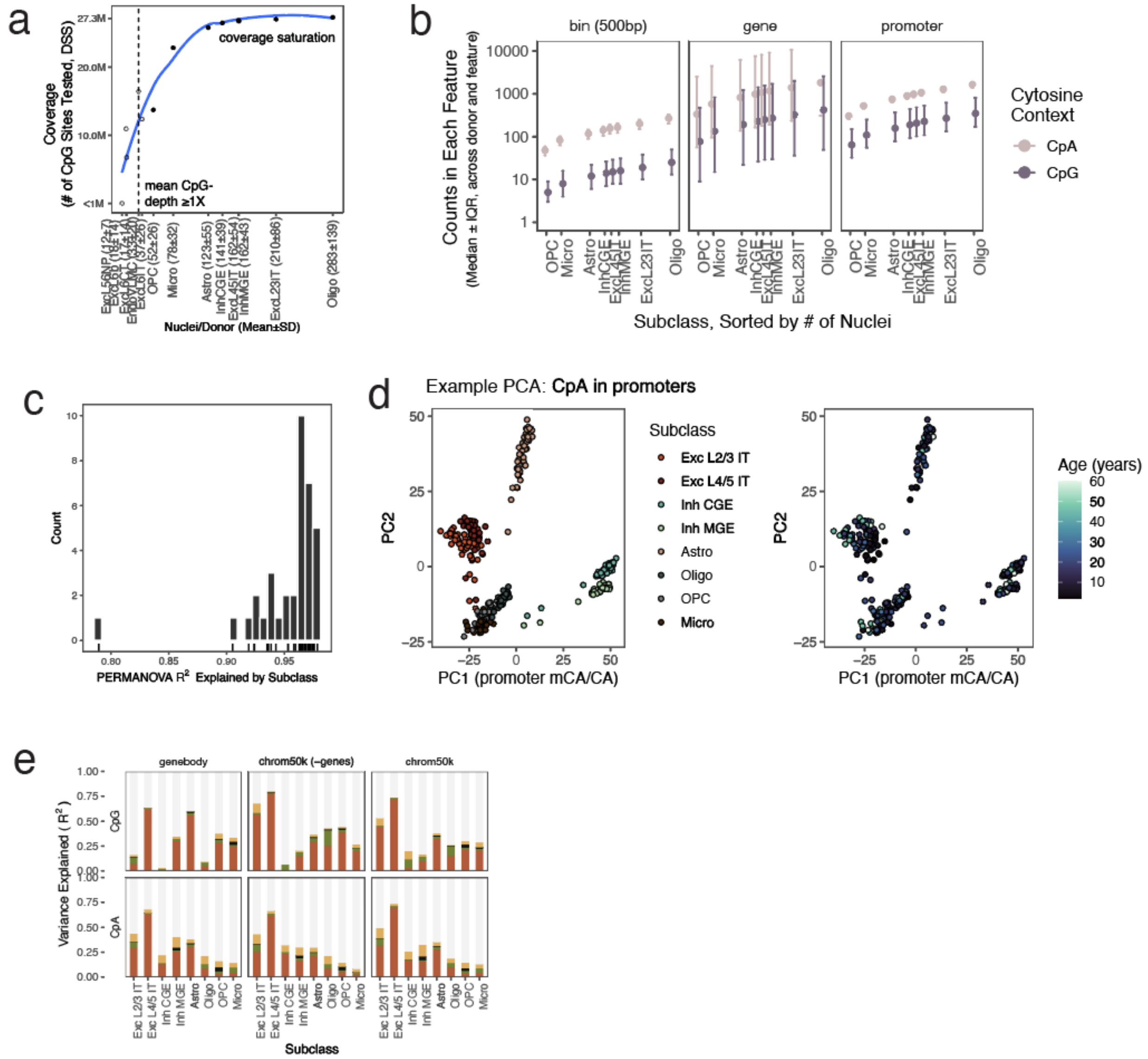


**Fig S7. Coverage levels and sources of variance in donor-by-subclass methylome pseudobulks.** (a) Number of single-cytosines included in DSS test (observed in ≥75% of donors, not overlapping ENCODE exclusion list), plotted by subclass abundance. Subclasses tested (closed circles) had ≥1X mean sequencing depth per CpG-site (dashed line) and were observed within all n=49 donors. (b) Summary of pseudobulk-level counts within pre-specified features, taking the median across donors and across features. Specifically, we tabulated the total counts $c_{isf}$ of donor *i*, in subclass *s*, overlapping prespecified feature *f*. For fixed *s,* the error bars show median ± IQR values of $c_{if}$. (c) Variance explained by subclass label estimated from applying PERMANOVA to principal components derived from different methylome features (e.g., genes; 100kb bins with less than 20% base pair overlap with genes [“-genes”]) and cytosine contexts (CpG-, CpA-, CpH-). The lowest smallest r2 is in CH-promoters. (d) Example of principal components analysis (PCA) input into PERMANOVA. Each point is a pseudobulk, shown colored by subclass then by donor age. (e) Percent variance explained by known donor covariates, with different feature sets.

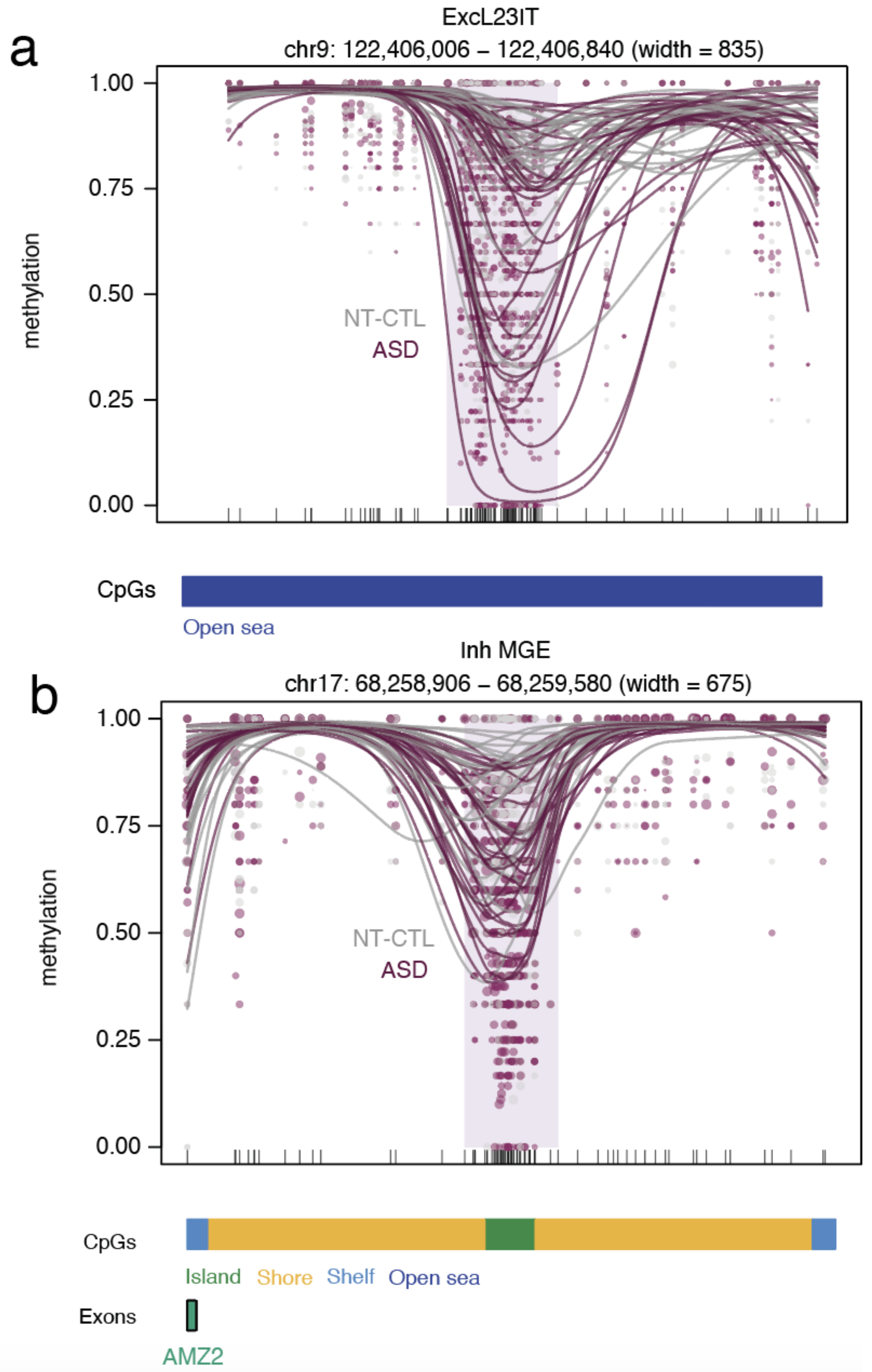


**Figure S8. Example *de novo* ASD DMRs.** Plots generated using the plotDMR function from dmrseq (Korthauer, Chakraborty, Benjamini, & Irizarry, 2019). Individual points reflect methylation estimates at individual cytosines in pseudobulk data. Point size reflects coverage and each line represents the smoothed, average methylation per donor.

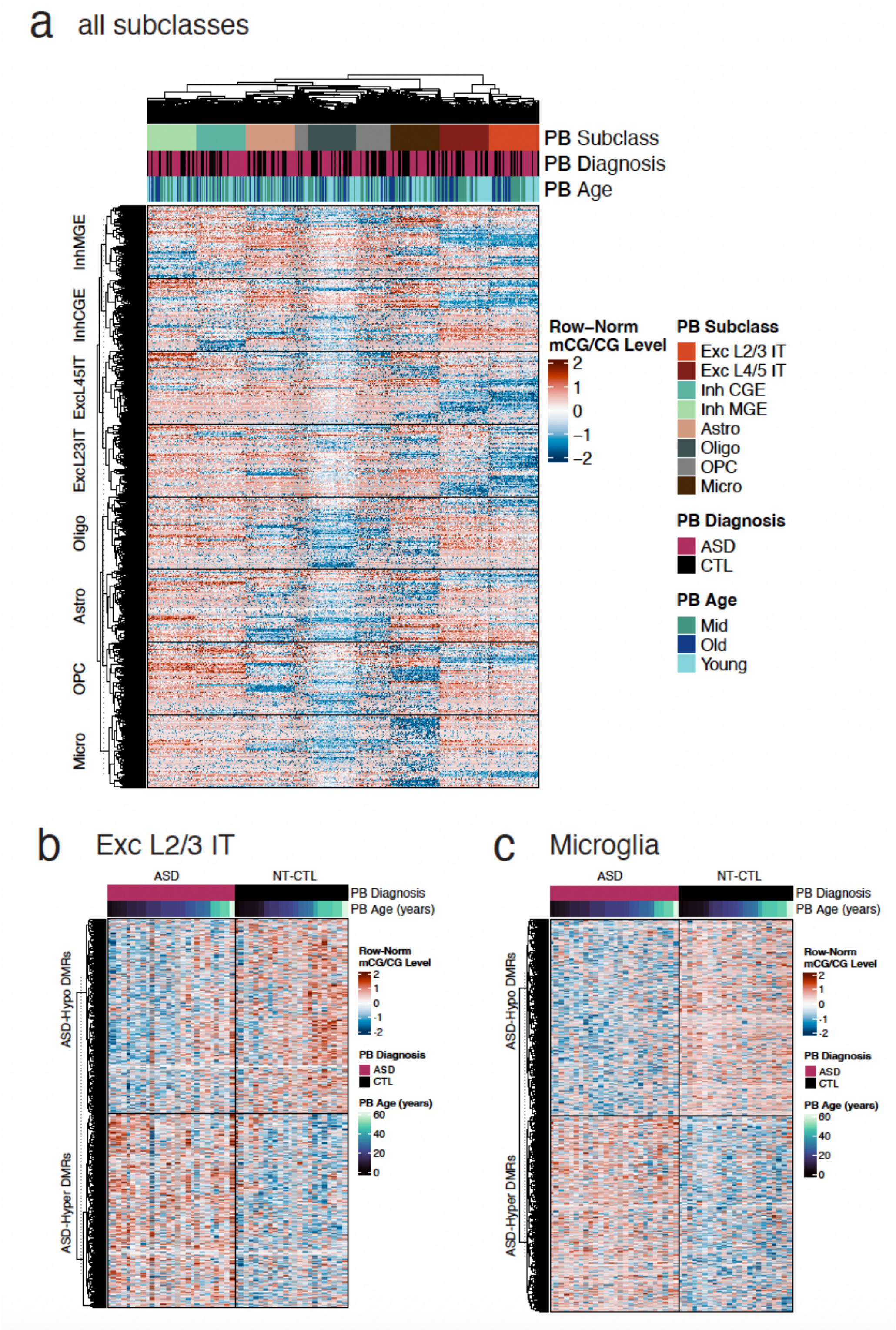


**Figure S9. Normalized mCG/CG in example ASD-DMRs.** (a) Normalized CG-methylation fraction in randomly selected ASD-DMR regions (rows, 100 DMRs from each subclass-by-direction [hypo/hyper] combination) from each donor-by-subclass pseudobulk (columns; PB = pseudobulk). The methylation fraction is quantile-normalized in each row, such that the pseudobulk with highest methylation fraction in the region have positive values (red) and lowest methylation fraction have negative values (blue). Rows and columns are ordered by their similarity from hierarchical clustering on Euclidean distance, resulting in unsupervised clustering by cell type. Within excitatory cell types, the pseudobulks further notably cluster by age. (b) Example of 500 ASD-hypo- and 500 ASD-hyper L2/3 DMRs by pseudobulk methylation fractions for L2/3 and (c) repeated for microglia, with pseudobulk columns sorted by diagnosis and age.

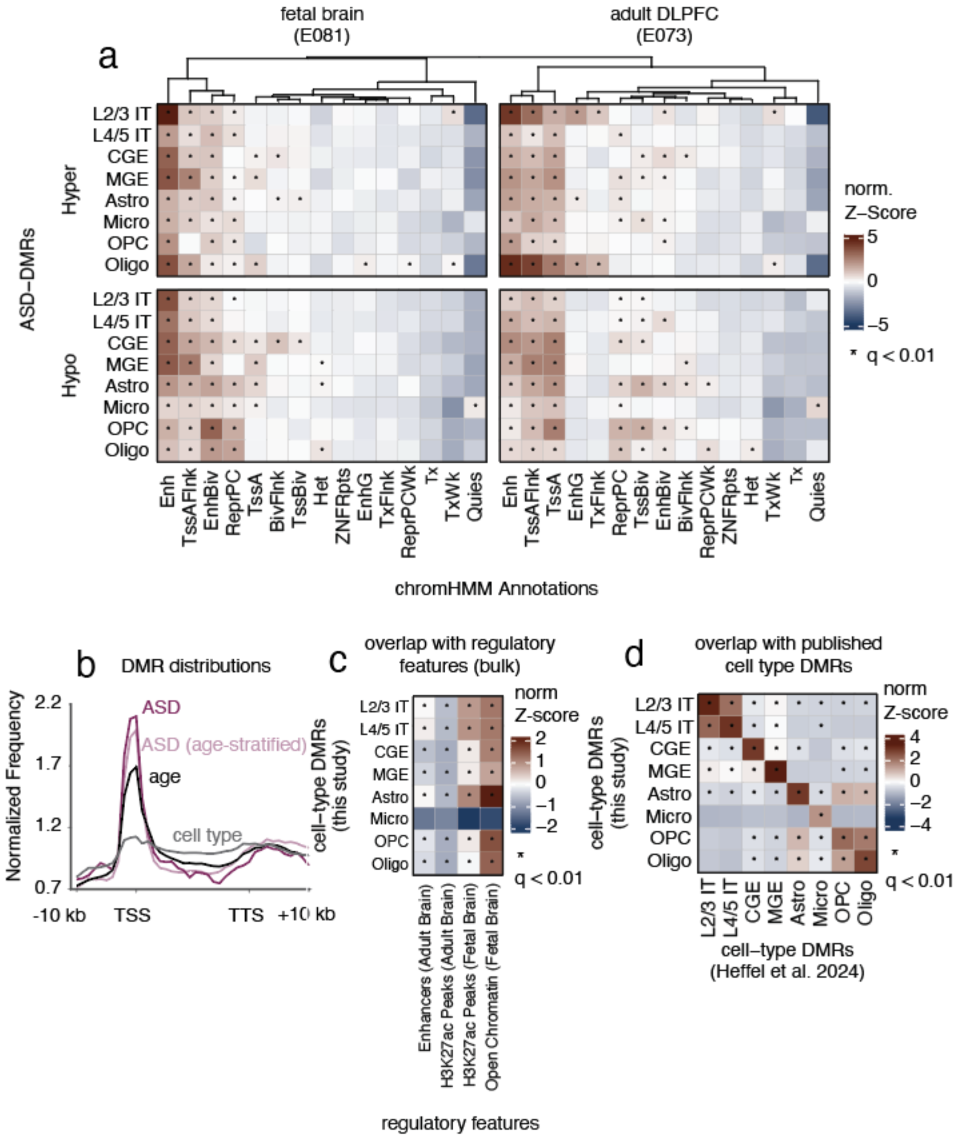


**Figure S10. Alternative ASD-DMR annotation and alternative DMR sets** (a) ChromHMM annotation of ASD-DMRs in two sample types; one, from adult dorsolateral prefrontal cortex and the other from fetal brain. (b) Distribution of multiple DMR sets (diagnosis, age, and cell type) around transcriptional start sites (TSSs) and gene bodies. Overlap of cell type DMRs with known regulatory regions (c) and previously published cell type DMRs (d).

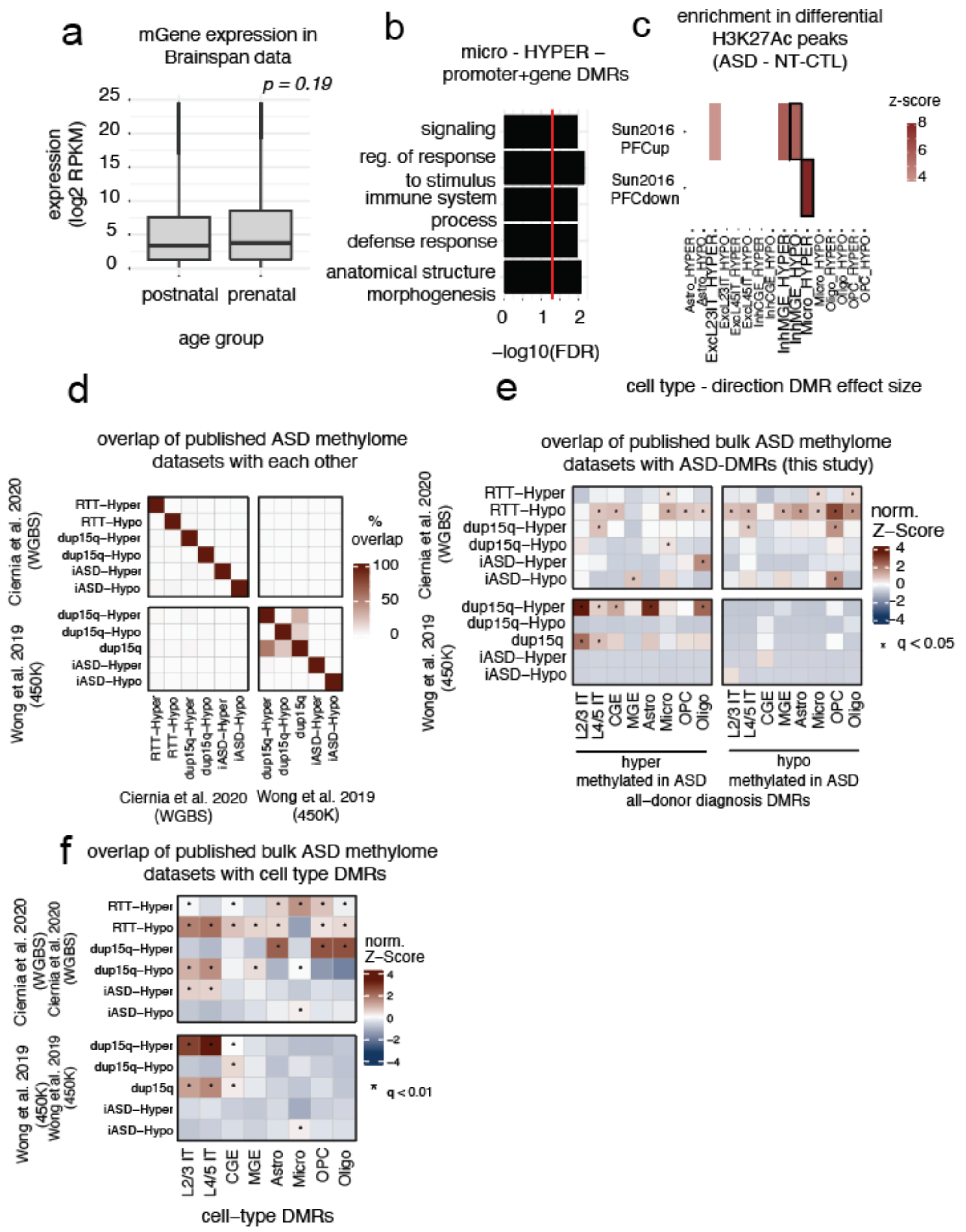


**Figure S11. Interpreting ASD-DMRs in the context of external datasets and gene expression**. (a) Expression of genes implicated by DMRs (mGene) in the BrainSpan database("Allen Human Brain Atlas: BrainSpan,"). Pvalue is the result of a paired t-test comparing gene-wise expression in pre- and post-natal samples. (b) Gene set enrichment analysis of genes implicated by ASD-DMRs that are hypermethylated in microglia. (c) Enrichment of ASD-DMRs with differential H3K27Ac ASD peaks  (Sun et al., 2016). (d) Percent overlap of DMRs identified in published analyses of ASD methylome (Vogel Ciernia et al., 2020; Wong et al., 2019). (e) Heatmap depicting the overlap between cell type DMRs and ASD-DMRs identified in bulk analysis. (f) Heatmap depicting the overlap between ASD DMRs (this study) and those identified in bulk analysis.

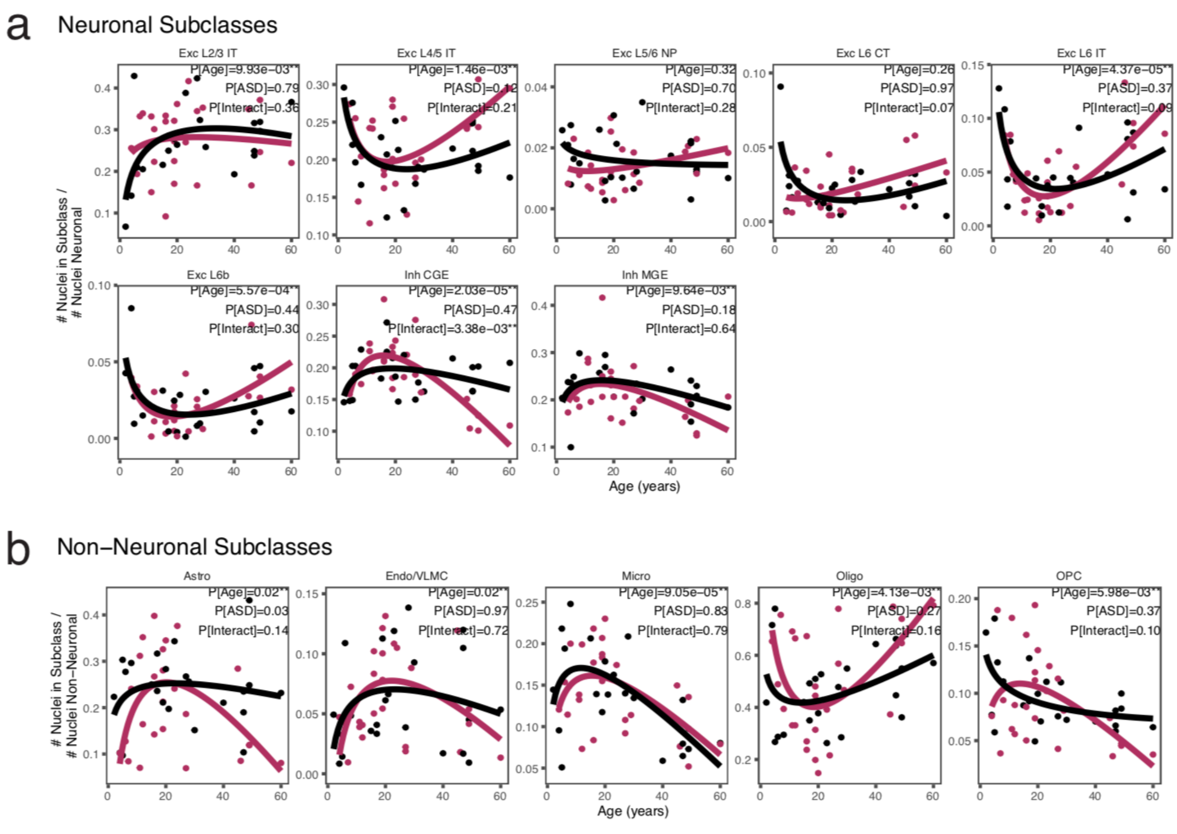


**Figure S12. Cell type composition analysis over age and between diagnostic groups**.

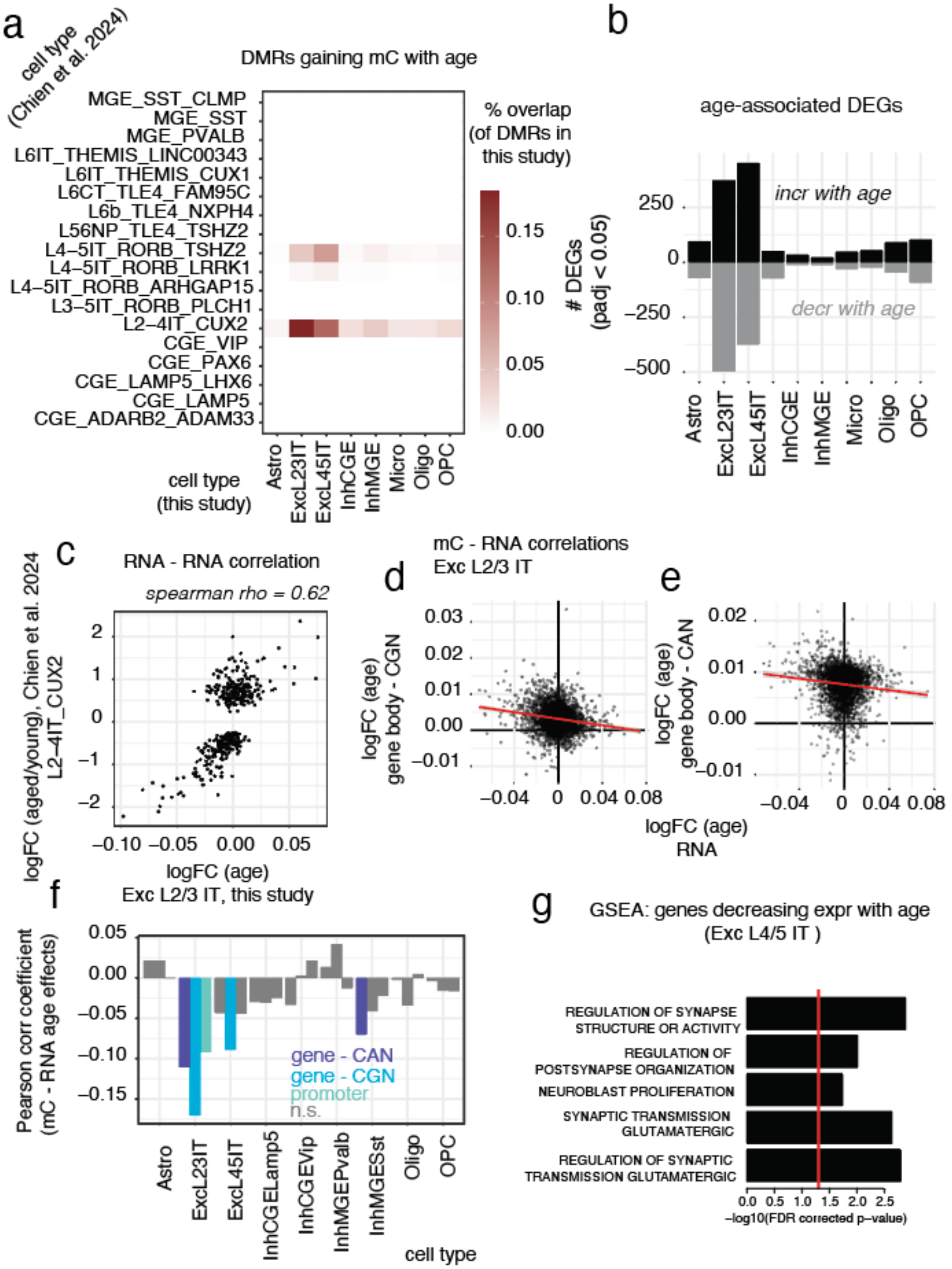


**Figure S13. Additional analyses on age-DMRs and age-DEGs.** (a) Overlap of age-DMRs identified in this study and (Chien et al., 2024) (b) Summary of age-DEGs (c) Correlation of gene expression effect size between age-DEGs in this study and (Chien et al., 2024). Examples from gene body mCG (d) and mCH (e); summary in (f). (g) Gene set enrichment analysis of genes downregulated in excitatory L4/5 IT neurons.

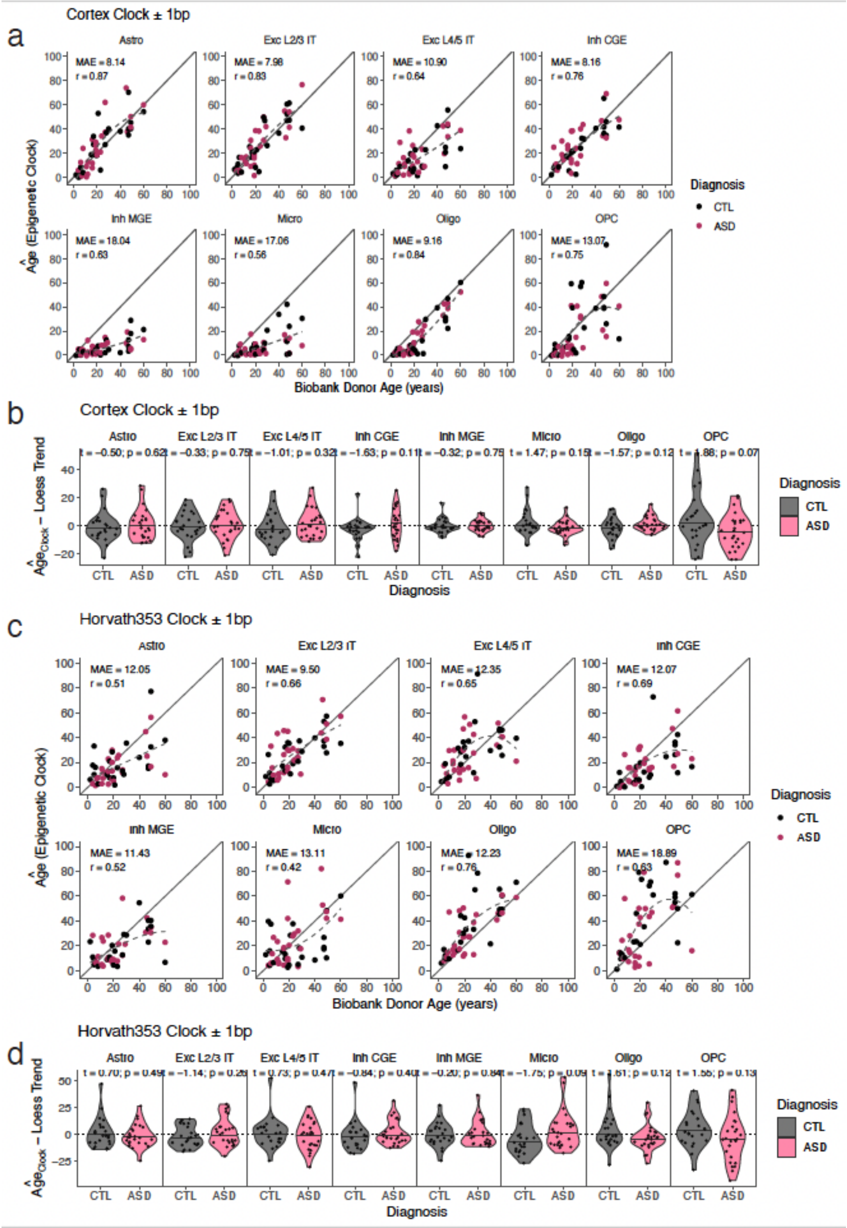

**Fig S14**. **Methylation clock performance in single-nuclei data.** (a) Performance of the cortex epigenetic clock by subclass, as illustrated by a scatterplot of the clock-estimated epigenetic age versus biobank recorded age. Each point represents one donor. The Mean Absolute Error (MAE = average absolute value of predicted minus true age; lower values indicate better performance) and Spearman correlation (between clock and true age; higher values indicate better performance) is shown for each celltype, across ASD and NT-CTL samples. The solid diagonal line indicates the y=x line of exact fit, whereas the dashed line indicates a loess regression (R stats::loess, span = 5) showing the mean trend between the predicted and true ages; this fit is analogous to a calibration line between true age and clock age. A given donor's epigenetic age appearing above this loess line would suggest advanced epigenetic age beyond the mean predictions for donors of that age range. (b) We computed each donor's predicted epigenetic age minus the loess trend mean for calibration, then compared this value by diagnosis (t-test) in each group. (c) Epigenetic predictions and (d) between-diagnosis comparison for the Horvath 353-probe clock.

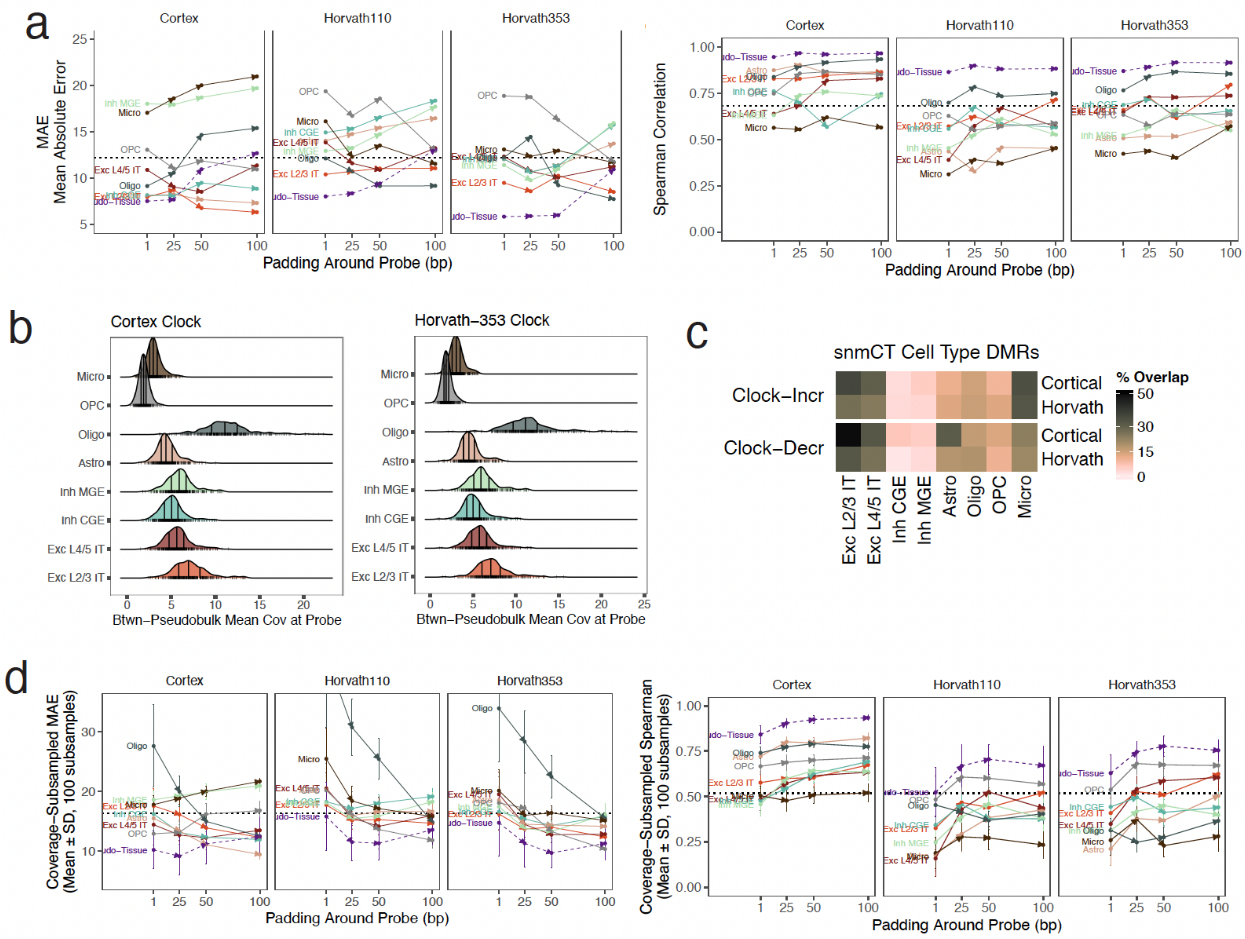


**Figure S15**. **Robustness of epigenetic age clock performance by cell type**. We constructed a high coverage “pseudo-tissue” by calculating a given donor’s methylation fraction from all 8 subclasses together; as each clock is validated in bulk data, this composite thus represents a performance bound due to technical effects of transferring the array-based technology to our sequencing-based measurements. Second, we hypothesized that given the spatial correlation of CG-methylation (55% in cortex, 81% in Horvath-343)—that padding fractional methylation calculations to include sites within a ±w base window of each probe may reduce noise by potentially increasing effective sequencing depth. (a) Clock performance (MAE, Spearman correlation) by subclass, for varying “padding” levels. The pseudo-tissue performance had similar, marginally better performance than the best performing subclasses for the cortex clock; however the Horvath clock performed noticeably better in the pseudo-tissue. Padding up to ±25bp increased performance for most cell-types, suggesting that this relatively simple method may aid in the development of clocks for single-nucleus data; however, padding still did not resolve the between-cell type differential performances. (b) The distribution of counts overlapping each clock probe, by cell type. Vertical lines on each distribution indicate median and interquantile range. (c) The fraction of clock probes covered by cell type DMRs. (d) Each pseudobulk and pseudo-tissue were sampled with replacement to the same mean coverage at each probe site. This process was repeated 100 times. The mean and sd across the sampling iterations are shown as error bars. Subsampling generally decreased performance for all cell types, but their relative rank of performance stays generally similar for all cell types, with the notable exception of Oligodendrocytes. The bulk “pseudo-tissue” retains the highest performance even with subsampling.

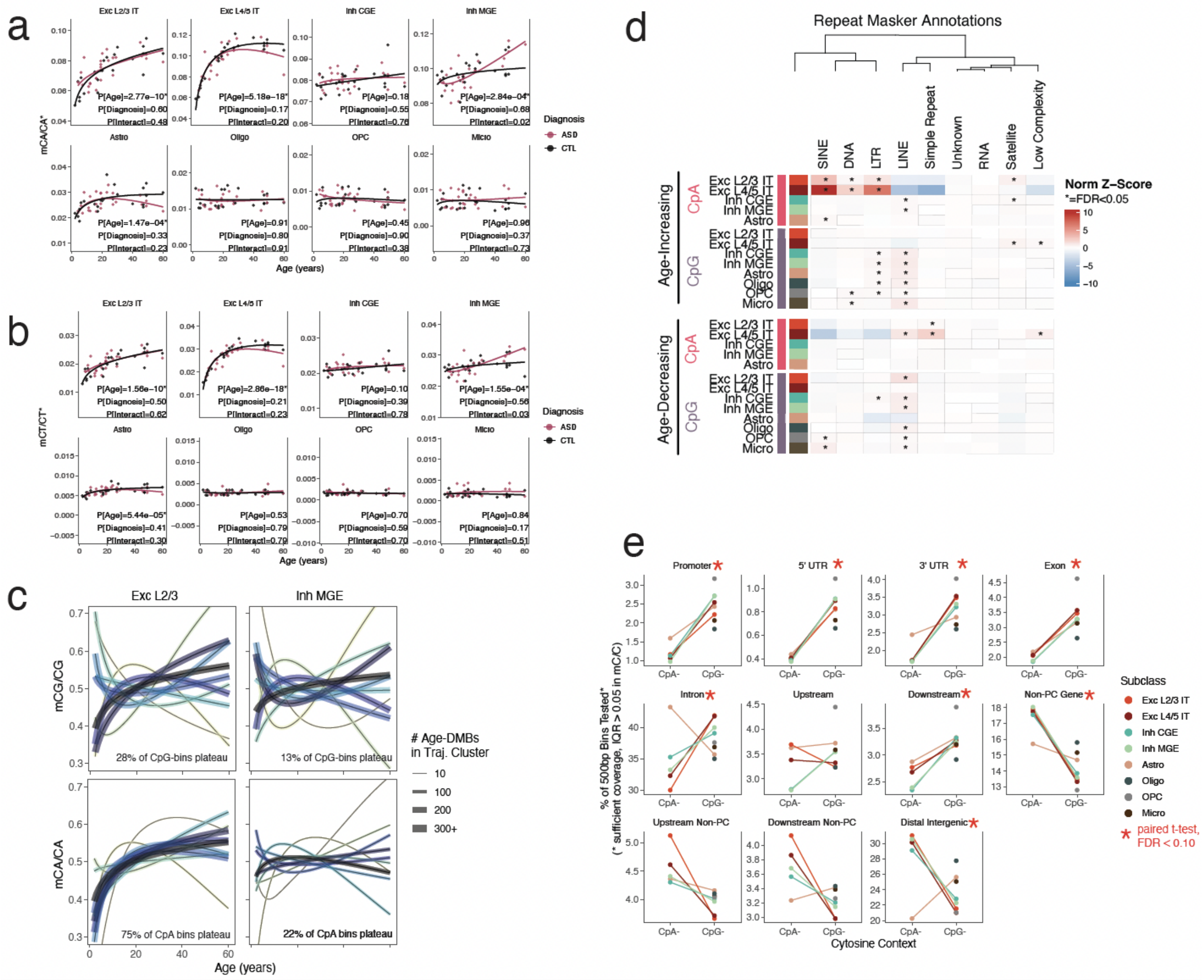


**Fig S16. Non-CG methylation over age and across cell types**. Global methylation levels plotted by age for (a) CpA- and (b) CpT- sequence contexts, minus mCCC/CCC as a proxy for potential bisulfite conversion levels. We detected significant changes with global levels with age in Exc L2/3, Exc L4/5, Inh MGE, Astro (asterisk indicates q-value < 0.05), but no diagnosis nor age-by-diagnosis interactions (although Inh MGE had a notable interaction P < 0.05). (c) Clustering was applied age-DMBs to characterize common patterns of change; each line depicts the mean pattern by cluster, with the line width indicating the number of bins the cluster. The most common pattern for Exc L2/3 and L4/5 were increases in CpG- and CpA-methylation that plateau before age 20-30. (d) Enrichments of RepeatMasker elements (transposable elements) with age-DMBs. (e) Overlap of between-donor variable 500bp bins (IQR>0.05) with GENCODE features, by sequence context. Asterisks indicate significant mean differences in overlap between CpA- and CpG- contexts.

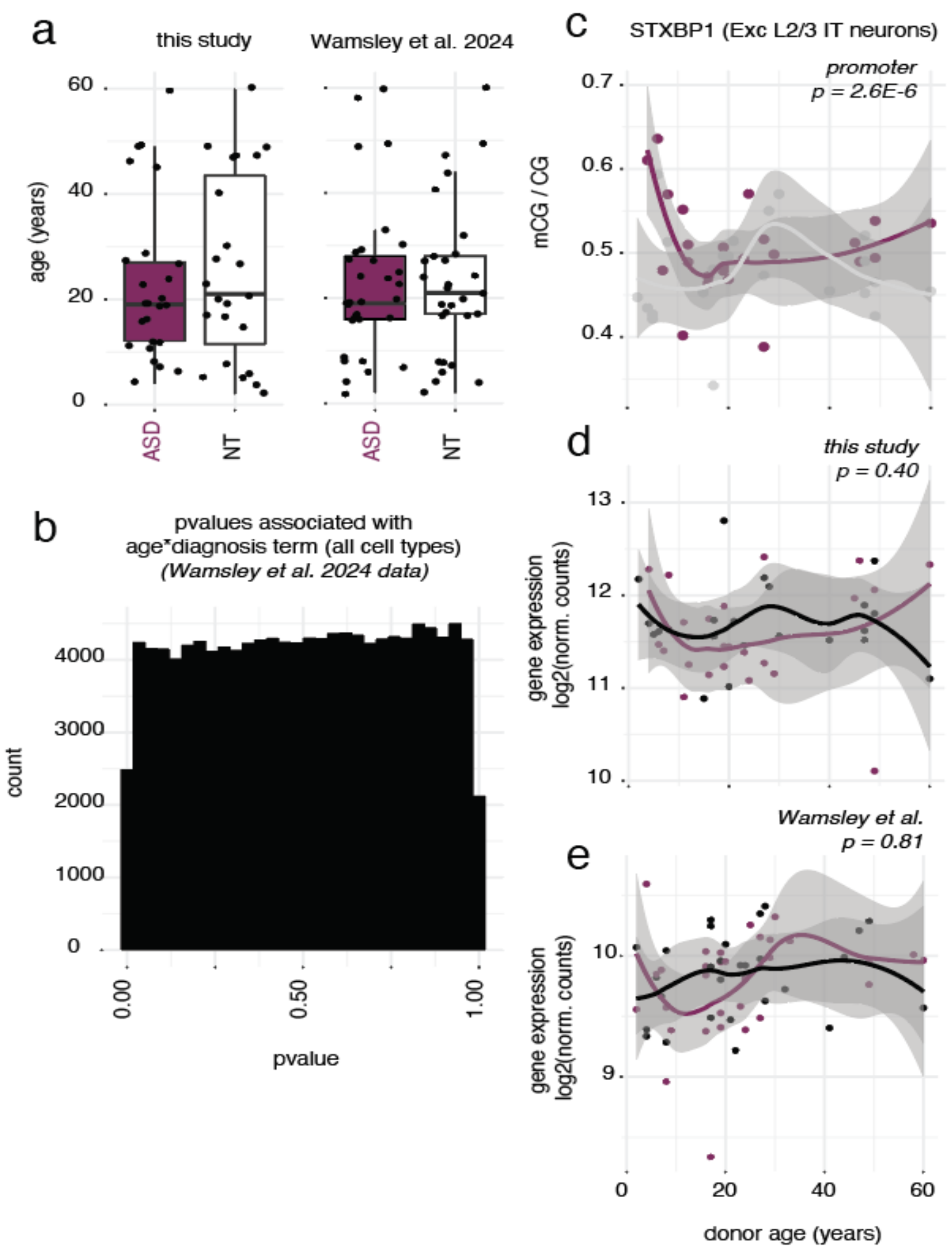


**Figure S17. Probing age-dependent diagnosis effects in scRNAseq data** (a) Age distribution of donors in this dataset and in (Wamsley et al., 2024) (b) Pvalue distribution for age*diagnosis interaction term in DESeq2 analysis of data from (Wamsley et al., 2024) across all cell types. Promoter methylation of top interaction feature *STXBP1* in Exc L2/3 IT neurons (c) and corresponding gene expression in 2 datasets (d, e).

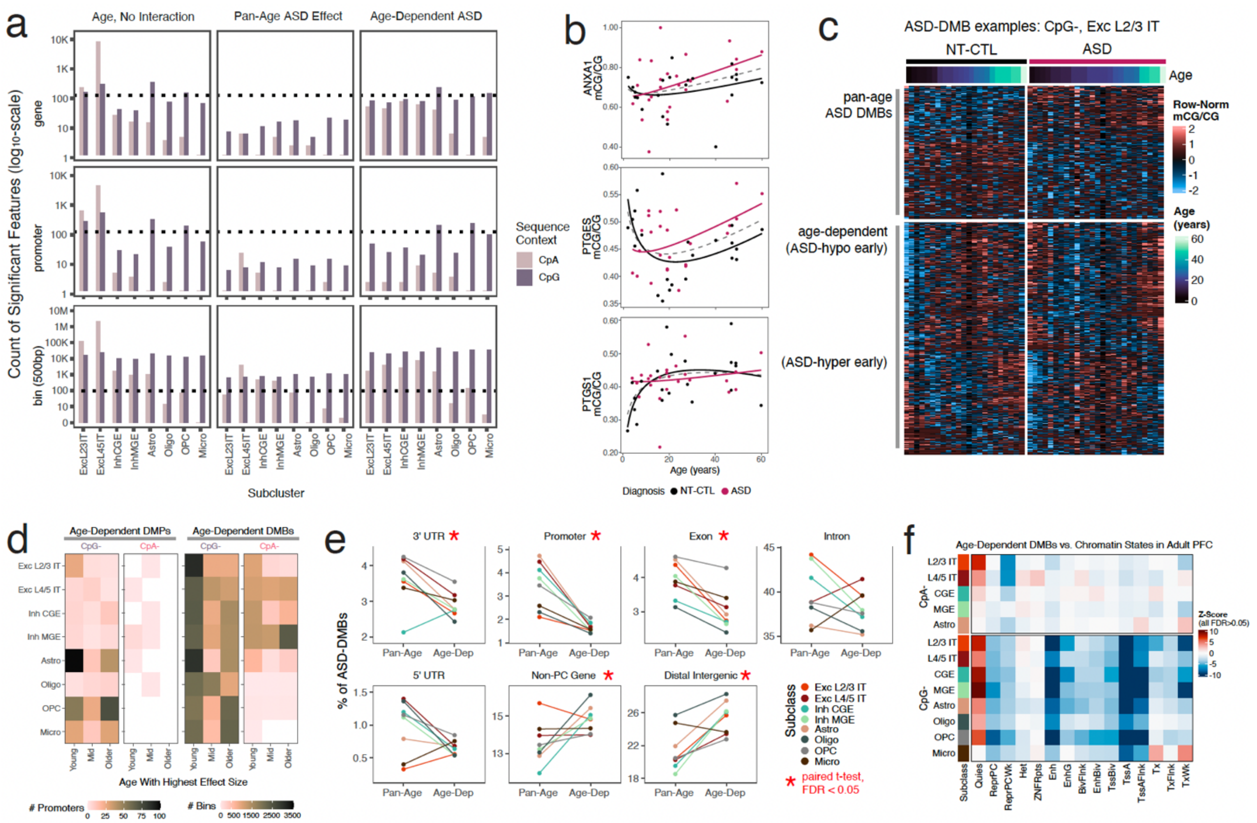


**Fig S18. Properties of age-dependent ASD methylome features.** (a) Counts of significant features (DMB, DMG, DMP) on $\log_{10}$scale, by subclass and cytosine context. (b) Example promoter mCG/CG in Exc L2/3 neurons, from the fatty metabolism pathway enriched for significant age-dependent effects (GSEA eFDR < 0.05). (c) Normalized methylation fraction for the top 200 pan-age and top 500 age-dependent ASD CG-bins (rows) by samples (columns) detected in Excitatory L2/3 IT neurons. DMBs were ranked by likelihood ratio p-value (model with interaction compared to reduced model without interaction). Each bin was quantile-normalized across columns, and were sorted using unsupervised hierarchical clustering. (e) Percent of age-dependent and pan-age DMBs assigned to each genomic feature. Significant differences between age-dependent and pan-age DMBs indicated by asterisk. (e) ChromHMM enrichments for age-dependent ASD effects in DMBs, split by context and cell type. No enrichments were statistically significant following multiple testing correction.

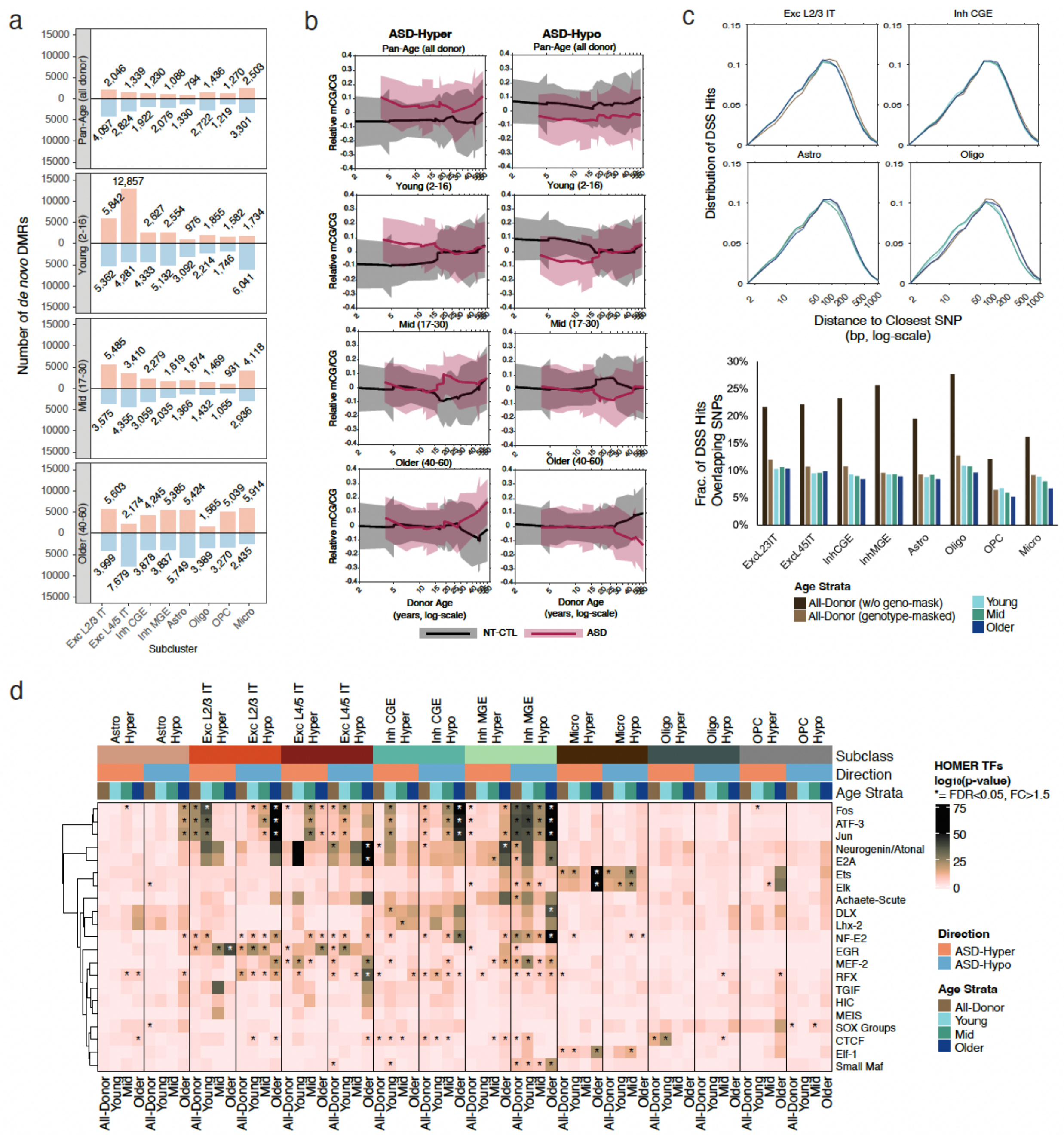

**Figure S19. Supplemental analysis of age-stratified *de novo* DMRs.** (a) Number of de novo ASD-DMRs detected in all-donor (pan-age) analysis and in each of the three age strata. (b) Composite of *de novo* DMRs called in Exc L4/5 in all-donor and age strata to visualize effect sizes. The raw methylation fractions in each pseudobulk were extracted over each ASD-DMR set, mean-centered, then averaged to form the composite line representing the methylation levels across ASD-DMRs (mean ± standard deviation). (c) Histogram depicting distance between ASD DMRs (all-donor and age-stratified) and donor-specific genetic variants. (d) More comprehensive set of TF motif enrichments for age-stratified *de novo* ASD-DMRs, including TF families beyond those reported in the pan-age (all-donor) analysis.

References


Allen Human Brain Atlas: BrainSpan.

Chien, J. F., Liu, H., Wang, B. A., Luo, C., Bartlett, A., Castanon, R., . . . Mukamel, E. A. (2024). Cell-type-specific effects of age and sex on human cortical neurons. *Neuron, 112*(15), 2524-2539 e2525. doi:10.1016/j.neuron.2024.05.013

Hodge, R. D., Bakken, T. E., Miller, J. A., Smith, K. A., Barkan, E. R., Graybuck, L. T., . . . Lein, E. S. (2019). Conserved cell types with divergent features in human versus mouse cortex. *Nature, 573*(7772), 61-68. doi:10.1038/s41586-019-1506-7

Korthauer, K., Chakraborty, S., Benjamini, Y., & Irizarry, R. A. (2019). Detection and accurate false discovery rate control of differentially methylated regions from whole genome bisulfite sequencing. *Biostatistics, 20*(3), 367-383. doi:10.1093/biostatistics/kxy007

Sun, W., Poschmann, J., Cruz-Herrera Del Rosario, R., Parikshak, N. N., Hajan, H. S., Kumar, V., . . . Prabhakar, S. (2016). Histone Acetylome-wide Association Study of Autism Spectrum Disorder. *Cell, 167*(5), 1385-1397 e1311. doi:10.1016/j.cell.2016.10.031

Velmeshev, D., Schirmer, L., Jung, D., Haeussler, M., Perez, Y., Mayer, S., . . . Kriegstein, A. R. (2019). Single-cell genomics identifies cell type-specific molecular changes in autism. *Science, 364*(6441), 685-689. doi:10.1126/science.aav8130

Vogel Ciernia, A., Laufer, B. I., Hwang, H., Dunaway, K. W., Mordaunt, C. E., Coulson, R. L., . . . LaSalle, J. M. (2020). Epigenomic Convergence of Neural-Immune Risk Factors in Neurodevelopmental Disorder Cortex. *Cereb Cortex, 30*(2), 640-655. doi:10.1093/cercor/bhz115

Wamsley, B., Bicks, L., Cheng, Y., Kawaguchi, R., Quintero, D., Margolis, M., . . . Geschwind, D. H. (2024). Molecular cascades and cell type-specific signatures in ASD revealed by single-cell genomics. *Science, 384*(6698), eadh2602. doi:10.1126/science.adh2602

Wong, C. C. Y., Smith, R. G., Hannon, E., Ramaswami, G., Parikshak, N. N., Assary, E., . . . Mill, J. (2019). Genome-wide DNA methylation profiling identifies convergent molecular signatures associated with idiopathic and syndromic autism in post-mortem human brain tissue. *Hum Mol Genet, 28*(13), 2201-2211. doi:10.1093/hmg/ddz052
