## Supplemental Methods for "A Single-Cell Atlas of DNA Methylation in Autism Spectrum Disorder Reveals Distinct Regulatory and Aging Signatures"

### Additional donor characteristics.

Scores from the Autism Diagnostic Interview – Revised (ADI-R) were available for some autistic donors and are summarized below.

| | Median | Range | Donors with information available |
|---|---|---|---|
| ADI-R (social) | 25 | 12 - 36 | 19 |
| ADI-R (communication - verbal) | 17 | 9 - 22 | 14*<br><br>*for 5 donors, only scoring on the nonverbal scale was available |
| ADI-R (communication – non-verbal) | 14 | 9 – 17 | 9 |
| ADI-R (RRB) | 6 | 3 - 10 | 19 |
| ADI-R (36 mos) | 5 | 1 - 5 | 19 |

5 / 26 autistic donors had a history of attention deficit hyperactivity disorder (ADHD), 5 / 26 donors had a history of intellectual and/or developmental delay and 8 donors had history of at least 1 seizure. 1 donor was also diagnosed with bipolar disorder.

### Nuclei isolation from human brain tissue

Small pieces (<50 mg) of tissue were ground in liquid nitrogen with pre-chilled mortar and pestle to generate a homogenous sample and stored at -80°C until use. On the first day of processing, ground tissue was resuspended in 3 mL NIBT:

*Nuclei isolation buffer + triton (NIBT)*

| Component | Final concentration (in mM, unless specified) | Product # |
| --- | --- | --- |
| Sucrose | 250 | various |
| Tris-Cl pH = 8 | 10 | various |
| KCl | 25 | various |
| MgCl2 | 5 | various |
| Triton X-100 | 0.1 % | various |
| DTT | 1 | various |
| Proteinase inhibitor | 1:100 dilution from stock | Sigma-Aldrich #P8340 |
| SUPERaseIn RNase Inhibitor | 0.02U/uL | ThermoFisher Scientific AM2694 |
| RNaseOUT RNase Inhibitor | 0.04U/uL | ThermoFisher Scientific 10777019 |

The lysate was transferred to a pre-chilled 7 mL Dounce homogenizer (Sigma-Aldrich D9063) and dounced using loose and tight pestles for 40 strokes each. The lysate was then mixed with 2 mL of 50% Iodixanol (Sigma-Aldrich D1556) to generate a nuclei suspension with 20% Iodixanol and gently layered on top of a 25% iodixanol cushion. Nuclei were pelleted by centrifugation at 10,000 x g at 4C for 20 min using a swing rotor prior to resuspension in ice-cold DPBS, supplemented with SUPERaseIn RNase Inhibitor and RNaseOUT RNase Inhibitor. For each sample, an aliquot was taken and counted on a Biorad TC20 Automated Cell Counter using trypan blue. The remaining suspension was stained with Hoechst 33342 (2 uM), blocked with 5mg/mL ultrapure BSA (ThermoFisher AM2618) and incubated with an anti-NeuN antibody conjugated to AlexaFluor-488 (clone A60, MilliporeSigma MAB377XMI).

**<u>Bioinformatic Alignment and Quantification.</u>**

We aligned the snmCT-seq data to the GENCODE v40 reference (GRCh38.p13 genome and genic annotation files), building upon the settings described extensively in the flagship manuscript for snmCT-seq [1]. Core changes are the implementation of paired-end alignment and quantification methods, increasing mapping rate and coverage and ensuring that read 1 and read 2 features are not quantified twice when insert size is low. Additionally, we add read deduplication to the RNA reads. Detailed descriptions are available on the pipeline git repository (github.com/chooliu/snmCTseq_Pipeline, release v1.1.0) and summarized below.

We used cell barcodes specific to each reaction well (of 384-well plate) to demultiplex sequence reads into individual nuclei with custom scripts, then trimmed Phred score base calls, Illumina adapter sequences and components specific to snmCT-seq library (the well-barcode, adaptase tail) using fastp v0.23.2 [2] (-f 17 -t 10 -F 15 -T 10 -l 30 --cut_right -q 20 -u 50 -y -Y 15 -x --adapter_fasta).

Methylome reads for each nuclei are then mapped using Bismark v0.24.0 [3] in a two-stage process in order to increase mapping efficiency for PBAT-based reactions and other single-cell methylome preparations [4]: (i) first, reads are mapped in paired-end mode, then (ii) reads unable to map in stage i and singleton reads are mapped in single-end mode. We define singletons as reads without its pair passing trimming quality control. The --maxins 2000 and --pbat (for paired-end and read 1) flags are used, with default settings otherwise. PCR and sequencer optical duplicates (pixel distance 2500) are then removed using Picard MarkDuplicates [5]. Reads with mapping score ≤ 10 were excluded (samtools view -h -q 10; samtools v1.16.1 [6]). A core innovation of the snmCT-seq bench workflow is its simultaneous profiling of RNA and DNA in one reaction well without requiring their physical separation; instead, we perform *in silico* separation of fully methylated RNA libraries introduced by methylated dNTPs (resulting in full methylation, including at CpH- contexts) from the low mCH/CH levels present in libraries originating from genomic DNA. Using the "XM:Z" alignment flag, we then therefore classified the Bismark alignments containing at least three CpH-context cytosines and mCH/CH ≤ 0.50 as DNA reads. Finally, the final resultant DNA reads were used to infer cytosine-level methylation states: for each cytosine *c,* we tabulated the number of alignments suggesting that the cytosine is methylated ($m_c$; C>T bisulfite conversion) versus unmethylated ($u_c$; no conversion) using allcools v1.0.8 [7] (".allc" file, github.com/lhqing/ALLCools).

In parallel, RNA-originating reads were mapped with STAR v2.7.10a [8], also using the two-stage (paired-end, followed by single-end) mapping procedure for RNA quantification. Non-default mapping settings included --alignEndsType EndToEnd --sjdbOverhang 149 --outFilterType BySJout --outFilterMultimapNmax 20 --alignSJoverhangMin 8 --alignSJDBoverhangMin 1 –outFilterMismatchNmax 999 --outFilterMismatchNoverLmax 0.04 --alignIntronMin 20 --alignIntronMax 1000000 --alignMatesGapMax 1000000 as established previously. Alignments that were duplicates identified by Picard or had mapping score ≤ 10 were excluded. To retain RNA reads, each alignment with at least three CpH- context cytosines and mCH/CH ≥0.90 based on the "MD:Z" flag were retained. Finally, RNA counts of each gene were then quantified using featureCounts [9].

We re-validated the performance of the *in silico* separation of RNA and DNA in this paired-end workflow by using three datasets generated with the related, DNA methylation-only assay (snmC-seq2) [10]. Using existing data of this DNA-only assay profiling similar nuclei (BA10 cortex from 25 year old male; Sequence Read Archive SRR6911760, SRR6911772, SRR6911776), we again found high *in silico* discrimination between RNA and DNA based on alignment mCH/CH. When the RNA pipeline (STAR - mCH/CH filtering) was applied to these DNA-only reads, the majority (98.9%) of these bisulfite-converted DNA-only reads failed to align to the genome, and only 0.09% of reads were ultimately incorrectly classified as RNA reads and 0.0089% were quantified as transcript counts using featureCounts. With the DNA pipeline (Bismark - mCH/CH filtering), 0.075% of reads were incorrectly classified as RNA reads and discarded.

**Nuclei-level quality control.**

*Methylome*

We used methylome quality control metrics previously established for brain data[1], including nuclei with a Bismark mapping rate > 50%, at least $1x10^5$ final aligned reads, and global methylation fractions of mCG/CG > 0.50, mCH/CH < 0.20, and mCCC/CCC < 0.03, where the last measure is a proxy for bisulfite conversion rate [1, 7]. Global methylation fraction is defined as $f := \sum_c m_c/(m_c+u_c)$ for all cytosines $c$ from the given cytosine sequence context.

We additionally calculated the relative coverage of chromosome X and Y in each cell's aligned methylome reads to identify potential sample mislabeling or incorrect biobank metadata. Data from 3 female donors was generated but not included in any analyses.

*Transcriptome*

Feature counts obtained for a given cell (across single- and paired-end alignments) were aggregated. All downstream steps were performed in Seurat [11]. Nuclei were included in transcriptomic analysis if they met the following criteria:

- 500 < nGenes / cell < 8,000
- % mitochondrial transcripts < 10
- nCount > 10,000
- % intergenic bases < 10 (output from Picard's RnaSeqMetrics)

These criteria removed almost all nuclei from 1 donor, with low RIN, who was subsequently excluded from RNA analysis.

Genes were included in downstream analysis if they were expressed (count > 0) in at least 10 nuclei.

**Cell Type Clustering and Annotation.**

*Methylome*

We used the allcools pipeline v1.0.8 [7] to cluster nuclei based on their methylome profiles, as described previously [1, 7]. Briefly, we tabulated the CpG- and the CpH-methylation fractions in non-overlapping 100kb genomic bins. Each fraction was normalized to the cell's global methylation fraction levels using a beta-binomial prior. Bins that were autosomal, had less than 20% base overlap with the ENCODE unified exclusion list (encodeproject.org/files/ENCFF356LFX), non-outlying between-cell mean coverage levels (<200, ≥2000), and among the top 25$^{th}$ percentile variable features (allcools calculate_hvf_svr) were used as input features into Principal Components Analysis for dimensionality-reduced features summarizing variability between-nuclei (balanced_pca). The top 54 CpH-PCs and 19 mCpG-PCs (significant_pc_test, p-value < 0.001) were then used as input into UMAP-based methylome-only embedding and an iterative clustering approach in which leiden clustering was repeated 1,000 times with random seeds and an 80% subsample of the nuclei (ConsensusClustering; *k* neighbors = 20, resolution = 1.5), resulting in a tally of 62 methylome-only leiden clusters persistent between subsamples.

*Transcriptome*

Variable feature selection, dimension reduction and clustering were performed using the default Seurat pipeline [11]. Counts were scaled and normalized using the *LogNormalize* method. We derived PCs from 3,000 variable genes and used 20 PCs to find neighbors and in UMAP embedding (leiden resolution = 0.8). We identified 17 clusters, using only gene expression data, and annotated them on the basis of canonical marker gene expression and predicted subclass labels based on data from [12] embedded in Azimuth [13].

*Modality integration*

We integrated transcriptome and methylome data from individual nuclei using a weighted nearest-neighbor approach implemented using methylome and RNA PCs and default parameters in the *FindMultiModalNeighbors* function in Seurat [13]. Dual modality clusters were annotated as described above.

**Transfer of Joint Cluster Annotations to Methylome-Only Nuclei.**

To include nuclei that had pass QC methylomes but not transcriptomes, we then framed the transfer of annotated cluster labels constructed from the joint RNA-methylome WNN embeddings to the methylome-only nuclei as a classification prediction modeling problem with class imbalances (i.e., mitigate biased prediction of rarer cell types like endothelial cells). Thus, we split the nuclei with ground truth joint cluster labels into a training and final test set (2.5% of labeled data; 567 nuclei, cluster-stratified). We mitigated training biases due to class imbalances by applying the Synthetic Minority Oversampling TEchnique algorithm [14] prior to each iteration of model fitting (SMOTE implementation in *themis* R package; 50% oversampling). A grid search across 50 iterations of 5-fold cross-validation was applied to training set to select the model hyperparameters based on mean accuracy, followed by ROC-AUC, PR-AUC. Ultimately, we predicted cluster labels (model outcome) from the 100kb-mCH and 100kb-mCG principal components (model predictors) using the weighted k-nearest neighbors algorithm [15] (L1-distance, kernel = “optimal”, k = 35; implementation from the *kknn* v1.3.1 and *tidymodels* v1.1.1 R packages) due to overall accuracy plateauing at this value of *k*. We further checked performance within each class- and subclass-, observing the lowest accuracy for the “Unannotated” nuclei class and observing mean cross-validation accuracy ≥94% for all subclasses.

In the final hold-out test set, accuracy was 98.7% at the subclass annotation level and 97.4% at the cluster-level (15/576 hold-out nuclei with predicted labels different than ground truth label). The majority of misclassifications were observed only between highly related clusters (e.g., “Inh CGE VIP 1” incorrectly predicted to be “Inh CGE VIP 2”) or due to the model erring on labeling nuclei as “Unannotated.” However, the former cluster-level annotation is a finer resolution than the primary “subclass” level of methylation analysis used throughout the text and thus has no primary analytical consequences, and the latter misclassification errs on being conservative with methylomes included in the final pseudobulk.

After applying the final model to methylome-only nuclei, we excluded nuclei that either had “Unannotated” labels or were members of methylation-only based leiden clusters with majority “Unannotated” labels (e.g., excluding methylation-only leiden-consensus-cluster “c26” had majority “Unannotated”), resulting in the final nuclei count of 65,023 single-nucleus methylomes.

**Pseudobulk DEG Calling.**

In our primary analysis, data were aggregated to pseudobulk and differential gene expression analysis was performed using DESeq2 [16]; both steps were implemented using muscat [17].

Clusters, donors and features were filtered using the following criteria, before pseudobulk aggregation:

- Tested clusters with greater than 500 total nuclei
- Generated pseudobulk count matrix from donors with at least 10 nuclei in a given cell type
- Tested genes expressed (count > 0) in 20% of nuclei within a cluster

We estimated and controlled for variables independent of the diagnosis effect using RUVseq [18] (using the residual-based method, RUVr; k = 5).

To perform the downsampling analysis in *SFig. 6e*, we randomly sampled 20 cells per donor per cell type across 500 permutations, repeated the procedure described above and reported the number of DEGs (padj < 0.1).

To assess whether we could recapitulate previously published results, we implemented the same type of differential expression models as the reference paper.

To compare DEGs with Velmeshev et al. [19] in *SFig. 6f*, we re-analyzed our data using MAST [20] and a linear mixed model similar to Velmeshev et al. [19] :

```
zlmDx <- zlm(formula = ~Diagnosis + (1|Donor) + nFeature_RNA + Age + RIN + PMI +
(1|batch) + percent.mt + BrainBank,
        sca = sca,
        method = "glmer",
        ebayes = F,
        silent=F)
```

To call age-DEGs in *SFig. 13c*, we used a model similar to Chien et al. [21], including only age and diagnosis in the analysis. Diagnosis effects (quantified as gene-wise logFC were very similar in this reduced model compared to the full model described above and are reported in *SFile 3*).

### GSEA/GO

*Transcriptome*

We performed gene set enrichment analysis of differentially expressed genes using the *fgseaMultilevel* function of fgsea [22] and the biological process (BP) and molecular function (MF) subsets of the C5 ontology gene sets available here: https://www.gsea-msigdb.org/gsea/msigdb/collections.jsp (download July 2024). Input lists were ordered by effect size (log FC) * -log10(pvalue) and we considered gene sets between 25 and 500 genes in size.

*Methylome*

We assessed gene ontology (GO) enrichment analysis on DMRs annotated to transcriptional start sites using gprofiler2 [23]. We observed an association between genic DMRs (in transcriptional start and termination sites, exons and introns) and gene length. To adjust for this, we performed GO analysis of genic DMRs using goseq [24].

**Whole Genome Sequencing and Genotyping.** Genomic DNA was extracted using Zymo's Quick-DNA mini-prep plus kit (#4068) and libraries were prepared using Roche's KAPA Hyper kit (#07962363001) and sequenced on a Novaseq6000 (in 2x100bp and 2x150bp configurations). The resulting reads were processed using the GATK4 [25] best practices workflow for short variant germline discovery (SNPs and indels; accessed in January 2023). The reads were aligned using bwa-mem v123. The alignments were deduplicated (gatk MarkDuplicates) then quality-corrected on a per-sample, -run, and -lane read group basis using Base Score Quality Recalibration (GATK ApplyBQSR v4.1.0) with reference to four databases of known variants (Homo_sapiens_assembly38.dbsnp138.vcf, hg38_v0_1000G_phase3_v4_20130502.sites.hg38.vcf, hapmap_3.3.hg38.vcf, Mills_and_1000G_gold_standard.indels.hg38.vcf; via GATK Google Cloud bucket). SNPs and indels were called per-sample using GATK HaplotypeCaller, then hard-filtered using the best practices recommendations for SNPs (QD < 2, QUAL < 30, SOR > 3, FS > 60, MQ < 40, MQRankSum < -12.5, ReadPosRankSum < -8) and indels (QD < 2, QUAL < 30, FS > 200, ReadPosRankSum < -20, SOR > 10). The final mean autosomal alignment depths were 40.9X ± 10.3 (across-sample mean and sd), with no significant mean difference in depth or other quality metrics in by diagnosis or age in years (t-test p-value > 0.59).

**Ancestry Inference and Correction.**

We then joined the case-control WGS genotypes to the 1000 Genomes reference panel (Byrska-Bishop [26] variant-filtered and phased release; internationalgenome.org/data-portal/data-collection/30x-grch38), subset to unrelated individuals. Using plink2 v2.00a3.7 [27] and bcftools v1.16 [6], we removed variants that were lower frequency (MAF < 0.05) or overlapped previously reported regions of long-range LD (e.g., HLA region; regions extracted from Price, et al. (2008) [28] and converted to GRCh38 coordinates using the liftOver R/Bioconductor package v1.18.0 with UCSC hg19ToHg38.over.chain.gz), followed by LD pruning (plink2 --indep-pairwise 1000 100 0.25). We ran principal component analysis on the resulting joined genotypes (flashPCA v2.0 [29]) and retained the first two PCs for modeling. Continental ancestry was assigned to each donor using the weighted k-nearest neighbor algorithm [15] calculated on the first ten PCs (*kknn*, *tidymodels* implementations; hyperparameters k = 18, distance kernel = 1, power = "biweight" selected to optimize multiclass-accuracy and PR-AUC via 10-fold cross-validation on the 1000 Genomes dataset, with balanced sampling of each superpopulation via SMOTE [14]). In parallel, the joined case-control and 1000 Genomes genotypes in ADMIXTURE v1.30.0 [30] with $k$ = 5 clusters, for one cluster per 1000 Genomes superpopulation. We assigned each donor to the continental ancestry with the majority admixture fraction. Both PCA and ADMIXTURE continental ancestry predictions were also consistent with biobank records of donor Self-Identified Race/Ethnicity, where available. We again checked the sex of the samples using chrX heterozygosity (plink2 --check-sex), which were again consistent with exclusion of n=3 female samples.

**Genotype Masking of the Methylome.**

Donor genetic deviation from the reference genome introduces noise into methylation quantification: for example, alignments over a loss-of-cytosine C>T mutation would incorrectly be interpreted as a bisulfite-converted demethylated cytosine, when it might more appropriately be quantified as "not applicable" for the given donor. Further, a change of dinucleotide context strongly changes the biophysical/enzymatic mechanisms of methylation deposition and loss, with CpG- methylation having higher methylation levels compared to CpH- methylation, where H is the IUPAC code for A, C, or T. We thus set the methylation counts and coverage of a given donor's cytosines to zero if the cytosine overlapped a (1) loss-of-cytosine, a (2) change in the cytosine's +1 context from CG>CH or CH>CG, or (3) an indel. These "genotype-masked" calls were subsequently used prior to any differential methylation testing by diagnosis, age, or cell-type.

**Differential Methylation Identification Strategies.**

We aggregated genotype-masked pseudobulks at the subclass-by-donor level, then called DMRs within each subclass using two complementary methods: first, we compared mean methylation levels by diagnosis over (1) pre-defined genomic features including genes, promoters, and genomic sliding windows, then compared (2) *de novo* regions based on DSS, a sensitive DMR caller that tests individual sites and exploits correlation between adjacent cytosines. These two approaches have different strengths. Notably, the former likely has the benefit of increased coverage and by extension robustness, but at the critical cost of sensitivity (e.g., if between-diagnosis methylation differences occur only in a portion of a gene or ≤500bp small interval). In addition, the former also enabled testing of non-CG methylation and more complex covariate adjustment, including non-linear age trends and age-by-diagnosis interactions.

***De Novo* DMRs.**

We then modeled the $m_c$ and $u_c$ counts at single CpG-context cytosines using the *DSS* v2.50.1 R package [32], which similarly applies a beta-binomial model, but tests for a significant mean difference between two categorical groups at individual cytosines, and applies a shrinkage adjustment that incorporates spatial correlation in methylation levels between adjacent CpG-context cytosines. This local smoothing adjustment may lend to DSS's high sensitivity, with benchmarks by independent DMR methods developers finding DSS to have the highest power among *de novo* DMR callers at <10X genomic count levels [33]. After removing cytosines overlapping the ENCODE Unified Exclusion list or had zero coverage in most samples (zero in ≥75% of samples), we ran DSS with the default prior and a 500bp smoothing window. To err on being conservative with reported results, we further elected to report significant regions instead of single cytosines (*de novo* regions called "DMRs" throughout the text for short). We thus aggregate the single cytosine results into larger candidate regions by aggregating sites within ±250bp, then retaining regions with at ≥2 multiple testing adjusted significant cytosines (Storey q-value < 0.05) and at least one significant cytosine with methylation fraction (*f)* estimated effect size $|\Delta| > 0.10$.

For the ASD-DMRs, which we expected to have smaller effect sizes, we additionally required a stringent criterion of low colocalization with permuted DMRs (POF < 0.20). We define the POF = Permutation Overlap Fraction := [# of overlaps with permuted DMRs] / [# permutations] for a given candidate region. The tabulation entailed a rigorous procedure in which the diagnosis labels corresponding to each donor were shuffled then the full DMR calling procedure was re-run (DSS, aggregation into candidate regions) (**Fig 2A**). We performed 500 permutations within each subclass, then tabulated how many times the observed candidate DMRs intersect with the permuted DMRs. Candidate ASD-DMRs with low POF represent regions that were unlikely to be observed under the permuted null distribution. We repeated this process including the stringent permutation for all-donor ASD-DMRs, as well as age-stratified subsets of the data: "young" (2-16), "mid" (17-30), and "older" (40-60) samples within a given subclass as a method to control for age effects. The threshold delineating young and mid for age-stratified ASD-DMRs were selected to approximately maintain the same sample sizes in each group (n=19, n=20, n=13).

Using this DSS to DMR procedure, we additionally called age-DMRs within each subclass by pairwise testing between the young, mid, and older age strata. Unless otherwise noted, we primarily report and analyze the union of the three pairwise tests within each subclass. To call cell-type DMRs, we repeated the DMR calling procedure between each pairwise comparison of the eight subclasses (e.g., Astro versus Exc L2/3, Astro versus Inh MGE, etc.). A cell-type DMR for subclass *s* are identified as hypo-methylated in cell-type *s* compared to ≥5 other subclasses.

**DMR and DMB Enrichment Testing.**

To assess the enrichment of DMRs and DMBs with external annotation datasets, we used a permutation-based approach that compared the number of overlaps in the observed DMRs and DMBs with the external dataset with that of 500 - 1 million random background sets (a subset of which were length matched). Each background was respectively extracted from the resampleRegions and randomizeRegions functions of regioneR and regioneReloaded v1.4.0 [34, 35], restricted to the candidate regions (i.e., at least two CG sites tested in DSS in ±250bp span, but without the Δ and significance criteria described above) or candidate bins (tested in model; $f_r > 0.05$, ≥2 counts in 75% of donors). Empirical p-values were generated by comparing the observed overlap counts to the distribution of background overlaps: p = (1 + number of permutations where |x_i - mean_permuted| ≥ |x_obs - mean_permuted|) / (1 + N) (Granges v1.54.1 [36], findOverlaps). The permutation based p-values for each covariate-annotation set (where covariates were ASD-, Age-, or Interaction-) were corrected for multiple testing (Benjamini-Hochberg [37] FDR < 0.05).

Sets of annotations included the following:

- Chromatin states (Roadmap Epigenomics [38] 15-state ChromHMM model from egg2.wustl.edu/roadmap/web_portal/chr_state_learning.html; adult PFC “E073” and fetal brain “E081” samples)
- Transposable elements and repeats (RepeatMasker [39], as obtained from genome.ucsc.edu/cgi-bin/hgTrackUi?g=rmsk and omitting ambiguous classifications marked with “?”)
- GENCODE v40 .gtf genic features, based on the following priority: promoter > 5′ UTR > 3’ UTR > exon > intron > 10kb upstream of protein-coding gene > 10kb downstream protein-coding gene > non-protein-coding gene > 10kb upstream of non-protein-coding gene > 10kb downstream of non-protein-coding gene > distal intergenic (not in any of above categories).
- Bulk ASD-DMR studies: External DMRs were obtained from bulk postmortem cortical studies of idiopathic ASD (iASD), Rett syndrome (RTT), and Dup15q syndrome. These included an Illumina 450k array dataset from Wong et al. (n = 223 across PFC, TC, and CB; hg19), from which cross-cortex iASD and Dup15q DMPs were used [40]. Coordinates were lifted over to hg38 using sesameData (v1.20.0) [41] and extended ±250 bp to approximate regional methylation. Wong et al. also reported cross-cortex Dup15q DMRs which were included here, although these were aggregated without regard to direction. Additional iASD-, RTT-, and Dup15q-associated DMRs were pulled from a whole-genome bisulfite sequencing (WGBS) dataset from Ciernia et al. (n = 49, BA9; hg38) [42].
- H3K27Ac fetal brain peaks were downloaded from [43]
- Open chromatin fetal brain (consensus ATACseq peaks) data was downloaded from [44]

- H3K27Ac peaks (DER-05_PFC_H3K27ac_peaks) and high confidence enhancers (DER-04b_hg38lft_high_confidence_PEC_enhancers) in adult brain were available from the PsychEncode I package (http://resource.psychencode.org/). ,
- Cell type specific ATACseq peaks (candidate cis regulatory regions) were downloaded from [45]. We then generated individual files aggregating all cCRE IDs associated with each cell type.

### Methylome Percent Variance Explained.

To assess the overall methylome variance explained by each known covariate, we extracted methylation corresponding to different features: genes, promoters, 100kb bins and 50kb bins. For the bins, we also considered a "-genes" version excluding bins with ≥0.20 fractional overlap with protein coding genes. For each feature set and for CpA-, CpG-, and CpH- methylation, we used the allcools pipeline described previously to calculate global methylation level level-normalized methylation fractions across features and extracted the top ten PCs. We then ran PERMANOVA (vegan v2.6-8 [46] with 10,000 permutations, Euclidean distance) on the top *k* PCs to extract the variance explained $R^2$ and significance of the association (permutation F-test). The value of *k* was selected using findPC v1.0 [47], using the median cross-method 'elbow' suggested, with a minimum of k = 2. We note that we elected to normalize for pseudobulk-level global mC levels; this results in non-zero percent variances of because global methylation levels vary with age, the normalized values may underestimate the proportion of variance explained by age in some cell types.

### Transcription Factor Motif Enrichment.

DMR motif enrichment analysis was performed using the *findMotifsGenome* function in HOMER [48] and the fasta file corresponding to the same genome used for alignment.

### Cross-Modality Correlation Analyses.

**Epigenetic Age Clocks.**

The probe lists were extracted from the 353-probe and 111-probe Horvath age clocks [53] and the cortex age clock [54], then converted to their corresponding GRCh38 single-cytosine coordinates using sesameData v1.20.0 [41]. We extracted a ±1bp region in each of the donor-by-subclass methylome pseudobulk to calculate the corresponding methylation fractions as a proxy for $f_p$ for probe $p$ (often "β" in array literature) at each probe, with the ±1bp expansion due to the high cross-strand symmetry and correlation of CpG-methylation. The epigenetic age estimate was then calculated for each of the pseudobulks, per usual clock calculations $\hat{}Age = g(\sum_p w_p f_p)$ where $w_p$ are the probe weights (LASSO coefficients) and where $g(x)$ is the calibration function shared by both clocks to account for the faster epigenetic changes reported before adult maturation[33,34]: $g(x) = (1+ \tau)\exp(x)-1$ for $x < 0$ and $(1 + \tau)*x + \tau$ for $x \geq 0$ for $\tau = 20$.

We evaluated evidence of cell-type specific epigenetic aging differences between ASD and CTL by evaluating whether the residual epigenetic age ($\hat{}Age - \hat{}Age_{loess}$) differed by group with a t-test, where the loess mean trend ($\hat{}Age_{loess}$, span=1) between the true age and epigenetic age was used as a "calibration curve" correcting for prediction errors between the clock age and chronological age specific to our dataset. We also evaluate the performance of each clock in predicting the biobank ground truth donor age by tabulating the Mean Absolute Error (MAE) and Spearman correlation between the predicted and ground truth ages.

We performed three additional analyses to assess the robustness of our findings to coverage. First, we constructed a high coverage "pseudo-tissue" by calculating a given donor's methylation fraction from all eight major subclasses together; as each clock is validated in bulk data, this composite thus represents a performance bound due to technical effects of transferring the array-based technology to our sequencing-based measurements. Second, we hypothesized that given the spatial correlation of CG-methylation—particularly with the high proportion of clock probes overlapping either a promoter or CpG-island (55% in cortex clock, 81% in Horvath-353)—that padding fractional methylation calculations to include sites within a ±*w* base window of each probe may reduce noise. We thus also evaluated the impact of padding the *f* calculation with all cytosines in a ±25bp, ±50bp, or ±100bp range. Third, we calculated the mean across-pseudobulk coverage at each probe site and repeated all epigenetic age prediction when each pseudobulk and pseudo-tissue were sampled with replacement to this mean coverage level. This coverage resampling was performed 100 times.

### References


1. Luo, C., et al., *Single nucleus multi-omics identifies human cortical cell regulatory genome diversity.* Cell Genom, 2022. **2**(3).
2. Chen, S., et al., *fastp: an ultra-fast all-in-one FASTQ preprocessor.* Bioinformatics, 2018. **34**(17): p. i884-i890.
3. Krueger, F. and S.R. Andrews, *Bismark: a flexible aligner and methylation caller for Bisulfite-Seq applications.* Bioinformatics, 2011. **27**(11): p. 1571-2.
4. Krueger, F. *Single cell PBAT.* 2019; Available from: https://felixkrueger.github.io/Bismark/faq/single_cell_pbat/.
5. Institute, B. *Picard Tools*. Available from: http://broadinstitute.github.io/picard/
6. Danecek, P., et al., *Twelve years of SAMtools and BCFtools.* Gigascience, 2021. **10**(2).
7. Liu, H., et al., *DNA methylation atlas of the mouse brain at single-cell resolution.* Nature, 2021. **598**(7879): p. 120-128.
8. Dobin, A., et al., *STAR: ultrafast universal RNA-seq aligner.* Bioinformatics, 2013. **29**(1): p. 15-21.
9. Liao, Y., G.K. Smyth, and W. Shi, *featureCounts: an efficient general purpose program for assigning sequence reads to genomic features.* Bioinformatics, 2014. **30**(7): p. 923-30.
10. Luo, C., et al., *Robust single-cell DNA methylome profiling with snmC-seq2.* Nat Commun, 2018. **9**(1): p. 3824.
11. Butler, A., et al., *Integrating single-cell transcriptomic data across different conditions, technologies, and species.* Nat Biotechnol, 2018. **36**(5): p. 411-420.
12. Bakken, T.E., et al., *Comparative cellular analysis of motor cortex in human, marmoset and mouse.* Nature, 2021. **598**(7879): p. 111-119.
13. Hao, Y., et al., *Integrated analysis of multimodal single-cell data.* Cell, 2021. **184**(13): p. 3573-3587 e29.
14. Chawla, N.V., et al., *SMOTE: Synthetic Minority Over-sampling Technique.* Journal of Artificial Intelligence Research, 2002. **16**: p. 321-357.
15. Schliep, K. and K. Hechenbichler *Weighted k-Nearest-Neighbor Techniques and Ordinal Classification*. 2004. **399**,  DOI: 10.5282/ubm/epub.1769.
16. Love, M.I., W. Huber, and S. Anders, *Moderated estimation of fold change and dispersion for RNA-seq data with DESeq2.* Genome Biol, 2014. **15**(12): p. 550.
17. Crowell, H.L., et al., *muscat detects subpopulation-specific state transitions from multi-sample multi-condition single-cell transcriptomics data.* Nat Commun, 2020. **11**(1): p. 6077.

18. Risso, D., et al., *Normalization of RNA-seq data using factor analysis of control genes or samples.* Nat Biotechnol, 2014. **32**(9): p. 896-902.
19. Velmeshev, D., et al., *Single-cell genomics identifies cell type-specific molecular changes in autism.* Science, 2019. **364**(6441): p. 685-689.
20. Finak, G., et al., *MAST: a flexible statistical framework for assessing transcriptional changes and characterizing heterogeneity in single-cell RNA sequencing data.* Genome Biol, 2015. **16**: p. 278.
21. Chien, J.F., et al., *Cell-type-specific effects of age and sex on human cortical neurons.* Neuron, 2024. **112**(15): p. 2524-2539 e5.
22. Korotkevich, G., et al., *Fast gene set enrichment analysis.* bioRxiv, 2021.
23. Kolberg, L., et al., *gprofiler2 -- an R package for gene list functional enrichment analysis and namespace conversion toolset g:Profiler.* F1000Res, 2020. **9**.
24. Young, M.D., et al., *Gene ontology analysis for RNA-seq: accounting for selection bias.* Genome Biol, 2010. **11**(2): p. R14.
25. Van der Auwera, G.A., et al., *From FastQ data to high confidence variant calls: the Genome Analysis Toolkit best practices pipeline.* Curr Protoc Bioinformatics, 2013. **43**(1110): p. 11 10 1-11 10 33.
26. Byrska-Bishop, M., et al., *High-coverage whole-genome sequencing of the expanded 1000 Genomes Project cohort including 602 trios.* Cell, 2022. **185**(18): p. 3426-3440 e19.
27. Chang, C.C., et al., *Second-generation PLINK: rising to the challenge of larger and richer datasets.* Gigascience, 2015. **4**: p. 7.
28. Price, A.L., et al., *Long-range LD can confound genome scans in admixed populations.* Am J Hum Genet, 2008. **83**(1): p. 132-5; author reply 135-9.
29. Abraham, G., Y. Qiu, and M. Inouye, *FlashPCA2: principal component analysis of Biobank-scale genotype datasets.* Bioinformatics, 2017. **33**(17): p. 2776-2778.
30. Alexander, D.H., J. Novembre, and K. Lange, *Fast model-based estimation of ancestry in unrelated individuals.* Genome Res, 2009. **19**(9): p. 1655-64.
31. Storey, J.D. and R. Tibshirani, *Statistical significance for genomewide studies.* Proc Natl Acad Sci U S A, 2003. **100**(16): p. 9440-5.
32. Feng, H. and H. Wu, *Differential methylation analysis for bisulfite sequencing using DSS.* Quant Biol, 2019. **7**(4): p. 327-334.
33. Peters, T.J., et al., *Calling differentially methylated regions from whole genome bisulphite sequencing with DMRcate.* Nucleic Acids Res, 2021. **49**(19): p. e109.
34. Gel, B., et al., *regioneR: an R/Bioconductor package for the association analysis of genomic regions based on permutation tests.* Bioinformatics, 2016. **32**(2): p. 289-91.
35. Malinverni, R., et al., *regioneReloaded: evaluating the association of multiple genomic region sets.* Bioinformatics, 2023. **39**(11).
36. Lawrence, M., et al., *Software for computing and annotating genomic ranges.* PLoS Comput Biol, 2013. **9**(8): p. e1003118.
37. Benjamini, Y. and Y. Hochberg, *Controlling the False Discovery Rate: A Practical and Powerful Approach to Multiple Testing.* Journal of the Royal Statistical Society Series B: Statistical Methodology, 1995. **57**(1): p. 289-300.
38. Roadmap Epigenomics, C., et al., *Integrative analysis of 111 reference human epigenomes.* Nature, 2015. **518**(7539): p. 317-30.
39. Smit, A., R. Hubley, and P. Green. *RepeatMasker Open-4.0.* 2013 - 2015; Available from: http://www.repeatmasker.org.

40. Wong, C.C.Y., et al., *Genome-wide DNA methylation profiling identifies convergent molecular signatures associated with idiopathic and syndromic autism in post-mortem human brain tissue.* Hum Mol Genet, 2019. **28**(13): p. 2201-2211.
41. Zhou, W., et al., *SeSAMe: reducing artifactual detection of DNA methylation by Infinium BeadChips in genomic deletions.* Nucleic Acids Res, 2018. **46**(20): p. e123.
42. Vogel Ciernia, A., et al., *Epigenomic Convergence of Neural-Immune Risk Factors in Neurodevelopmental Disorder Cortex.* Cereb Cortex, 2020. **30**(2): p. 640-655.
43. Li, M., et al., *Integrative functional genomic analysis of human brain development and neuropsychiatric risks.* Science, 2018. **362**(6420).
44. Trevino, A.E., et al., *Chromatin and gene-regulatory dynamics of the developing human cerebral cortex at single-cell resolution.* Cell, 2021. **184**(19): p. 5053-5069 e23.
45. Li, Y.E., et al., *A comparative atlas of single-cell chromatin accessibility in the human brain.* Science, 2023. **382**(6667): p. eadf7044.
46. Oksanen J, et al. *vegan: Community Ecology Package*. 2025; Available from: https://vegandevs.github.io/vegan/.
47. Zhuang, H., H. Wang, and Z. Ji, *findPC: An R package to automatically select the number of principal components in single-cell analysis.* Bioinformatics, 2022. **38**(10): p. 2949-2951.
48. Heinz, S., et al., *Simple combinations of lineage-determining transcription factors prime cis-regulatory elements required for macrophage and B cell identities.* Mol Cell, 2010. **38**(4): p. 576-89.
49. Liu, W., et al., *Smccnet 2.0: a comprehensive tool for multi-omics network inference with shiny visualization.* BMC Bioinformatics, 2024. **25**(1): p. 276.
50. Kuleshov, M.V., et al., *Enrichr: a comprehensive gene set enrichment analysis web server 2016 update.* Nucleic Acids Res, 2016. **44**(W1): p. W90-7.
51. Gene Ontology, C., et al., *The Gene Ontology knowledgebase in 2023.* Genetics, 2023. **224**(1).
52. Sarda-Espinosa, A. *dtwclust: Time Series Clustering Along with Optimizations for the Dynamic Time Warping Distance*. 2024; Available from: https://cran.r-project.org/package=dtwclust.
53. Horvath, S., *DNA methylation age of human tissues and cell types.* Genome Biol, 2013. **14**(10): p. R115.
54. Shireby, G.L., et al., *Recalibrating the epigenetic clock: implications for assessing biological age in the human cortex.* Brain, 2020. **143**(12): p. 3763-3775.
